# Climatic niche pre-adaptation in mainland Europe facilitated the colonization of Madeira by ivies (*Hedera* L., Araliaceae)

**DOI:** 10.1101/2021.07.16.452604

**Authors:** Alejandro Alonso, Angélica Gallego-Narbón, Marina Coca de la Iglesia, David Monjas, Nagore G. Medina, Mario Fernández-Mazuecos, Virginia Valcárcel

## Abstract

**Background and aims:** The way plants cope with biotic and abiotic selective pressures determines their success in the colonization of remote oceanic islands, which ultimately depends on the phylogenetic constrains and ecological response of the lineage. In this study we aim to evaluate the relative role of geographical and ecological forces in the origin and evolution of the Madeiran ivy (*H. maderensis*).

**Methods:** To determine the phylogenetic placement of *H. maderensis* within the western polyploid clade of *Hedera* (three species), we analysed 40 populations (92 individuals) using genotyping-by-sequencing and including *H. helix* as outgroup. Climatic niche differences among the four study species were evaluated using a database with 706 records representing the entire species ranges. To test species responses to climate, a set of 19 vegetative and regenerative functional traits were examined for 70 populations (335 individuals).

**Key results:** Phylogenomic results revealed a nested pattern with *H. maderensis* embedded within *H. iberica*. Gradual niche differentiation from the coldest and most continental populations of *H. iberica* to the warm and stable coastal population sister to *H. maderensis* parallels the geographical pattern observed in the phylogeny. Similarity in adaptive traits is observed for *H. maderensis* and *H. iberica*. The two species show leaves with higher SLA, lower LDMC and thickness and smaller fruits than those of *H. hibernica*.

**Conclusions:** Acquisition of the Macaronesian climatic niche and the associated functional syndrome in mainland European ivies (small fruits, leaves high SLA, and low LDMD and thickness) was a key step in the colonization of Madeira by the *H. iberica/H. maderensis* lineage, which points to climatic pre-adaptation as a driver of island colonization (dispersal and establishment). Once in Madeira, speciation was driven by geographical isolation, while ecological processes are regarded as secondary forces with a putative impact in the lack of further in situ diversification.

## INTRODUCTION

The ways in which species disperse to oceanic islands, and then potentially establish and speciate on them, remain challenging open questions in biogeography (Patiño *et al*. 2017). Achieving island colonization (including dispersal and establishment) involves the successful interaction between ecological and evolutionary processes (Alzate *et al*. 2020). The dispersal filter depends on the propagule pressure (frequency of dispersal events and number of propagules per dispersal event), which is mainly determined by the traits of the propagules and the distance from the source to the recipient island. The environmental filter depends more on the biotic and abiotic pressures during dispersal and mostly upon arrival, and it is related to pre-adaptation or easiness to adapt to new environments. The course of evolution after island colonization (non-speciation, speciation without diversification, speciation with diversification or radiation) depends on the time elapsed since colonization, the degree of isolation from the mainland, the geographical and ecological opportunities for speciation (biotic and abiotic pressures), the propagule pressure and the phylogenetic constraints of the lineage (Stuessy *et al*. 2006; Takayama *et al*. 2015).

Macaronesia is a biogeographic region including four major oceanic archipelagos (Azores, Madeira, Canary Islands and Cape Verde) and 42 main islands in the northeast Atlantic Ocean. As many other oceanic archipelagos, these are characterized by high levels of plant endemism that combine spectacular radiations with a surprisingly high proportion of endemics that have not further diversified (single-species endemics; Patiño *et al*. 2014). However, the proximity of most Macaronesian archipelagos to the mainland makes their patterns of colonization and speciation unusual in the context of the world’s oceanic archipelagos (Kim *et al*. 2008) and these patterns have deserved much attention from naturalists and scientists since Humdbolt’s visit to Tenerife in 1799 (Gebauer 2014). The archipelago of Madeira includes three major emerged islands (Madeira, Porto Santo, and Deserta Grande; 32 - 33° N, 16° - 17° W) that, together with several additional islands and islets and five sunken palaeo-islands, constitute the Madeiran Volcanic Province (Fernández-Palacios *et al*. 2011). Madeira, the largest island of the archipelago, is moderately small in comparison to other Macaronesian islands (c. 750 km^2^), and its emergence date (5.6 Ma; Geldmacher *et al*. 2000) is relatively recent considering the long history of the Madeiran Volcanic Province and of Macaronesia as a whole (Fernández-Palacios *et al*. 2011). Despite its comparatively small area, Madeira is the second richest island of Macaronesia in terms of vascular plants (1,136 species, including 94 endemics; Borges *et al*. 2008), only surpassed by Tenerife (Canary Islands).

The genus *Hedera* includes 12 liana species (ivies, 13 taxa; Valcárcel et al. 2017a; Valcárcel & Vargas, 2010) and naturally occurs in three continents (Europe, Asia and Africa), as well as in Macaronesia. The ability of ivies to disperse over long distances and occupy a wide range of environments regarding sun exposure, water availability, soil preferences and altitude (Kollman *et al*. 1999; Sack *et al*. 2000; author’s *pers. obs*.) may be partly responsible for the broad distribution range of the genus. Yet, half of the ivy taxa (seven) are regional endemics, most of which (five) occur on islands (Valcárcel 2008). In fact, the complex evolutionary history of ivies includes three independent colonization events of three different Macaronesian archipelagos (Vargas *et al*. 1999). Each colonization event has produced a single-endemic species, with no further in situ diversifications. Thus, endemic Macaronesian ivies are interesting study systems for analysing the origin of single-species endemics. One of these island endemics is the Madeiran ivy (*Hedera maderensis* K. Koch ex Rutherford) a frequent element in the laurel forests of the island of Madeira (Capelo *et al*. 2005).

Previous phylogenetic studies in *Hedera* based on a small number of DNA regions (Vargas *et al*. 1999; Grivet and Petit 2002; Ackerfield and Wen 2003; Valcárcel *et al*. 2003; Green *et al*. 2011) pointed to a complex evolutionary history involving (i) multiple hybridization events coupled with polyploidizations, which affected almost half of the species (Vargas *et al*. 1999; Valcárcel *et al*. 2003; Escudero *et al*. 2014a), (ii) recurrent periods of range expansion and retraction in parallel with major extinctions and speciation events at the regional scale (Valcárcel *et al*. 2017b), and (iii) an old evolutionary history for the genus despite recent diversification of extant species (Valcárcel *et al*. 2014). The limited power of traditional Sanger sequencing to provide enough DNA sequence variability to resolve phylogenetic relationships in such complex scenarios is compounded by the fact that ivies, and the whole Araliaceae family, have a slow nucleotide substitution rate (Valcárcel and Wen 2019). Indeed, the limited variability of the DNA regions analysed so far is responsible for the low resolution and large polytomies in *Hedera* phylogenies (Vargas *et al*. 1999; Ackerfield and Wen 2003; Valcárcel *et al*. 2003, 2017b; Green *et al*. 2011) that have prevented clarification of monophyly and sister-group relationships of several species (Valcárcel and Vargas 2010). This is the case of *H. maderensis*, whose geographic origin and phylogenetic affinities remain uncertain (Vargas *et al*. 1999; Valcárcel *et al*. 2003, 2017b). According to reconstructions based on nuclear DNA, the Madeiran ivy is part of a polytomy in the western polyploid clade of *Hedera*, together with *H. iberica* and *H. hibernica* (G. Kirchner) Bean (Vargas *et al*. 1999; Valcárcel *et al*. 2003). While *H. maderensis* and *H. iberica* are hexaploid endemic species with relatively restricted distributions in Madeira and SW Iberian Peninsula (Fig. 1), *H. hibernica* is a tetraploid species widely distributed across the western faces of the British Isles, France and northern half of the Iberian Peninsula (Fig. 1).

**Figure 1.**
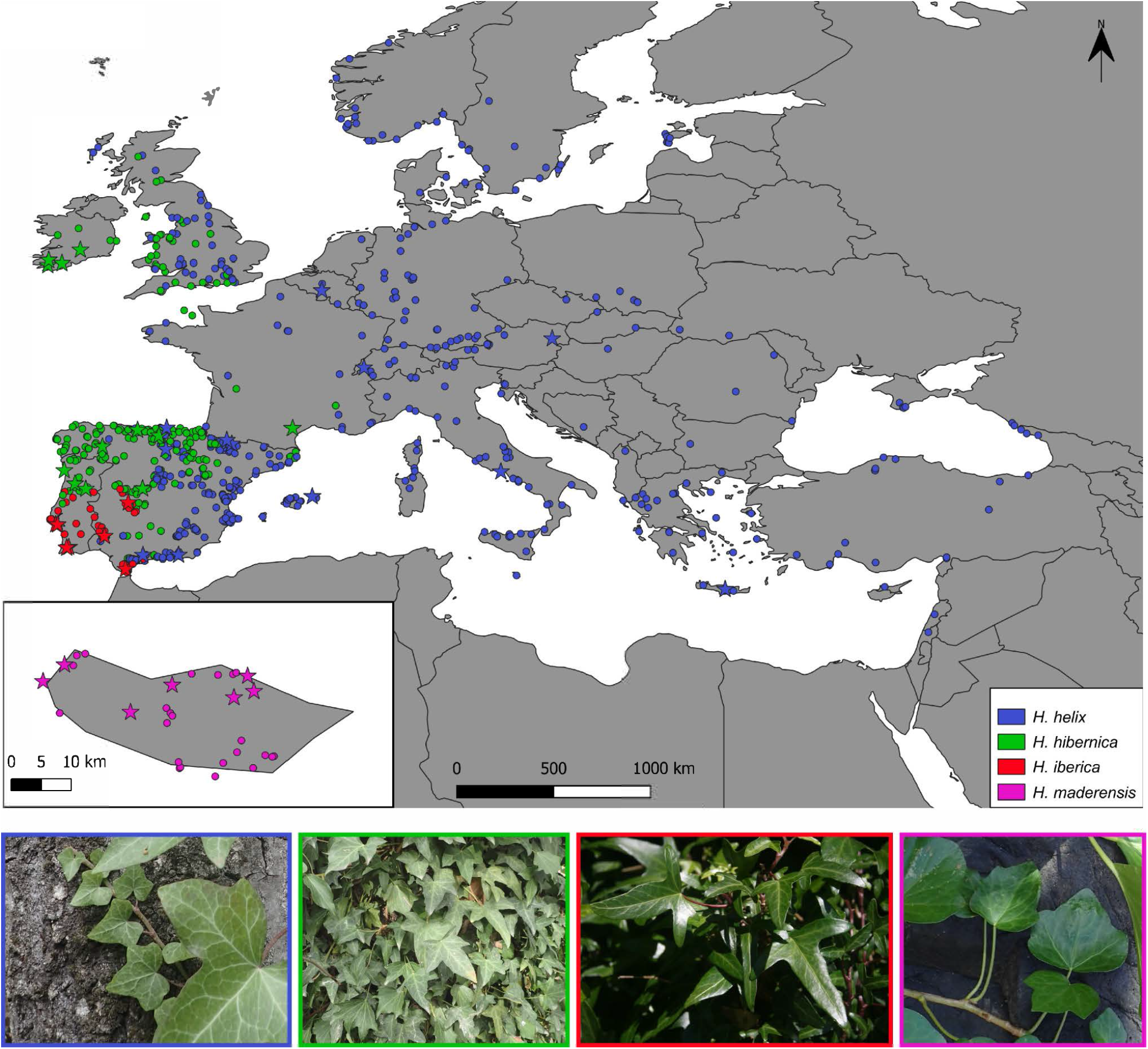
Occurrence-based map of the distribution ranges of the three species of the western polyploidy clade of *Hedera* (*H. hibernica, H. iberica, H. maderensis*) and *H. helix*. Circles indicate samples included for the climatic niche study. Stars indicate samples also included in the phylogenetic study.

Based on cytogenetic and geographical patterns, speciation in the western polyploid clade of *Hedera* has been traditionally attributed to the interplay between polyploidization and geographical barriers (Vargas *et al*. 1999; Valcárcel *et al*. 2003; Escudero *et al*. 2014a). On the one hand, polyploidization has been considered the main driving force of speciation in the mainland, as shown by the tetraploid *H. hibernica* and the hexaploid *H. iberica*, which come into contact in the central-western Iberian Peninsula (Fig. 1). On the other hand, isolation by distance may have led to differentiation of the two geographically disjunct hexaploid species (the mainland *H. iberica*, and the insular *H. maderensis*). In addition, ecological processes may also have been important forces in the evolution of the western polyploid clade as it has been suggested at a macroevolutionary scale for the Asian Palmate clade of Araliaceae, where ivies are included (Valcárcel *et al*. 2014; Valcárcel and Wen 2019). The three western polyploid species of the genus *Hedera* inhabit moist places with low to moderate temperature contrast. Specifically, *H. maderensis* occurs in the laurel forests (laurisilva) of Madeira, and *H. iberica* in several SW Iberian locations frequently considered refugia for subtropical Tertiary relict plants (Rutherford 1989; Costa *et al*. 1997; Calleja *et al*. 2009), while *H. hibernica* spreads towards more northern latitudes under mild oceanic conditions (McAllister 1990).

In this study, we hypothesize that speciation processes in the western polyploid clade of *Hedera* have been driven by a combination of geographical and ecological factors, and that the interaction of both was particularly important for this lineage to pass through the environmental filter in the colonization of Madeira as well as for in situ evolution. To test this hypothesis, we explored phylogenetic relationships, climatic niche preferences and functional diversity in the western polyploid clade of *Hedera*. We sampled the whole distribution range of the three species of the western polyploid clade (plus the diploid outgroup species *H. helix*) and conducted a phylogenomic analysis using genotyping-by-sequencing (GBS). This technique is based on high-throughput sequencing and allows screening of the whole genome (Elshire *et al*. 2011; Escudero *et al*. 2014b), thus helping resolve complex lineages in which a small number of Sanger-sequenced DNA regions provide limited nucleotide variation (Escudero *et al*. 2014b; Fernández-Mazuecos *et al*. 2018, 2020). A georeferenced database of *Hedera* occurrences was used to characterize the climatic preferences of species and estimate climatic niche overlap. To further evaluate the species responses to climate, we also analysed a set of 19 vegetative and regenerative functional traits obtained from leaves and fruits. Our specific objectives were (1) to clarify the phylogenetic relationships among the western polyploid species of *Hedera*, (2) to characterize the differences and similarities in climatic niche preferences of the western polyploid species, (3) to analyse whether the divergence pattern is coupled with functional trait and climatic niche differentiation, and (4) to clarify the evolutionary history and biogeographic origin of *H. maderensis*.

## MATERIALS AND METHODS

### Sampling

A total of 40 populations (92 individuals, 1-5 per population; Supplementary Data Table 1, Fig. 1) were collected for the phylogenomic study: 13 populations *of H. helix* (25 individuals), 11 populations of *H. hibernica* (20 individuals), 9 populations of *H. iberica* (26 individuals) and 7 populations of *H. maderensis* (21 individuals). This sampling strategy covered the whole distribution range of the three species of the western polyploid clade and a good representation of the range of the outgroup species *H. helix* (Fig. 1). Special sampling effort was made to represent the distribution in the Iberian Peninsula of the two most widespread species (*H. helix* and *H. hibernica*) because this is the area where the mainland species come into contact (Valcárcel 2008, 2017) and where the greatest number of plastid haplotypes have been detected (Valcárcel *et al*. 2017).

**Table 1.**
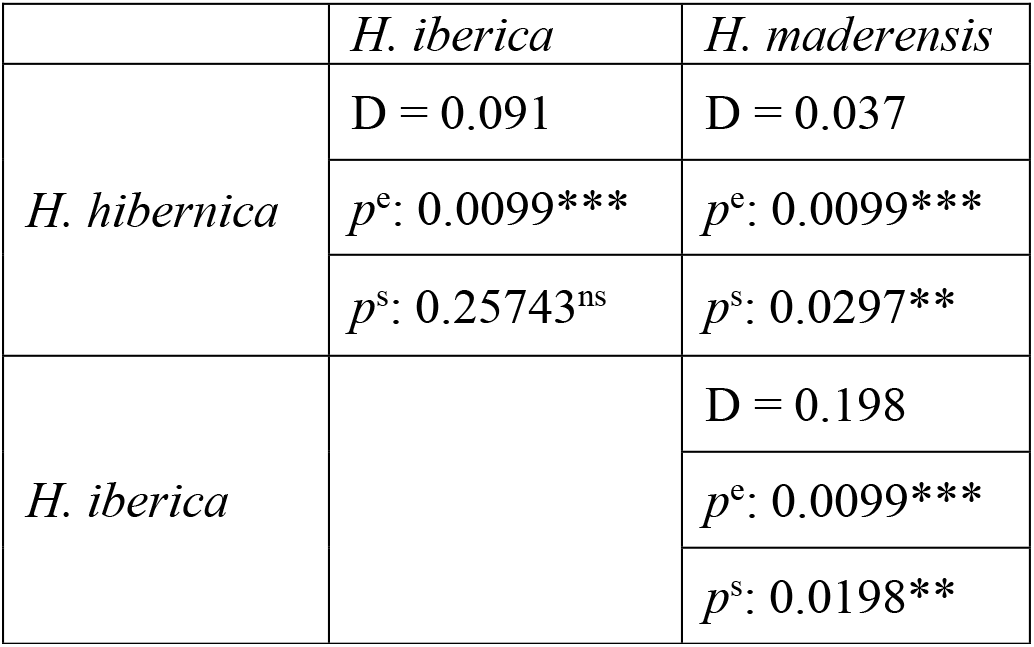
Pairwise climatic niche overlap statistics of the western polyploid clade of *Hedera*. For each species pair, the first row indicates the niche overlap metric D, the second row (*p*^e^) is the *p*-value of the niche equivalency test and the third row is the *p*-value of the similarity test (*p*^s^). Level of significance is shown. ***: p ≤ 0.001, **: p ≤ 0.01, ns: p > 0.05.

A total of 70 populations (335 individuals; Supplementary Data Table 2, map on Fig. 2) were sampled in the Iberian Peninsula and Madeira for the study of functional traits in the western polyploid clade (*H. hibernica*: 52 populations, 248 individuals; *H. iberica:* 9, 53; *H. maderensis*: 9, 34). Ivies have heteroblasty with two distinct phases: the juvenile non-flowering phase (herein called “vegetative phase”) and the adult reproductive phase (herein called “reproductive phase”). The most conspicuous difference between the two phases is the type of growth, which is plagiotropic in the vegetative phase and orthotropic in the reproductive phase (Robbins 1957; Stein and Fosket 1969; Poethig 1990). As a result, the vegetative phase grows as part of the forest understory (it spreads on the ground or climbs up on tree trunks or rocks) and its leaves can be considered shade leaves. Instead, the reproductive phase occurs in the forest canopy and develop sun leaves (Rehm *et al*. 2014). Because of this, for each individual we collected five leaves from the vegetative phase and five from the reproductive phase. Additionally, we also collected five fruits per individual from different umbels.

**Figure 2.**
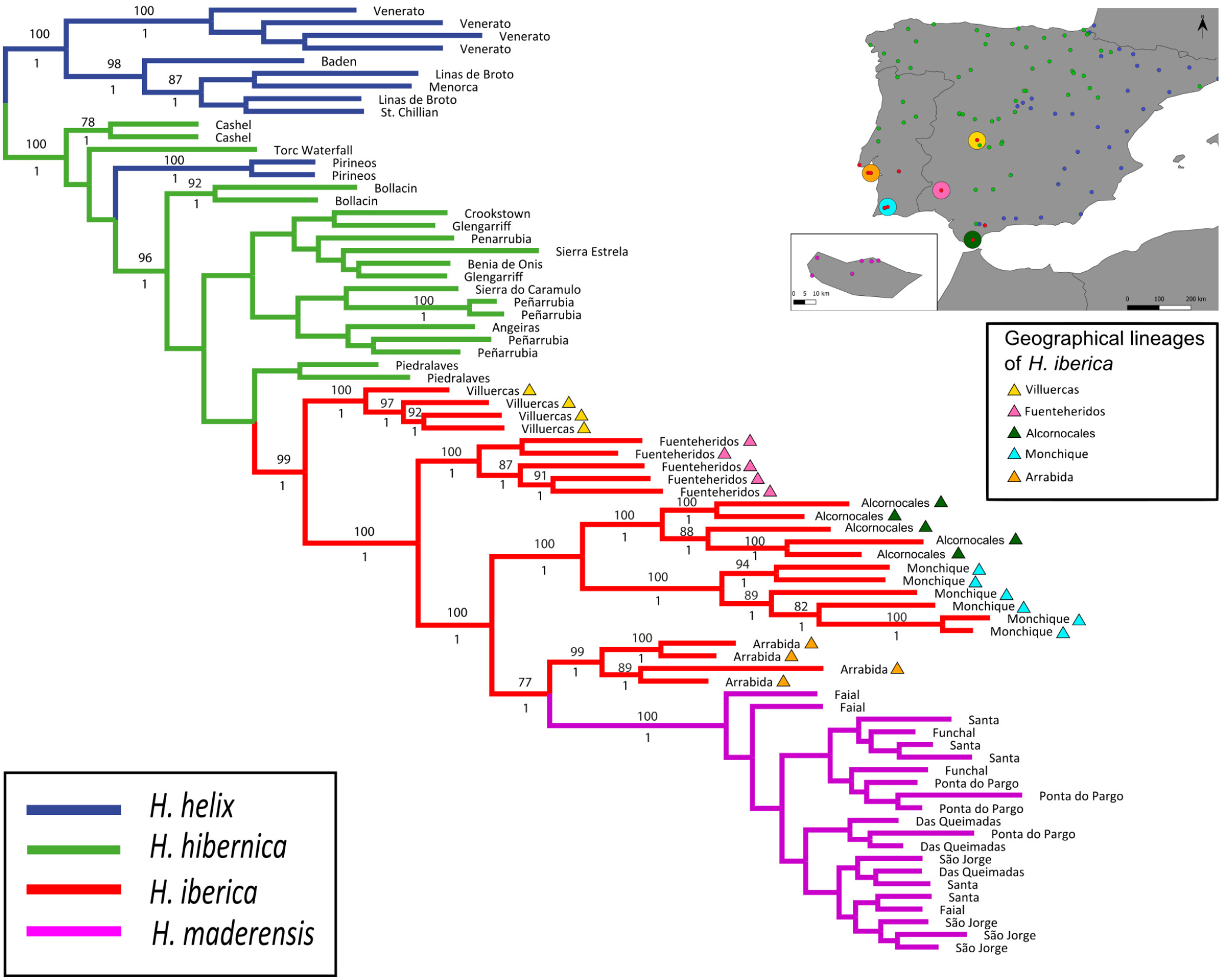
Phylogenetic tree of the western polyploid clade of *Hedera* using *H. helix* as outgroup and based on phylogenomic analysis of GBS data (*c*80*m*15*p*6*r*3 dataset). The maximum-likelihood tree obtained in RAxML is shown, with bootstrap support values ≥75% indicated above branches. Posterior probability values obtained in the Bayesian analysis in ExaBayes are shown below branches. The inset shows a map of the Iberian Peninsula and Madeira indicating sampling localities for the study of functional traits and the geographical location of *H. iberica* lineages.

### Genotyping-by-sequencing library preparation

Total genomic DNA was isolated using a modified CTAB protocol (Doyle and Doyle 1987; Cullings 1992). DNA concentrations were assessed with a Qubit 3.0 Fluorometer (Invitrogen, Carlsbad, CA, USA) using the dsDNA BR Assay Kit. A GBS library was prepared using 500 ng of DNA per sample and the *Pst*I-HF restriction enzyme. We followed the protocol of Fernández-Mazuecos *et al*. (2018), which is based on the original GBS protocol of Elshire *et al*. (2011) and incorporates modifications published by Escudero *et al*. 2014b and Grabowski *et al*. (2014). After the digestion, ligation and pooling steps, DNA fragments were PCR-amplified for 19 cycles using the NEB 2X Taq MasterMix (NEB, MA, USA) and 35 ng of starting DNA in an Eppendorf Mastercycler ep gradient S (Eppendorf, Hamburg, Germany). The amplified library was purified using AMPure XP magnetic beads (Beckman Coulter, CA, USA). Quantification and quality control of the library were carried out using a 2100 Bioanalyzer (Agilient, CA, USA). Additionally, cloning was conducted using a pGEM-T Easy Vector (Promega Biotech Ibérica, Spain), and the inserts of 10 positive colonies were PCR-amplified and sequenced using the Sanger method (Sanger *et al*. 1977) to confirm the correct ligation of adapters. The library was submitted to Macrogen Inc. (Seoul, South Korea) for sequencing using Illumina HiSeq4000 (Illumina, Inc., San Diego, CA, USA) paired-end technology.

### Genotyping-by-sequencing raw data assembly and matrices compilation

GBS matrices were assembled from FASTQ files in seven steps using the ipyrad software (http://www.ipyrad.readthedocs.io) with the methodology for *de novo* assembly (without a reference genome; Eaton 2014; Eaton and Overcast 2020) and treating data as single-end. Assembly parameters followed Fernández-Mazuecos *et al*. 2020, except where indicated. For the demultiplexing step, a maximum of two base mismatches was allowed in barcode sequences. Three different clustering threshold values (*c*: 80, 85 and 90) and two minimum taxon coverage values (*m*: 4 and 15) were used. Additionally, since the study species display three ploidy levels (2x, 4x and 6x), we assembled the data using two alternative values (*p*: 2 and 6) for the maximum number of alleles per locus, corresponding to the maximum and minimum ploidy levels. Therefore, combination of values for these three parameters led to 12 assemblies of GBS loci.

Individuals with low locus recovery showed uncertain positions in preliminary phylogenetic analyses. To avoid the phylogenetic noise caused by these individuals, three matrices were prepared from each of the 12 assemblies: two matrices including only individuals with at least 500 and 1,000 recovered loci in preliminary assemblies (hereafter denoted as *r*1 and *r*2, respectively); and a matrix including individuals with at least 1,000 loci that did not change phylogenetic position across analyses (hereafter denoted as *r*3). Therefore, a total of 36 datasets were generated, each of them denoted as *c*X*m*Y*p*Z*r*W, with X being the clustering threshold, Y the minimum taxon coverage, Z the ploidy level, and W the level of filtering of individuals (see above).

### Phylogenetic analyses

Concatenation-based maximum-likelihood (ML) phylogenies were constructed for all 36 matrices using RAxML-HPC 8.2.12 (Stamatakis 2014) with the GTR+CAT substitution model during tree search, followed by evaluation of the final tree under the GTR+GAMMA model. The number of bootstrap replicates was determined by the bootstopping criterion (Pattengale *et al*. 2010). The highest bootstrap support values for major clades in RAxML analyses were obtained for the *r*3 matrices obtained from the dataset generated using a clustering threshold of 0.80, a minimum taxon coverage of 15 and a maximum number of alleles per loci of 6 (*c*80*m*15*p*6*r*3 dataset). This matrix was selected for all remaining analyses.

We conducted an additional concatenation-based analysis of the the *c*80*m*15*p*6*r*3 dataset using Bayesian inference (BI), implemented in ExaBayes 1.5 (Aberer *et al*. 2014). The GTR+GAMMA substitution model and default priors were used. Two parallel MCMC runs, with two coupled chains each, were carried out for 1.2 million generations, sampled every 500 generations. A parsimony starting tree and default priors were used. Convergence of each run was assessed by evaluating trace plots and ESS values in Tracer 1.7 (Rambaut *et al*. 2018). An extended majority-rule (MR) consensus tree was calculated for each run after discarding a burn-in of 10%. Although the two runs converged to slightly different likelihood values, the resulting MR trees were virtually identical. Therefore, the two runs were combined for calculation of the final MR tree.

We also analysed the *c*80*m*15*p*6*r*3 dataset using the coalescent-based method SVDquartets (Chifman and Kubatko 2014) implemented in PAUP* 4.0a (Swofford 2002). The multispecies coalescent option was used, with exhaustive quartet sampling and eight taxon partitions corresponding to *H. helix, H. hibernica, H. maderensis* and the five disjunct populations of *H. iberica*. We ran 100 bootstrap replicates, and the resulting trees were summarized in a 50% majority-rule consensus tree.

All phylogenetic analyses were carried out through the CIPRES Science Gateway V. 3.3 (http://www.phylo.org).

### Population genetic structure

To evaluate the genetic structure of the study group and test the correspondence between species and genetic clusters, we analysed an unlinked single nucleotide polymorphism (SNP) matrix corresponding to the sequence matrix selected from phylogenetic analyses (*c*80*m*15*p*6*r*3, see above). Genetic PCA was performed using the Jalview 2 software (Waterhouse *et al*. 2009). Bayesian analysis of population structure was carried out in BAPS (Corander *et al*. 2008). Two BAPS analyses were conducted, one of them taxonomically unconstrained and the other taxonomically constrained. For the taxonomically unconstrained analysis, a two-step analysis was conducted to identify the total number of clusters. First, a mixture analysis was run using all individuals, which detected two clusters. Then, two further mixture analyses were conducted to discover finer genetic structure within each of them. As a result of this two-step taxonomically unconstrained approach, we detected a total of three clusters (K=3), which were used as pre-defined populations in a subsequent admixture analysis including all populations together again and a minimum population size of 5, 100 iterations, 200 reference individuals from each population and 20 iterations for reference individuals. For the taxonomically constrained analysis, an admixture analysis was conducted with the same parameters and using the four species as pre-defined populations (K=4).

### Climatic niche study

To characterize the climatic niches and evaluate niche shifts during speciation, the database of Coca et al. (M Coca, UAM, Madrid, Spain, unpubl. res.) was used. This database includes 706 records from the entire geographical ranges of the three species of the western polyploid clade (*H. hibernica*: 189; *H. iberica*: 66 and *H. maderensis*: 25) and *H. helix* (426). In this database, species identification of the records is taxonomically certain because they were all identified by the specialist of the genus (V. Valcárcel). Climatic data were obtained from WorldClim (Hijmans *et al*. 2005) with a resolution of 2.5 min. The 19 bioclimatic variables were clipped in ArcGIS 10.6.1 (ESRI) using the maximum and minimum latitude and longitude coordinates in our database of occurrences. We calculated pairwise Pearson correlations among the 19 bioclimatic variables. To avoid collinearity effects, when two variables had pairwise correlations above 0.8 we excluded one of them. Subsequent analyses were done using the resulting nine variables (bio3: isothermality; bio7: temperature annual range; bio8: mean temperature of wettest quarter; bio9: mean temperature of driest quarter; bio10: mean temperature of warmest quarter; bio11: mean temperature of coldest quarter; bio15: precipitation seasonality; bio16: precipitation of wettest quarter; and bio18: precipitation of warmest quarter).

Climatic niches of the three species of the western polyploidy clade were evaluated using a PCA-env approach implemented in the R package *ecospat* (Warren *et al*. 2008; R Core Team 2013; Di Cola *et al*. 2017). PCA-env is a principal component analysis calibrated using the entire environmental space of the study area (Broennimann *et al*. 2012). Niche overlap for all three species pairs was quantified by means of the *D* metric (Schoener 1970), which was selected due to its long history of use and simplicity of application (Warren *et al*. 2008). *D* values range from 0 (no overlap between niches) to 1 (complete overlap between niches).

Niche equivalency and niche similarity tests (Warren *et al*. 2008; Broennimann *et al*. 2012) were also conducted in *ecospat* for three pairs of species. Niche equivalency tests were used to evaluate niche divergence, i.e. whether the observed niche overlap between two species (*D*) is significantly lower than a null distribution generated by randomly reallocating the occurrences of both species between their ranges (the alternative=“lower” option was used). Specifically, for each pair of species the null distribution was obtained by randomly generating 100 pseudo-replicates that kept the number of occurrences for each species constant and reallocated occurrence between them. The equivalence hypothesis was rejected when the observed *D* was below the lower limit of the 95% confidence interval of the null distribution.

Niche similarity tests were used to evaluate niche conservatism, i.e. whether the observed niche overlap between two species (*D*) is significantly greater than a null distribution obtained by allowing random shifts of the niches within the environmental space (the alternative=“greater” option was used). These tests were based on the chi-square test of Peterson *et al*. (1999) with the modifications described in Warren *et al*. (2008). Null distributions were generated with 100 iterations, and allowing both ranges to be randomly shifted (rand.type=1). Niche conservatism was inferred if the *D* value was above the upper limit of the 95% confidence interval of the null distribution.

R scripts for all niche analyses are available online in Supplementary Data Script 1.

### Functional traits

We analysed six functional traits both on vegetative and reproductive leaves (leaf area, leaf fresh mass, leaf dry mass, leaf dry matter content (LDMC), specific leaf area (SLA) and leaf thickness) and seven regenerative traits (fruit fresh mass, fruit dry mass, fruit dry matter content, seed dry mass, number of seeds per fruit, total seed dry mass per fruit, pulp dry matter content). Sampling, storing and measurements were done following the methodology provided in Pérez-Harguindeguy *et al*. (2013). Mass measurements are provided in grams and area in mm^2^. For fresh weight measures all samples were stored at 4°C in zipped plastic bags with moist paper for the 24 h following field collection to ensure samples were at full turgor. Leaf area and leaf dry mass were measured on pressed leaves after oven-drying samples for 48 h at 70°C. Scaled pictures of leaves were taken and leaf area was calculated with Fiji software (Schindelin *et al*. 2012). Images were converted to 32-bit RGB and to a 3-slice stack (blue, green, red) and blue images were kept because they provided the highest contrast.

SLA was calculated as the ratio of leaf area to leaf dry mass (mm^2^/g). LDMC was computed as the ratio of leaf dry mass to leaf fresh mass. Leaf thickness was measured as leaf fresh mass divided by leaf area. Fruit dry mass was computed after oven-drying the samples for 48 h at 70°C. Seeds were extracted from dry fruits and the number of seeds per fruit were counted and seeds were weighed. Total dry seed mass was calculated per fruit as the sum of the dry masses of all seeds. Fruit dry matter content was estimated as the ratio between fruit dry mass and fruit fresh mass. The pulp dry matter content was computed as the difference between fruit dry mass and total dry seed mass.

The differences in the 19 functional traits among the three species of the western polyploid clade were analysed using ANOVA since data fitted a normal distribution according to the Shapiro–Wilk test (p < .05). Multiple comparisons were done using t student test with the “stat_compare_means” function in the ggpbur (Kassambara 2020). Data were summarized in the form of violinplots using the function “geom_violin” in the ggplot2 package (Wickham 2016).

## RESULTS

### GBS data assembly

Illumina sequencing yielded 514,582,774 paired-end raw reads with a length of 101 bp. The GC content was 41.83% and the percentage of bases with a quality score of Q20 was 96.62%. Numbers of filtered loci, sites (bp), SNPs and phylogenetically informative sites (PIS) and percentages of missing data varied with assembly parameters and kept individuals, as summarized in Supplementary Data Table 3. The dataset selected for extensive analysis (*c*80*m*15*p*6*r*3) had 10,799 filtered loci, 913,235 bp, 53,123 SNPs, 30,793 PIS and 42.4% of missing data. In general, a higher number of loci were recovered for the most recent samples.

### Phylogenetic inference

Results from maximum likelihood and Bayesian inference analyses of the *c*80*m*15*p*6*r*3 dataset were congruent and revealed two well supported main clades, one including most of the samples of *H. helix* (which was used to root the tree) and the other including the three species of the western polyploid clade plus one population of *H. helix* from northern Spain (100% ML-BS, 1 PP; Fig. 2). Within the western polyploid clade, three samples of *H. hibernica* appeared in an early-diverging position together with the *H. helix* population. The remaining samples of *H. hibernica* were clustered with all samples of *H. iberica* and *H. maderensis* in a strongly supported clade (96% ML-BS, 1 PP). Within this clade, samples of *H. hibernica* formed a large basal grade with poorly supported relationships, whereas all samples of *H. iberica* and *H. maderensis* constituted a well-supported clade (99% ML-BS, 1 PP). *Hedera maderensis* appeared embedded within this clade as sister to the Arrabida population of *H. iberica* (77% ML-BS, 1 PP; Fig. 2). Both the ML phylogeny and BI phylogenies supported a strong population and geographical structure in *H. iberica* with diverging lineages corresponding to different populations located in distant geographical areas. Villuercas, the most northern and inland population of *H. iberica*, was the earliest diverging population (100% ML-BS, 1 PP; Fig. 2). Fuenteheridos, the second most inland population of *H. iberica*, formed the second diverging lineage (100% BS, 1 PP; Fig. 2). The next diverging clade included the two populations of *H. iberica* from Alcornocales and Monchique, both located along the southwestern Iberian coast (100% BS, 1 PP; Fig. 2). The latter clade was sister to another clade including the Arrabida population (located on the western Iberian coastline) as sister to *H. maderensis* (Fig. 2). Fully congruent population-level relationships within the *H. iberica*/*H. maderensis* clade were recovered by the coalescent-based SVDquartets analysis, with BS values ranging from 86 to 100% (Supplementary Data Figure 1). The same relationships were supported by most ML analyses using other datasets, but these had generally lower support values (Supplementary Data Figure 2).

### Population genetic structure

The first three components of the genetic PCA (Supplementary Data Figure 3A) accounted for 72% of the variability. Four clusters were identified, corresponding to species delimitation. The *H. maderensis* cluster was clearly isolated, while the other three clusters overlapped to some extent. In particular, the *H. hibernica* cluster was placed in an intermediate position between *H. helix* and *H. iberica* and overlapped with these two species. The narrowest multivariate space of the genetic PCA was occupied by *H. helix*, followed by *H. hibernica, H. maderensis* and *H. iberica*. In the taxonomically unconstrained BAPS analysis with K=3, one cluster included all individuals of *H. helix* and *H. hibernica* as well as the two inland populations of *H. iberica* (Villuercas and Fuenteheridos), another cluster contained *H. iberica* individuals from the three populations closest to the coast, and the third cluster included all the individuals of *H. maderensis*. In the taxonomically constrained BAPS analysis with K=4, the four genetic groups matched species delimitation, with some significant admixture suggested, including admixture from *H. hibernica* into the Villuercas population of *H. iberica* (Supplementary Data Figure 3B).

### Climatic niche

The first two principal components of the climatic PCA accounted for 77.05% of the observed variance (Fig. 3). The first axis mostly accounts for temperature variables (bio9: mean temperature of the driest quarter; bio10: mean temperature of the warmest quarter; and bio11: mean temperature of the coldest quarter), precipitation during the warmest quarter (bio18) and precipitation seasonality (bio15). This axis runs from warm places with high isothermality and dry summers to cold places with low isothermality and rainy summers (right to left axis 1, Fig. 3A). As a result, the first dimension discriminates the three species of the western polyploid clade (Supplementary Data Figure 4A). The second axis mostly accounts for annual temperature range (bio7), precipitation of wettest quarter (bio16) and mean temperature of wettest quarter (bio8). This axis runs from places with low annual temperature contrast, rainy and warm wettest season to places with high annual temperature contrast, and dry and cold wettest season (up to down axis 2, Fig. 3A). The climatic niches of the species of *Hedera* occur on the upper half of this axis (positive values of axis 2; Fig. 3A). This second dimension discriminates *H. maderensis* that occurs on the upper part of *H. iberica* that tends to concentrate at lower values of the axis (Supplementary Data Figure 4B). In all cases, except for *H. maderensis*, species density plots show more than one peak (Fig. 3A). *Hedera helix* and *H. hibernica* are the species with the broadest niches followed by *H. iberica* and *H. maderensis. Hedera helix* occupies the central part of the multivariate space and comprises a broad range of environmental conditions, including cold areas with low isothermality and rainy summers (left density peak) and warm areas with high isothermality and dry summers (right density peak). The niche of *H. hibernica* does not include the coldest areas occupied by *H. helix*, but also occurs in contrasting environments including areas with low annual temperature contrast and a relatively cold and rainy wettest season (upper density peak) and areas with higher contrast in annual temperature and with a warmer and drier wettest season (lower density peak). The climatic niche of *H. iberica* includes three density peaks, all occurring in the warmest environments occupied by the ivies in Europe and spreading across a gradient from low annual temperature contrast and relatively mild and rainy wettest season (upper density peak; Alcornocales, Monchique and Arrabida, see map in Fig. 2) to more contrast in annual temperature areas with warm and dry wettest season (lower density peak; Fuenteheridos, see map in Fig. 2) The niche of *H. maderensis* only has one density peak and is placed at one of the extremes of the European climatic space of ivies in warm areas with a rainy wettest season and high isothermality (upper right part). The analysis of the climatic niche overlap (Table 1) reveals certain degree of overlap between all species pair comparisons with the lowest value of overlap detected between *H. hibernica* and *H. maderensis* (D = 0.037) and the highest value between *H. iberica* and *H. maderensis* (D = 0.198). The climatic niches of the species are not equivalent (Table 1). Climatic niche similarity is detected between *H. maderensis* and the other two species of the western polyploid clade (Table 1).

**Figure 3.**
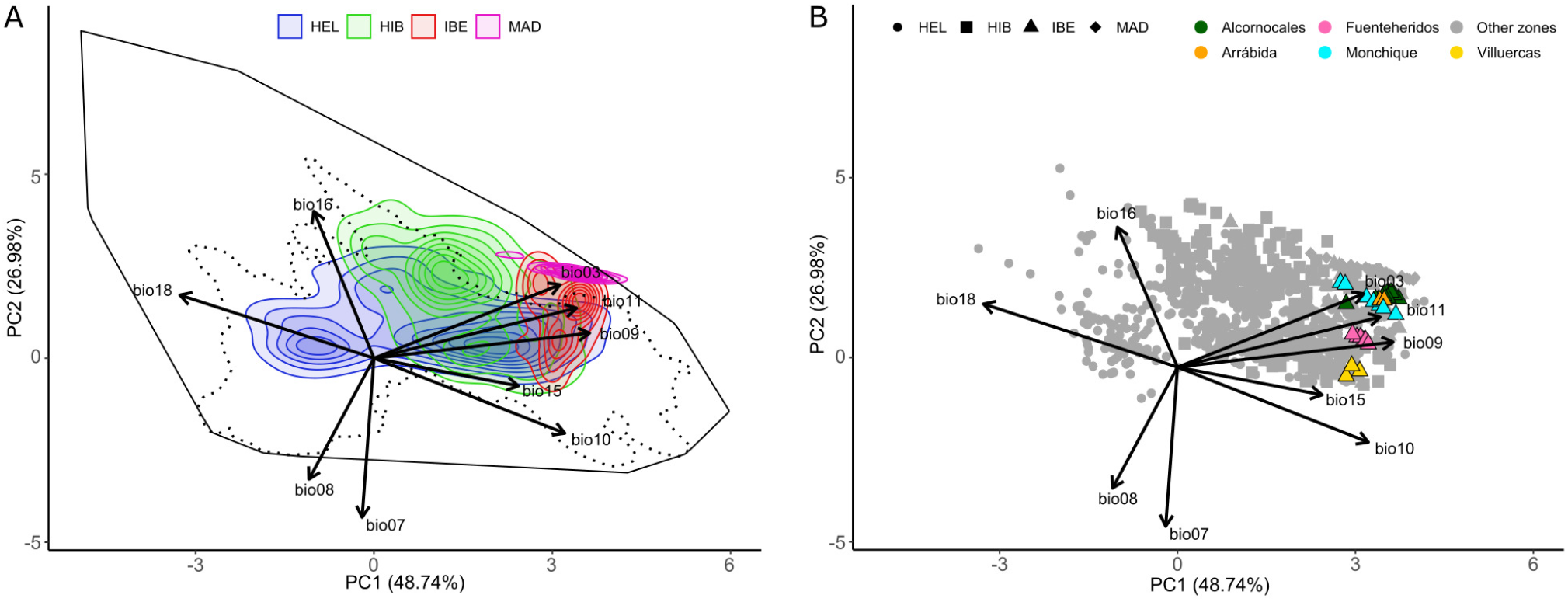
Climatic niche ordination analysis. (A) Climatic niche space of the three species of the western polyploidy clade of *Hedera* (*H. hibernica, H. iberica, H. maderensis*) and *H. helix* as inferred from their entire geographical ranges. The colours of the density envelopes correspond to species, with shading intensity indicating the density based on point occurrences. The solid black line delimits the multivariate space corresponding to the 100% of the available environment for all the species, and the dotted line contains the 95% of the available environment. (B) Climatic space occupied by the diverging lineages of *H. iberica* according to the GBS phylogeny (Fig. 2). Contributions of the original WorldClim variables are provided: bio03, isothermality; bio7, temperature annual range; bio8, mean temperature of wettest quarter; bio9, mean temperature of driest quarter; bio10, mean temperature of warmest quarter; bio11, mean temperature of coldest quarter; bio15, precipitation seasonality; bio16, precipitation of wettest quarter; bio18, precipitation of warmest quarter.

### Functional traits

Significant differences were detected in almost all the functional traits analysed when comparing *H. hibernica* to *H. iberica* and *H. maderensis*, whereas the latter two species only differed in seven out of the 19 traits analysed (Fig. 4, Supplementary Data Figure 5). On the one hand, fruits and seeds of *H. hibernica* are bigger in size and have greater pulp dry matter content than those of the other two species, although no difference is detected for the number of seeds per fruit and the total seed dry mass per fruit (Fig. 4A-D). On the other hand, the leaves of *H. hibernica* (reproductive and vegetative) are smaller in size (low values of leaf area and leaf dry mass), denser (high LDMC) and with lower SLA and thickness values than those of *H. iberica* and *H. maderensis*. On the contrary, the fruits and seeds of *H. iberica* and *H. maderensis* do not show any significant differences, except for the seed size, which is lower in *H. maderensis* (Fig. 4C). The leaves of *H. maderensis* are smaller in size (leaf area, leaf dry mass, Fig. S4D, E, G) than those of *H. iberica*, but no difference is detected for any of the other leaf traits measured.

**Figure 4.**
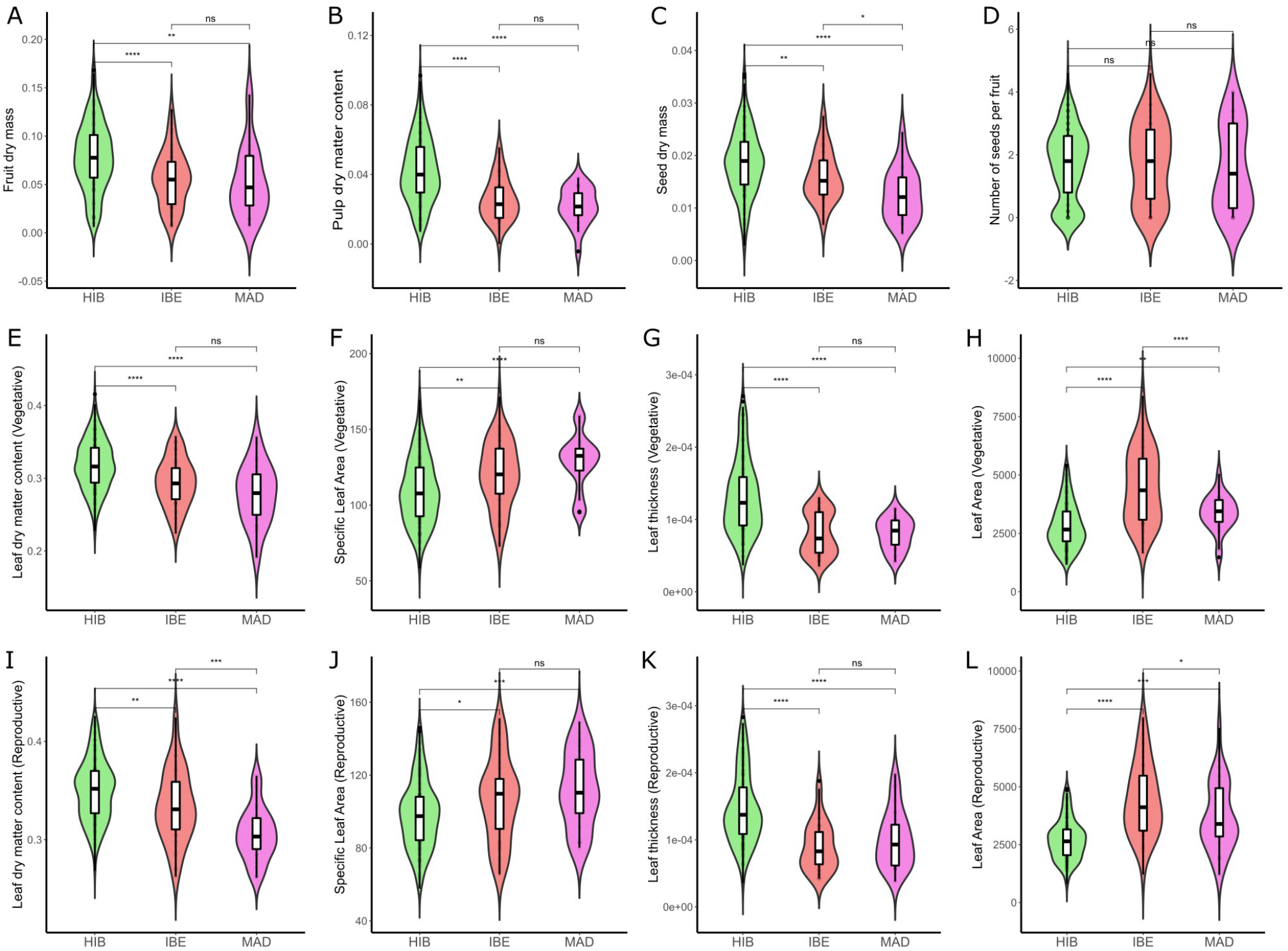
Vegetative and regenerative functional trait differences among the species of the western polyploid clade of *Hedera*: *H. hibernica* (HIB), *H. iberica* (IBE), and *H. maderensis* (MAD). The crossbar within the boxplot shows the median, and the length of the box indicates the interquartile range. Shape of the violinplot reflects the kernel density plot of data. Pairwise comparisons are indicated with the correspondent p.value. Level of significance is shown. ****: p ≤ 0.0001, ***: p ≤ 0.001, **: p ≤ 0.01, *: p ≤ 0.05, ns: p > 0.05.

## DISCUSSION

### Budding speciation in the western polyploid clade of *Hedera*

The application of the GBS technique has helped elucidate species-level phylogenetic relationships within the western polyploid clade of *Hedera* and has fully resolved population-level relationships in the *H. iberica*/*H. maderensis* clade. Indeed, our results clarify the evolutionary history of the group by revealing a nested phylogenetic pattern of speciation in which monophyly is only recovered for one of the three species of the clade (*H. maderensis*). The progenitor-derivative species relationship typical of nested speciation (Crawford 2010) was robustly supported for *H. iberica* and *H. maderensis* (Fig. 2). The monophyletic *H. maderensis* is embedded within *H. iberica*, thus making the latter species paraphyletic. Although reciprocal monophyly is not observed, the two species are clearly distinct in macro- and micromorphological characters (Ackerfield and Wen 2003; Valcárcel and Vargas 2010), and they are also distinct genetically (Supplementary Data Figure 3). The lack of reciprocal monophyly can therefore be attributed to a case of budding speciation. This type of speciation was originally proposed by Mayr (1954) as “peripatric speciation” to describe the differentiation of a new species (budded species) from a small new population that achieves reproductive isolation after dispersal from a widespread parental species. This concept essentially describes an evolutionary process in which a new species is originated through evolutionary change without the extinction of the parental species (“budding”, Mayr and Bock 2002). In the short-term, this results in a progenitor-derivative species pair scheme rather than a true sister relationship (Crawford 2010). In fact, monophyly is not expected for the parental species soon after the speciation event because sorting is not likely to be completed (Neigel and Avise 1986), while monophyly is expected for the budded species early in its evolution because few individuals are involved in the origin, and evolutionary change is accelerated via founder effect and genetic drift (Rieseberg and Brouillet 1994). In our study case, it is probable that the recovery of paraphyly for the parental species (*H. iberica*) is due to its relatively large and patchy range (68 km^2^ of area of occupancy vs. 139.708 km^2^ of extent of occurrence; M Coca, UAM, Madrid, Spain, unpubl. res.) rather than a mere lack of time to coalesce, given that a middle Pliocene divergence time has been estimated for the western polyploid clade (Valcárcel *et al*. 2017). Indeed, strong geographical structure is detected, with diverging lineages matching populations located in distant geographical areas (Fig. 2). In such cases of high levels of geographic differentiation, achieving complete lineage sorting for a majority of the genome (thus leading to monophyly in phylogenomic analyses) may be impeded even over long periods of time (Rieseberg and Brouillet 1994; Funk and Omland 2003). Interestingly, the area of occupancy of *H. iberica* is twice the area of *H. maderensis* (68 km^2^ vs. 28 km^2^, M Coca, UAM, Madrid, Spain, unpubl. res.), which is consistent with predictions of budding speciation where an asymmetry in the ranges is expected, with widespread parental *vs*. restricted budded species (Fitzpatrick and Turelli 2006; Anacker and Strauss 2014; Barraclough and Vogler 2000). Also, the climatic space occupied by *H. maderensis* is almost three times less than that of *H. iberica* and overlaps with one of its extremes (Table 1, Fig. 3A). This parallelism in the geographical range and niche breadth asymmetries is also typical of budding speciation (Grossenbacher *et al*. 2014) due to the positive correlation between the extent of geographical range and realized niche breadth (Slatyer *et al*. 2013).

In the past, budding speciation was considered a rather unusual speciation model. As molecular phylogenies have become routine in systematic studies, evidence of budding speciation has greatly increased and this phenomenon is now accepted as a common evolutionary scenario for plant speciation (Gottlieb 2004; Crawford 2010; Hörandl and Stuessy 2010). However, the frequency of this mode of speciation is unevenly distributed across phylogeny and geography (Grossenbacher *et al*. 2014). For example, organisms with dispersal limitations or strong local adaptation are more prone to speciate following a budding-off pattern (i.e., *Mimulus*, Grossenbacher *et al*. 2014). In the same way, regions with heterogeneous landscape and complex geology may concentrate cases of budding speciation because isolation and local adaptation are facilitated (i.e. California Floristic Province, Anacker and Strauss 2014). Similarly, oceanic islands like Madeira may favour budding speciation (Vanderpoorten and Long 2006). Our results also suggest that the *H. iberica/H. maderensis* clade may have evolved as another progenitor-derivative case from *H. hibernica*. Nevertheless, the exact way in which this happened remains an open question. This is partly due to the poor internal resolution of *H. hibernica*, but also to the potential introgression with *H. helix* (Fig. 2), which may have obscured evolutionary patterns at the early steps in the evolution of the western polyploid clade. A more comprehensive and targeted sampling of *H. helix* and *H. hibernica* is needed to evaluate the extent and timing of the introgression and further clarify the patterns of speciation and population differentiation within the western polyploid clade of *Hedera*.

### Adaptive island colonization and non-adaptive in situ evolution of *H. maderensis*

Most of the studies of adaptive pressures on oceanic islands have been focused on the effects of ecology on in situ evolution and its relevance as a driver of speciation (Emerson 2002), while its effects during the preliminary steps of the colonization process have received comparatively little attention (Patiño *et al*. 2017). Besides, our knowledge on adaptive pressures is biased towards radiating lineages (Losos and Ricklefs 2009) while we know little about the in situ speciation processes of non-radiating insular lineages (Patiño *et al*. 2017). In this study, we provide insights on the impact of ecological drivers on the dispersal, colonization and speciation processes of the Madeiran ivy. To do so, we have combined information from the phylogeny and environmental niches together with regenerative and vegetative traits of the studied species.

Regarding the dispersal step in the colonization of the island, monophyly of *H. maderensis* supports a single successful event of dispersal from the continent, and the close relationship between *H. maderensis* and the coastal population of *H. iberica* in Arrabida (Central Portugal) suggests an origin of dispersal in that geographic region. This route involves travelling 1,000 km over the Atlantic Ocean. Assuming endozoochorous dispersal, relatively big fleshy fruits would be expected because large birds select larger fruits (Jordano 2000) and tend to travel longer distances (Jordano *et al*. 2007). Interestingly, the dispersal event that led to the colonization of Madeira does not fit this assumption. The analysis of the regenerative traits shows that the fruits of *H. maderensis* are similar in size to those of the parental species *H. iberica*, and happen to be smaller than those of *H. hibernica* (Fig. 4A-B). According to available evidence, small (warblers, robins, starlings) and medium-sized frugivorous birds (blackbirds) are the most frequent dispersers of ivies (Guitian 1987; Pasinelli 2003; Heleno *et al*. 2011). Since some of these small frugivores occur both in the Iberian Peninsula and Madeira (Bourne 1984; Jepson and Zonfrillo 1986; Zino *et al*. 1995), and small birds tend to select smaller fruits (Jordano 2000) it is possible that the dispersal event was driven by small birds. This points to a disperser-mediated selection towards small fruits in the western polyploid clade of *Hedera* as a key step to pass through the dispersal filter and successfully colonized the island of Madeira.

Regarding the establishment of the lineage in Madeira, our findings show that *H. maderensis* represents the last step of gradual climatic niche differentiation in *H. iberica* that brought about the pre-adaptations needed for the colonization of Madeira. Upon arrival in Madeira, these pre-adaptations and the climatic similarity between mainland and island localities contributed to population establishment, followed by speciation without further local diversification. The Madeiran ivy occupies a narrow climatic niche restricted to the wettest and least continental areas of the climatic space of *H. iberica* (Fig. 3A). Rather than displaying a monotonic space, the niche of *H. iberica* shows three density peaks (Fig. 3A). Interestingly, these three peaks mimic the geographic pattern observed in the phylogenetic tree (Fig. 2). Indeed, the early-diverging populations in the tree (Villuercas and Fuenteheridos) are located far away from the coast (Fig. 2) and occupy the coldest and most continental areas within the climatic preferences of *H. iberica* (Fig. 3B), whereas later-diverging populations (Monchique, Alcornocales and Arrabida) occur close to the coast (Fig. 2) and occupy the warmest and least continental areas within the species preferences (Fig. 3B). The latter three populations, among which the sister lineage to *H. maderensis* is included (Arrabida), occupy the warmest and the most oceanic part of the climatic niche of *H. iberica*, coinciding with the niche of *H. maderensis* (Fig. 3A). This climatic gradient in *H. iberica* and *H. maderensis* is part of a broader evolutionary pattern in the western polyploidy clade starting from the putatively ancestral cold environments occupied by *H. hibernica* and leading to the colonization of progressively warmer areas. Evolutionary patterns that include gradual climatic niche changes congruent with phylogenetic spatial divergence and budding speciation could be more common than previously thought (*i*.*e*., Otero *et al*. 2019) but they have rarely been reported for the origin of Macaronesian lineages mostly because they have not been explicitly tested.

The geographically structured variation in the climatic preferences of the western polyploid clade across a spatial phylogenetic pattern is also accompanied by a congruent divergence in the species functional traits. Functional divergence associated with climate has been well documented for plant radiations on oceanic islands (*Aeonium*, Jorgensen and Olesen 2001; *Sonchus* alliance, Santiago and Kim 2009; Hawaiian lobeliads, Givnish *et al*. 2009), while examples focused on non-radiating lineages like the Madeiran ivy are more limited (but see *Periploca* and Kleinia, García-Verdugo *et al*. 2019, 2020). In our case study, *H. hibernica* displays small leaves with low SLA and high LDMC and thickness (Figs. 4E-L, Supplementary Data Figure 5), which is related to high investment in the structural component of leaves and indicates that the species has more “conservative” leaves (Díaz *et al*. 2016). In contrast, *H. iberica* and *H. maderensis* have more “acquisitive” leaves, that is, relatively big leaves with high SLA and low LDMC and thickness (Figs. 4E-L, Supplementary Data Figure 5). These acquisitive leaves are cheap in terms of economic investment and have higher productivity and shorter lifespans (Díaz *et al*. 2016) which are typical of the mild climatic conditions with low abiotic stress occupied by the species. Altogether, these results point to divergent selection across habitats (Anacker and Strauss 2014) as a determinant process in the evolution of the mainland lineages of the western polyploid clade resulting in a transition from more “conservative” to more “acquisitive” leaves (Fig. 4E-L). However, the directionality of the mainland functional divergence is broken upon the establishment in the more oceanic conditions of Madeira. Indeed, the leaves of the island lineage (*H. maderensis*) are smaller in size than those of its mainland parental species (*H. iberica*, Fig. 4H, 4L) which challenges the expected increase in leaf size from mainland to island species (García-Verdugo *et al*. 2010, 2019; Burns *et al*. 2012). Also, contrary to expectations that predict an increase in leaf traits related to photosynthesis and growth efficiency in the island taxa due to the more oceanic conditions (García-Verdugo *et al*. 2010), the leaf economic spectrum (SLA, LDMC and thickness) of *H. iberica* and *H. maderensis* does not show significant differences. Yet, this result is not surprising because the oceanic climatic niche of *H. maderensis* is embedded within that of *H. iberica* (Fig. 3A). This suggests that the mainland populations were well adapted to the conditions found in Madeira prior to the migration event, and points to some sort of niche pre-adaptation that may have facilitated establishment after dispersal, and therefore the success of island colonization. Furthermore, the fact that the smaller leaves of *H. maderensis* display an economic spectrum similar to that of *H. iberica* points to a proportional reduction of the investment in structural tissues (leaf density and thickness) in *H. maderensis* (Figs. 4G, 4K) to compensate the reduction in size and keep the photosynthesis and growth efficiency under the Madeiran’s climate.

Finally, the prior acquisition of the adaptations to mild climates in the mainland populations marked the evolutionary path of the lineage once on the island. Indeed, speciation of *H. maderensis* with respect to the parental *H. iberica* was probably driven by geographic isolation between island and mainland populations, while the economic leave spectrum remained unchanged (Fig. 4) likely as a consequence of the climatic similarity between the source population in coastal Iberia and Madeira. This lack of difference is striking because there are major morphological differences between the two species in other features less affected by selective pressures (Ackerfield and Wen 2003; Valcárcel and Vargas 2010). It is intriguing why no further in situ diversification has occurred in Madeira, since dispersal limitations seem to boost local adaptations for fleshy fruit plants in tropical forests (Givnish *et al*. 2009). Indeed, an additive positive effect of small fruits and islands on the speciation rate of tropical palms has been reported (Onstein *et al*. 2017). However, in a study conducted in Macaronesia (Patiño *et al*. 2014), the laurel forest appeared as the ecosystem with the highest proportion of single-species endemics. A lack of island radiation seems to be the dominant pattern for ivies considering that the other two Macaronesian species (*H. canariensis* and *H. azorica*) are single-species endemics as well. Since all of the Macaronesian ivies occur in the laurel forest, we wonder if climatic niche pre-adaptation has played a role in the successful colonization of the three Macaronesian archipelagos and if this pre-adaptation also had a key role in the lack of in situ diversification.

Our preliminary hypothesis stated that ecological processes were key in the speciation of *H. maderensis* because islands are usually associated with in situ local adaptations (García-Verdugo *et al*. 2019). While our results indicate that ecology was key to the colonization of Madeira, they also discard ecology as a main driver in the speciation of *H. maderensis*. To our knowledge, this is the first evidence that explicitly supports mainland pre-adaptation in a Macaronesian plant lineage, which is not totally surprising given that most of the populations of *H. iberica* occur in mainland Iberian locations that have been frequently considered as climatically and floristically related to Macaronesia (Rutherford 1989; Costa *et al*. 1997; Calleja *et al*. 2009). However, this mainland pre-adaptation contrasts with previous findings that show phenotypic divergence between sister plant lineages in Macaronesian islands and the mainland suggesting adaptive pressures for in situ evolution (García-Verdugo *et al*. 2019; García-Verdugo *et al*. 2020). Future research will examine the prevalence of this evolutionary pattern in single-species island endemics, and particularly in Macaronesian ivies.

## ACKNOWLEDGMENTS

The authors specially thank M. Sequeira for his essential field assistance during our collecting trip in Madeira, J.A. Calleja for his free-handed contribution of crucial information on large ivy populations in Spain, M. Leo, G. García-Saúco, P. Bella, F. Lara and D. Gómez for providing on ivy populations, as well as Emilio Cano for laboratory support.

## SUPPLEMENTARY MATERIAL

Supplementary data consist of nine files. Supplementary script 1: R script employed for the niche overlap analyses. Supplementary table 1: List of plant material used for the molecular study. Supplementary table 2: List of plant material used for the functional trait study. Supplementary table 3: Characteristics of 36 genotyping-by-sequencing datasets used in phylogenetic analyses. Supplementary figure 1: Consensus trees of the RAxML analyses from the 36 datasets. Supplementary figure 2: Colaescent-based consensus tree. Supplementary figure 3: Results from population genetic structure analyses (genetic PCA and BAPS). Supplementary figure 4: Violinplots of the functional traits. Supplementary figure 5: Violinplots of the first two axes of the climatic PCA.

