## Supplementary material for "Climatic niche pre-adaptation in mainland Europe facilitated the colonization of Madeira by ivies (*Hedera* L., Araliaceae)"

Supplementary table 1. List of plant material used for the molecular study indicating the assigned SVDQuartet. Abbreviations of the SVDQuartet groups are as follows: ALC represents the *H. iberica* quartet from S Spain; ARR represents the *H. iberica* quartet from SW Portugal; EST represents the *H. hibernica cf.* quartet from C Portugal; FUE represents the *H. iberica cf.* quartet from SW Spain; HEL the *H. helix* quartet; HIB represents the *H. hibernica* quartet; MAD represents the *H. maderensis* quartet; MON represents the *H. iberica* quartet from S Portugal; PIE represents the *H. hibernica cf.* quartet from C Spain; VIL represents the *H. iberica cf.* quartet from C Spain. All specimens are kept at MAUM herbarium in Universidad Autónoma de Madrid.

| Species | Population Number (N individuals) | Country | Locality | Voucher | Coordinates |
|---|---|---|---|---|---|
| *H. helix* | 1(2) | Austria | Niederösterreich, Baden | Valcárcel, V. 43VV03(4,10) | 48.007103, 16.20197 |
| *H. helix* | 2(1) | Belgium | Hainaut, Charleroi | Vargas, P. 114PV03(1) | 50.465588, 4.424228 |
| *H. helix* | 3(2) | France | Pyrénées-Atlantiques, Gabas | Vargas, P. 338PV02(1,7) | 42.817114, -0.403234 |
| *H. helix* | 4(1) | France | Occitanie, Saint-Chinian | Vargas, P. 229PV06 | 43.426471, 2.934946 |
| *H. helix* | 5(5) | Greece | Creta, Venerato | Martín Bravo, S. 338SMB05(1-5) | 35.195155, 25.042142 |
| *H. helix* | 6(2) | Italy | Lazio, Gaeta | Vargas, P. 209PV01(3,9) | 41.213530, 13.576462 |
| *H. helix* | 7(1) | Spain | Almería, Fondón | Vargas, P. 12PV05(1) | 36.948539, -2.869739 |
| *H. helix* | 8(1) | Spain | Burgos, Atapuerca | Vargas, P. 129PV01(1) | 42.350021, -3.519477 |
| *H. helix* | 9(2) | Spain | Cantabria, Santoña | Vargas, P. 125PV01(5,10) | 43.447193, -3.477687 |
| *H. helix* | 10(2) | Spain | Huesca, Linas de Broto | Vargas, P. 335PV02(1,9) | 42.625886, -0.149739 |
| *H. helix* | 11(2) | Spain | Menorca, Cala Galdana | Vargas, P. 329PV02(1,7) | 39.938619, 3.936604 |
| *H. helix* | 12(2) | Spain | Málaga, Valle de Abdalajís | Valcárcel, V. 8VV02(2,12) | 36.939987, -4.68868 |
| *H. helix* | 13(2) | Switzerland | Vaud, Lausanne | Vargas, P. 132PV04(3,6) | 46.520887, 6.57852 |
| *H. hibernica* | 1(2) | Ireland | Tipperary, Cashel | Vargas, P. 180PV10(1,5) | 52.516994, -7.891180 |
| *H. hibernica* | 2(1) | Ireland | Cork, Crookstown | Vargas, P. 170PV10 | 51.843063, -8.830303 |
| *H. hibernica* | 3(2) | Ireland | Cork, Glengarriff | Vargas, P. 177PV10(1,5) | 51.750558, -9.552308 |
| *H. hibernica* | 4(2) | Ireland | Kerry, Torc Waterfall | Vargas, P. 171PV10(1,5) | 52.007438, -9.507470 |
| *H. hibernica* | 5(1) | Portugal | Lavra, Angeiras | Fiz, O. 221OF00 | 41.267134, -8.710806 |
| *H. hibernica cf.* | 6(1) | Portugal | Loriga, Serra da Estrela | Fiz, O. 220OF00 | 40.322023, -7.6132 |

| | | | | | |
|---|---|---|---|---|---|
| *H. hibernica* | 7(1) | Portugal | Viseu, Serra do Caramulo | Ribeiro, P. 336PR | 40.588686, -8.166273 |
| *H. hibernica* | 8(1) | Spain | Asturias, Benia de Onís | Vargas, P. 412PV00 | 43.33634, -4.96627 |
| *H. hibernica cf.* | 9(2) | Spain | Ávila, Piedralaves | Valcárcel, V. 02VV18(1*,10) | 40.340236, -4.70847 |
| *H. hibernica* | 10(2) | Spain | Cantabria, Bollacín | Vargas, P. 127PV01(3,10) | 43.04324, -3.87686 |
| *H. hibernica* | 11(5) | Spain | León, Carucedo | Nieto Feliner, G 4615(2,4,6,7,8). | 42.49166, -6.76109 |
| *H. iberica* | 1(1) | Portugal | Setúbal, Arrabida | Valcárcel, V. 04VV18(1) | 38.50551, -9.149715 |
| *H. iberica* | 2(4) | Portugal | Setúbal, Arrabida | Valcárcel, V. 05VV18(2,4,7,9) | 38.497361, -9.054198 |
| *H. iberica* | 3(1) | Portugal | Algarve, Monchique | Valcárcel, V. 06VV18 | 37.342989, -8.484002 |
| *H. iberica* | 4(1) | Portugal | Algarve, Foia peak | Valcárcel, V. 07VV18 | 37.342989, -8.484002 |
| *H. iberica* | 5(2) | Portugal | Algarve, Monchique | Valcárcel, V. 08VV18(1,5) | 37.307021, -8.58707 |
| *H. iberica* | 6(2) | Portugal | Algarve, Foia peak | Valcárcel, V. 09VV18(1,2) | 37.314972, -8.591478 |
| *H. iberica cf.* | 7(5) | Spain | Cáceres, Villuercas | Valcárcel, V. 03VV18(1,2,3,4*,5*) | 39.620785, -5.439747 |
| *H. iberica* | 8(5) | Spain | Cádiz, Alcornocales | Valcárcel, V. 10VV18(1,3,5,6,8) | 36.22511, -5.582609 |
| *H. iberica cf.* | 9(5) | Spain | Huelva, Fuenteheridos | Valcárcel, V. 11VV18 (1*,3*,5*,7*,10*) | 37.90834, -6.658308 |
| *H. maderensis* | 1(3) | Portugal | Madeira, Santana | Valcárcel, V. 01VV08(1-3) | 32.794021, -16.870569 |
| *H. maderensis* | 2(3) | Portugal | Madeira, Das Queimadas | Valcárcel, V. 03VV08(1-3) | 32.782955, -16.906232 |
| *H. maderensis* | 3(4) | Portugal | Madeira, São Vicente | Valcárcel, V. 04VV08(1-4) | 32.80527, -17.016722 |
| *H. maderensis* | 4(5) | Portugal | Madeira, Achadas da Cruz | Valcárcel, V. 05VV08(2-6) | 32.841595, -17.209293 |
| *H. maderensis* | 5(4) | Portugal | Madeira, Ponta do Pargo | Valcárcel, V. 06VV08(1-4) | 32.811861, -17.247745 |
| *H. maderensis* | 6(1) | Portugal | Madeira, São Vicente | Navarro, C. 3394CN | 32.756727, -17.091288 |
| *H. maderensis* | 7(1) | Portugal | Madeira, Santana | Velayos, M. 9818MV | 32.821443, -16.881931 |

Supplementary table 2. List of plant material used for the functional trait study. All specimens are kept at MAUM herbarium in Universidad Autónoma de Madrid.

| Locality | Voucher | Coordinates |
|---|---|---|
| ***Hedera hibernica*** | | |
| Spain, Sevilla, Las Navas de la Concepción | 01VV20 (1-5) | 37.926989, -5.489869 |
| Spain, Ávila, Piedralaves | 02VV18 (1-10) | 40.340236, -4.70847 |
| Spain, Cádiz, Grazalema, Benamahoma | 02VV20 (1-5) | 36.770783, -5.478387 |
| Spain, Segovia, Sepúlveda | 03AG20 | 41.29317, -3.776106 |
| Spain, Zaragoza, Moncayo | 03AG21 (1-5) | 41.811524, -1.819827 |
| Spain, Cáceres, Guadisa | 03VV19 (1-5) | 39.375049, -5.028613 |
| Spain, Segovia, Villaseca | 04AG20 (1-5) | 41.292368, -3.842561 |
| Spain, Zaragoza, Anento | 04AG21 (1-5) | 41.071658, -1.327443 |
| Spain, Salamanca, Las Casas del Conde | 05AG20 | 40.505107, -6.045838 |
| Spain, Salamanca, Mogarraz | 06AG20 (1-5) | 40.505084, -6.045221 |
| Spain, Ávila, Navatejares | 08AG20 (1-5) | 40.328257, -5.522129 |
| Spain, Barcelona, Cerdanyola del Vallès | 08AG21 (1-5) | 41.445822, 2.129541 |
| Spain, Gerona, Calonge | 10AG21 (1-5) | 41.897548, 3.058039 |
| Spain, Ciudad Real, Fuencaliente | 10VV20 (1-5) | 38.422496, -4.297459 |
| Spain, Albacete, Tarazona de la Mancha | 11AG20 (1-5) | 39.562831, -4.584926 |
| Spain, Gerona, Castellón de Ampurias | 11AG21 (1-5) | 42.222625, 3.089219 |
| Spain, Córdoba, Trasierra | 11VV20 (1-5) | 37.940796, -4.895269 |
| Spain, Toledo, Navas del Estena | 12AG20 (1-5) | 39.488786, -4.610263 |
| Spain, Segovia province | 12VV18 (1,2) | 40.953708, -4.126171 |
| Spain, Cáceres, Guadalupe | 13AG20 (1-5) | 39.442638, -5.351212 |
| Spain, Soria, Velilla de Medinaceli | 14AG20 (1-5) | 41.157843, -2.34093 |
| Spain, Zaragoza, Huérmeda | 14AG21 (1-5) | 41.388157, -1.594476 |
| Spain, Orense, Ginzo da Limia | 14VV19 (1-5) | 42.073943, -7.707421 |
| Spain, Soria, Valdelavilla | 15AG20 (1-5) | 41.971229, -2.205101 |
| Spain, Orense, Ribadavia | 15VV19 | 42.303708, -8.130766 |
| Portugal, Braga, Caldas de Jerez | 16AG21 (1-5) | 41.760674, -8.149292 |
| Spain, Pontevedra, A Barosela | 16VV19 (1-5) | 42.565822, -8.608592 |
| Portugal, Viseu, Castro Daire | 17AG21 (1-5) | 40.920817, -7.962651 |
| Spain, A Coruña, Carnota | 17VV19 (1-5) | 42.853045, -9.070197 |
| Portugal, Guarda, Manteigas | 18AG21 (1-5) | 40.416113, -7.527149 |
| Spain, Lugo, Germade | 18VV19 (1-5) | 43.402009, -7.786319 |
| Portugal, Serra do Açor, Benfeita, Fraga da Pena waterfall | 19AG21 (1-5) | 40.22032354, -7.936101732 |
| Spain, Galicia, Lugo, Nadela, service road to San Mamede | 19VV19 (1-5) | 42.977833, -7.516703 |
| Spain, Burgos, Covarrubias | 20AG20 (1-5) | 42.052734, -3.558726 |
| Portugal, Leiría, Louriçal | 20AG21 (1-5) | 40.014277, -8.785078 |
| Spain, Asturias, Tapia de Casariego | 20VV19 (1-5) | 43.56304, -6.912319 |
| Spain, Burgos, Valle de Sedano | 21AG20 (1-5) | 42.777558, -3.769798 |
| Portugal, Leiría, Porto de Mós | 21AG21 (1-5) | 39.578466, -8.802222 |
| Spain, Asturias, Nava | 21VV19 (1-5) | 43.357293, -5.448783 |
| Spain, Asturias, Ponga | 22VV19 (1-5) | 43.189776, -5.080974 |
| Spain, León, Crémenes | 23VV19 (1-3) | 42.885759, -5.153301 |
| Spain, La Rioja, Ezcaray | 24AG21 (1-5) | 42.295444, -2.978026 |
| Spain, Zamora, Benavente | 24VV19 (1-5) | 42.007651, -5.658967 |
| Spain, Navarra, Mendaza | 25AG21 (1-5) | 42.681938, -2.268423 |
| Spain, Zamora, Ferreruela | 25VV19 (1-5) | 41.76362, -6.069816 |
| Spain, Burgos, Las Bárcenas de Cirión | 26AG21 (1-5) | 43.096063, -3.158155 |
| Spain, Palencia, Aguilar de Campoo | 26VV19 (1-5) | 42.895, -4.240833333 |
| Spain, Cantabria, Suances | 27AG21 (1-5) | 43.418241, -4.032012 |
| Spain, Gipuzkoa, Eibar | 28AG21 (1-5) | 43.178718, -2.465117 |

| | | |
|---|---|---|
| Spain, Madrid, San Lorenzo de El Escorial | 28VV9 (1-5) | 40.574948, -4.152708 |
| Spain, Navarra, Igantzi | 29AG21 (1-5) | 43.217033, -1.700117 |
| Spain, Navarra, Romanzado | 30AG21 (1-5) | 42.662142, -1.222917 |
| ***Hedera iberica*** | | |
| Spain, Cáceres, Villuercas | 03VV18 (1-5) | 39.620785, -5.439747 |
| Portugal, Setúbal, Arrábida | 04VV18 | 38.50551, -9.149715 |
| Portugal, Setúbal, Arrábida | 05VV18 (1-10) | 38.497361, -9.054198 |
| Portugal, Algarve, Monchique | 08VV18 (1-5) | 37.307021, -8.58707 |
| Portugal, Algarve, Foia peak | 09VV18 (1,2) | 37.314972, -8.591478 |
| Spain, Cádiz, Alcornocales | 10VV18 (1-10) | 36.22511, -5.582609 |
| Spain, Huelva, Fuenteheridos | 11VV18 (1-10) | 37.90834, -6.658308 |
| Portugal, Lisboa, Colares | 22AG21 (1-5) | 38.771511, -9.446594 |
| Portugal, Évora, Nossa Senhora da Boa Fé | 23AG21 (1-5) | 38.55477, -8.102469 |
| ***Hedera maderensis*** | | |
| Portugal, Madeira, São Vicente | 04VV19 (1-5) | 32.729942, -17.030522 |
| Portugal, Madeira, Ponta do Pargo | 06VV19 (1-5) | 32.764073, -17.027144 |
| Portugal, Madeira, Ponta Delgada | 07VV19 (1-3) | 32.825539, -16.981865 |
| Portugal, Madeira, Achadas da Cruz | 08VV19 (1,2) | 32.840347, -17.193568 |
| Portugal, Madeira, Levada Grande | 09VV19 (1-4) | 32.847879, -17.194113 |
| Portugal, Madeira, Prazeres | 10VV19 (1-5) | 32.755229, -17.217884 |
| Portugal, Madeira, São Gonçalo | 11VV19 (1-5) | 32.673849, -16.86322 |
| Portugal, Madeira, Sao Jorge | 12VV19 | 32.827432, -16.902176 |
| Portugal, Madeira, Terras da Fora | 13VV19 (1-4) | 32.823432, -16.934766 |

**Supplementary table 3.** Characteristics of 36 genotyping-by-sequencing datasets used in phylogenetic analyses of the western polyploid clade of *Hedera*. Each dataset is denoted as *c*X*m*Y*p*Z*r*W, with X being the clustering threshold, Y the minimum taxon coverage, Z the ploidy level, and W the level of filtering of individuals (see text). Numbers of filtered loci, sites, single nucleotide polymorphisms (SNPs), phylogenetically informative sites (PIS) and percentage of missing data for each dataset are indicated. *Dataset selected for extensive analysis.

| | # filtered loci | # sites | # SNPs | # PIS | % missing data |
|---|---|---|---|---|---|
| ***c*80*m*4*p*2*r*1** | 22055 | 1854939 | 136025 | 51336 | 81.7 |
| ***c*80*m*4*p*2*r*2** | 21508 | 1808144 | 131711 | 49601 | 80.9 |
| ***c*80*m*4*p*2*r*3** | 20955 | 1761164 | 127993 | 48157 | 80.6 |
| ***c*80*m*4*p*6*r*1** | 30737 | 2602302 | 215125 | 94676 | 74.8 |
| ***c*80*m*4*p*6*r*2** | 30109 | 2547255 | 208530 | 91779 | 74.1 |
| ***c*80*m*4*p*6*r*3** | 29511 | 2496295 | 204200 | 90035 | 73.7 |
| ***c*80*m*15*p*2*r*1** | 5900 | 494976 | 18805 | 8703 | 51.1 |
| ***c*80*m*15*p*2*r*2** | 5664 | 474635 | 17348 | 8000 | 49.2 |
| ***c*80*m*15*p*2*r*3** | 5595 | 468697 | 17121 | 7844 | 49.0 |
| ***c*80*m*15*p*6*r*1** | 11235 | 951544 | 57110 | 32880 | 44.2 |
| ***c*80*m*15*p*6*r*2** | 10881 | 920287 | 53571 | 31046 | 42.6 |
| ***c*80*m*15*p*6*r*3*** | 10799 | 913235 | 53123 | 30793 | 42.4 |
| ***c*85*m*4*p*2*r*1** | 21978 | 1850633 | 101330 | 38327 | 81.1 |
| ***c*85*m*4*p*2*r*2** | 21445 | 1804681 | 97990 | 36942 | 80.4 |
| ***c*85*m*4*p*2*r*3** | 20910 | 1759330 | 95476 | 35952 | 80.1 |
| ***c*85*m*4*p*6*r*1** | 30978 | 2625360 | 175593 | 79495 | 74.1 |
| ***c*85*m*4*p*6*r*2** | 30361 | 2570970 | 170188 | 76998 | 73.4 |
| ***c*85*m*4*p*6*r*3** | 29769 | 2520811 | 167056 | 75685 | 73.0 |
| ***c*85*m*15*p*2*r*1** | 6141 | 515247 | 18170 | 8279 | 51.2 |
| ***c*85*m*15*p*2*r*2** | 5908 | 495068 | 16925 | 7675 | 49.3 |
| ***c*85*m*15*p*2*r*3** | 5832 | 488595 | 16665 | 7504 | 49.1 |
| ***c*85*m*15*p*6*r*1** | 11748 | 995395 | 56941 | 32661 | 44.1 |
| ***c*85*m*15*p*6*r*2** | 11389 | 963773 | 53692 | 30950 | 42.5 |
| ***c*85*m*15*p*6*r*3** | 11297 | 955882 | 53232 | 30673 | 42.3 |
| ***c*90*m*4*p*2*r*1** | 22356 | 1885295 | 73526 | 27690 | 80.3 |
| ***c*90*m*4*p*2*r*2** | 21793 | 1836877 | 70798 | 26623 | 79.5 |
| ***c*90*m*4*p*2*r*3** | 21261 | 1791862 | 69019 | 25938 | 79.2 |
| ***c*90*m*4*p*6*r*1** | 31495 | 2670428 | 138063 | 63483 | 73.5 |
| ***c*90*m*4*p*6*r*2** | 30846 | 2613484 | 133377 | 61435 | 72.7 |
| ***c*90*m*4*p*6*r*3** | 30247 | 2562629 | 130964 | 60392 | 72.4 |
| ***c*90*m*15*p*2*r*1** | 6658 | 559417 | 18232 | 8231 | 51.1 |
| ***c*90*m*15*p*2*r*2** | 6407 | 537801 | 17025 | 7685 | 49.3 |
| ***c*90*m*15*p*2*r*3** | 6324 | 530706 | 16797 | 7544 | 49.0 |
| ***c*90*m*15*p*6*r*1** | 12397 | 1049683 | 54663 | 30941 | 44.3 |
| ***c*90*m*15*p*6*r*2** | 12010 | 1015764 | 51523 | 29370 | 42.7 |
| ***c*90*m*15*p*6*r*3** | 11898 | 1006193 | 51012 | 29043 | 42.5 |

**Supplementary script 1**. R script employed for the niche overlap analyses. Comments between hash marks (###) are informative and are not essential to run the analyses.

```
### Load functions and packages ###

install.packages("ecospat")

install.packages("ade4")

install.packages("adehabitatHR")

install.packages("sp")

install.packages("dplyr")

library(ecospat)

library(ade4)

library(adehabitatHR)

library(sp)

library(dplyr)

library(ggplot2)

### Preparation of datasets ###

### Load climate variable for all site of the study area 1 (column names should be x,y,X1,X2,...,Xn) ###

clim<-na.exclude(read.delim("Variables_Hedera2.txt",h=T,sep=","))

### Loading occurence sites for the species (column names should be x,y) ###

sp<-read.delim("ENSAMBLAJE5.txt",h=T,sep="\t",dec=".")

colnames(sp)[1] <- "Species" ###change the name of the column###

sp$x <- as.numeric(as.character(sp$x)) ### Change to numeric ###

sp$y <- as.numeric(as.character(sp$y))

sp$Species<-factor(sp$Species,levels = c("HIB","IBE","MAD","HEL"))  ###Set the species###

### Sample environmental values for all occurences ###

occ.sp<na.exclude(ecospat.sample.envar(dfsp=sp,colspxy=2:3,colspkept=1:3,dfvar=clim,colvarxy=21:22,colvar=2:20,resolution=0.16666))

occ.sp<-cbind(occ.sp,sp[,1]) ###add species names ###

### List of species###

sp.list<-levels(occ.sp[,1])

sp.nbocc<-c()

for (i in 1:length(sp.list)){sp.nbocc<-c(sp.nbocc,length(which(occ.sp[,1] == sp.list[i])))}
```

```
###Calculate the number of occurrences per species###

sp.list<-sp.list[sp.nbocc>4] ### Remove species with less than 5 occurrences ###

nb.sp<-length(sp.list) ###Number of species###

### Selection of climatic variables to include in the analyses ###

Xvar<-c(2:20)

nvar<-length(Xvar)

### Number of iterations for the tests of equivalency and similarity ###

iterations<-100

#Resolution of the gridding of the climate space

R=100


### PCA ###

data.0<-rbind(occ.sp[,Xvar+2],clim[,Xvar]) ###Dataset for the analysis, includes all the sites of the study area and the occurrences for all the species ###

###Remove autocorrelated variables ###

var.remove.0 <- c("bio01","bio02","bio05","bio06","bio12","bio13","bio14", "bio17", "bio19")

data <- data.0 [, !(colnames(data.0) %in% var.remove.0), drop=FALSE]

w<-c(rep(0,nrow(occ.sp)),rep(1,nrow(clim))) ###Vector of weight, 0 for the occurences, 1 for the sites of the study area###

pca.cal <-dudi.pca(data, row.w = w, center = T, scale = T, scannf = F, nf = 2) # The PCA is calibrated on all the sites of the study area###

### Plot all the the species ###

sc1<- pca.cal$li[,1]

sc2<- pca.cal$li[,2]

### Selection of species ###

sp.choice<- c("HIB","HEL","MAD","IBE") ###choose the set of species for pairwise analyses###

sp.combn<-combn(sp.choice,2)

nsp<-ncol(sp.combn)

overlap<-matrix(nrow=nb.sp,ncol=nb.sp,dimnames = list(sp.choice,sp.choice))  ###Matrix to store overlap values###

equivalency<-matrix(nrow=nb.sp,ncol=nb.sp,dimnames = list(sp.choice,sp.choice))  ###Matrix to store the p-values for sp1-sp2 equivalency tests###
```

```
similarity<-matrix(nrow=nb.sp,ncol=nb.sp,dimnames = list(sp.choice,sp.choice)) ###Matrix to store the p-values for sp2 vs. sp1 similarity tests###

for(i in 1:ncol(sp.combn)) { ###For each combination of species###

row.spa<-which(occ.sp[,1] == sp.combn[1,i]) ###Rows in data corresponding to spa###

row.spb<-which(occ.sp[,1] == sp.combn[2,i]) ### Rows in data corresponding to spb###

name.spa<-sp.combn[1,i]

name.spb<-sp.combn[2,i]

### Predict the scores on the axes###

scores.clim<- pca.cal$li[(nrow(occ.sp)+1):nrow(data),]   ###Scores for global climate###

scores.spa<- pca.cal$li[row.spa,] ###Scores for spa###

scores.spb<- pca.cal$li[row.spb,] ###Scores for spb###

### Calculation of occurrence density and test of niche equivalency and similarity ###

za<- ecospat.grid.clim.dyn(scores.clim,scores.clim,scores.spa,R)

zb<- ecospat.grid.clim.dyn(scores.clim,scores.clim,scores.spb,R)

###Test of niche equivalency and similarity according to Warren et al. 2008###

equ<-ecospat.niche.equivalency.test(za,zb,rep=100, alternative="lower") ### At least 100 for final analyses###

sim<-ecospat.niche.similarity.test(za,zb,rep=100,rand.type = 1, alternative="greater") ### Both za and zb are randomly shifted in the background (previous versions of ecospat implemented rand.type =2) ###


###plot###

pdf(file=paste(name.spa," x ",name.spb,".pdf",sep="")) ### Create a pdf file named from the names of the 2 species###

layout(matrix(c(1,1,2,2,1,1,2,2,3,3,4,5,3,3,6,7), 4, 4, byrow = TRUE))

ecospat.plot.niche(za,title=name.spa,name.axis1="PC1",name.axis2="PC2")

ecospat.plot.niche(zb,title=name.spb,name.axis1="PC1",name.axis2="PC2")

ecospat.plot.contrib(pca.cal$co,pca.cal$eig)

plot.new();text(0.5,0.5,paste("nicheoverlap:","\n","D=",round(as.numeric(ecospat.niche.overlap(za,zb,cor=T)[1]),3)))

plot.new()

ecospat.plot.overlap.test(equ,"D","Equivalency")

ecospat.plot.overlap.test(sim,"D","Similarity")
```

```
dev.off()

overlap[sp.combn[1,i],sp.combn[2,i]]<-ecospat.niche.overlap(za,zb,cor=T)[[1]]  ###Store
overlap value###

equivalency[sp.combn[1,i],sp.combn[2,i]]<-equ$p.D ###Store equivalency value###

similarity[sp.combn[1,i],sp.combn[2,i]]<-sim$p.D ###Store similarity value###}

### Niche position and breadth###

niche<-
matrix(nrow=nb.sp,ncol=4,dimnames=list(sp.choice,c("pos1","breadth1","pos2","breadth2")))
###Matrix to store niche characteristics###

for(i in 1:length(sp.choice)) { ###For each chosen species###

row.sp<-which(occ.sp[,1] == sp.choice[i]) ###Rows in data corresponding to sp###

name.sp<-sp.choice[i]

scores.sp<- pca.cal$li[row.sp,] ###Scores for sp###

### Calculation of occurrence density and test of niche equivalency and similarity ###

z<- ecospat.grid.clim.dyn(scores.clim,scores.clim,scores.sp,R)

c<-sample(1:(R*R),1000,prob=values(z$z.uncor)) ###Indices of 1000 random pixel weighted by
density in the CP1 and CP2 PCA space ###

y=(c%/%R)+1;x=c%%R ### Coordinates of the pixels along CP1 and CP2###

CP.sim<-z$x[x] ###Scores of random pixels on CP1###

niche[i,1]<-median(CP.sim) ### Niche position on CP1###

niche[i,2]<-var(CP.sim) ### Niche breadth on CP1###

CP2.sim<-z$y[y] ### Scores of random pixels on CP2###

niche[i,3]<-median(CP2.sim) ### Niche position on CP2###

niche[i,4]<-var(CP2.sim) ### Niche breadth on CP2###}
```

Supplementary figure 1. Coalescent-based reconstruction from SVDquartets of the three species of western polyploid clade of *Hedera* plus *H. helix*. Partitions correspond to *H. helix*, *H. hibernica*, *H. maderensis* and the five disjunct populations of *H. iberica*. Numbers indicate bootstrap support values.

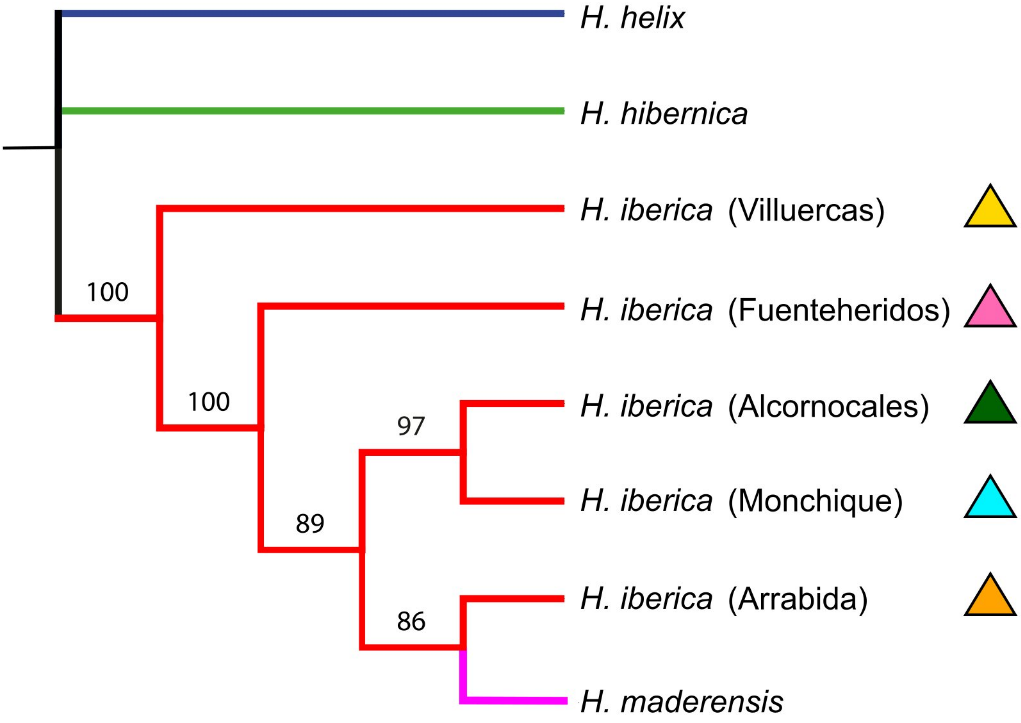

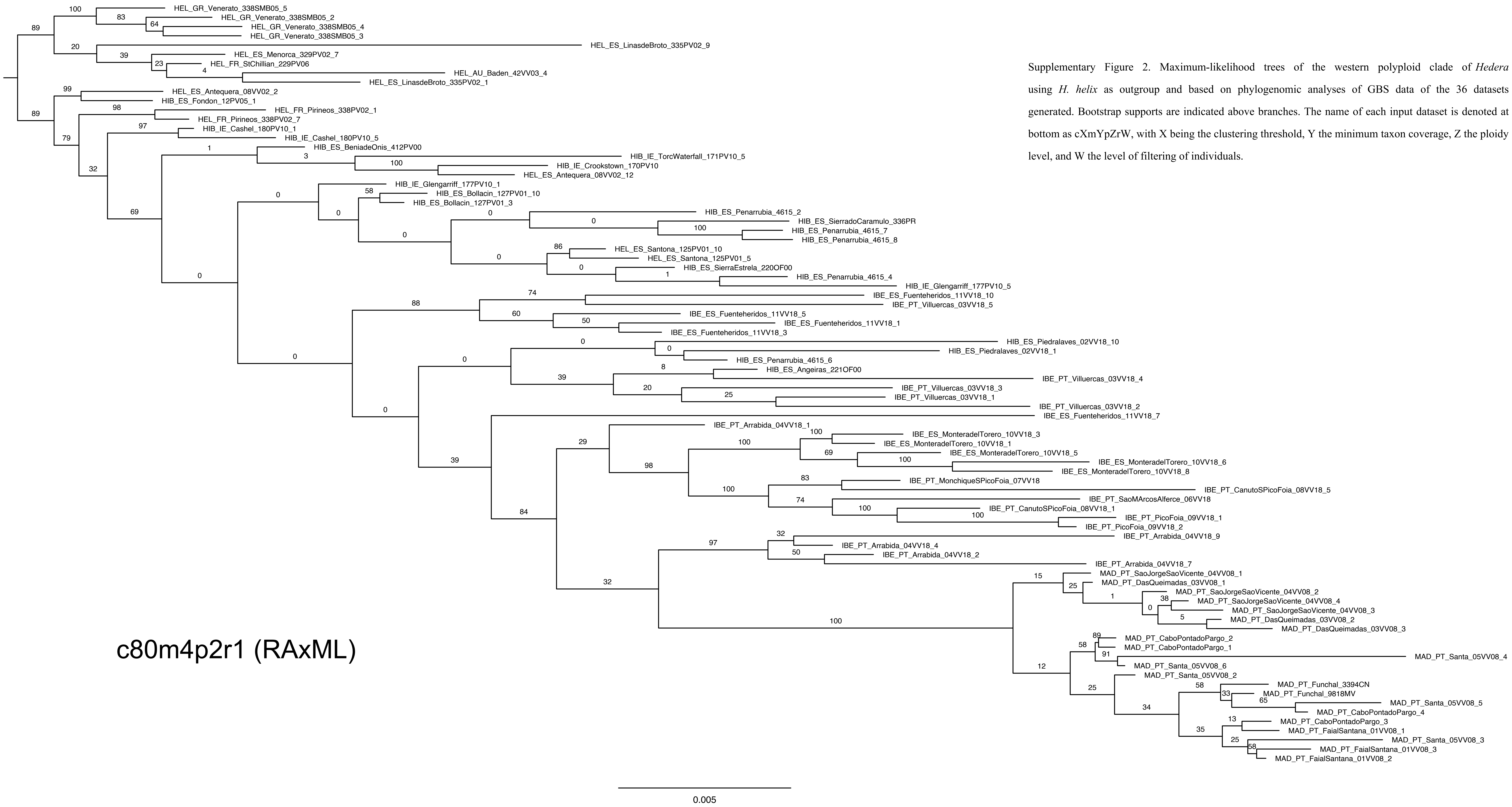


Supplementary Figure 2. Maximum-likelihood trees of the western polyploid clade of *Hedera* using *H. helix* as outgroup and based on phylogenomic analyses of GBS data of the 36 datasets generated. Bootstrap supports are indicated above branches. The name of each input dataset is denoted at bottom as cXmYpZrW, with X being the clustering threshold, Y the minimum taxon coverage, Z the ploidy level, and W the level of filtering of individuals.

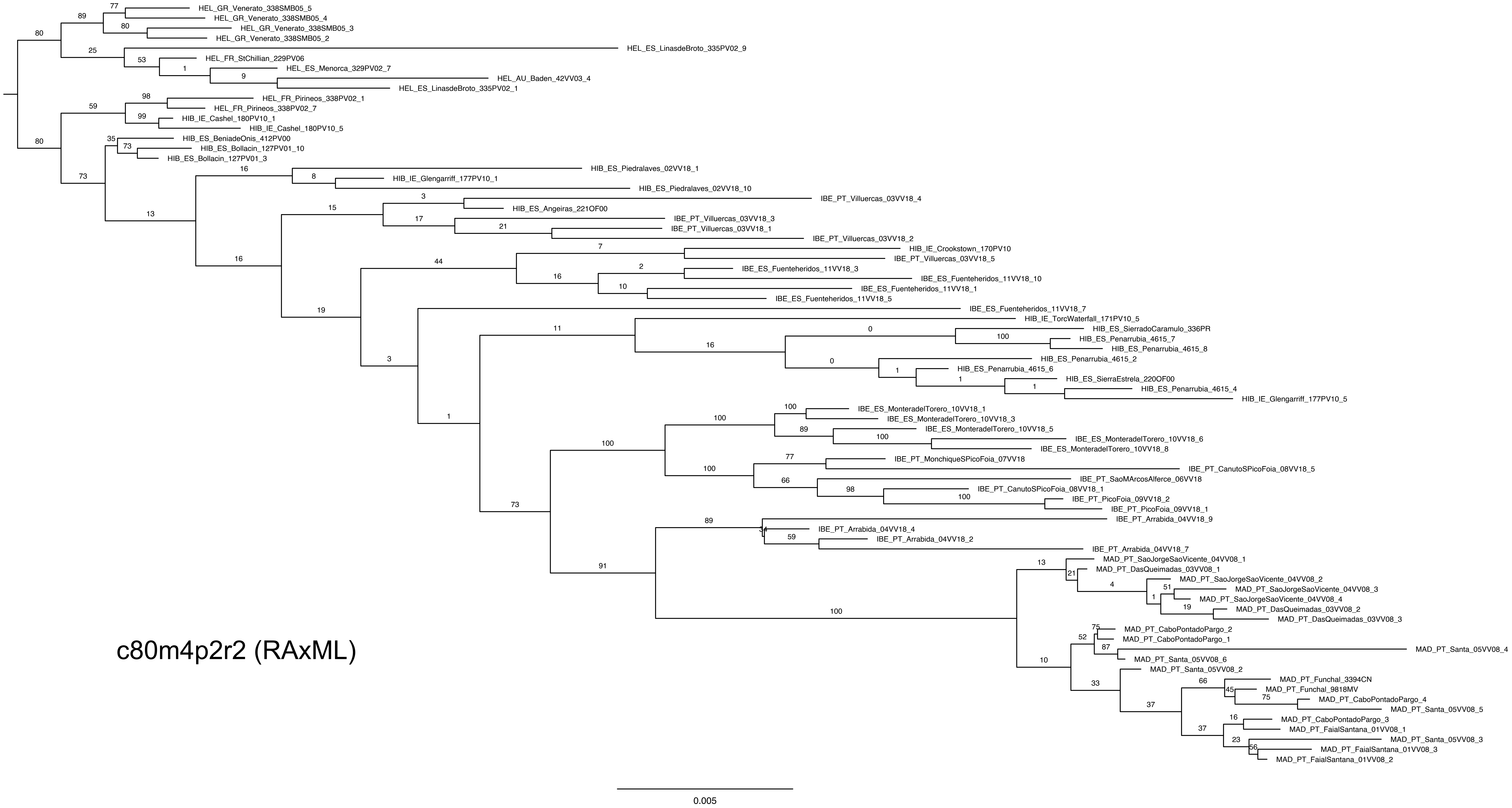
HEL_GR_Venerato_338SMB05_5
HEL_GR_Venerato_338SMB05_4
HEL_GR_Venerato_338SMB05_3
HEL_GR_Venerato_338SMB05_2
HEL_ES_LinasdeBroto_335PV02_9
HEL_FR_StChillian_229PV06
HEL_ES_Menorca_329PV02_7
HEL_AU_Baden_42VV03_4
HEL_ES_LinasdeBroto_335PV02_1
HEL_FR_Pirineos_338PV02_1
HEL_FR_Pirineos_338PV02_7
HIB_IE_Cashel_180PV10_1
HIB_IE_Cashel_180PV10_5
HIB_ES_BeniadeOnis_412PV00
HIB_ES_Bollacin_127PV01_10
HIB_ES_Bollacin_127PV01_3
HIB_ES_Piedralaves_02VV18_1
HIB_IE_Glengarriff_177PV10_1
HIB_ES_Piedralaves_02VV18_10
IBE_PT_Villuercas_03VV18_4
HIB_ES_Angeiras_221OF00
IBE_PT_Villuercas_03VV18_3
IBE_PT_Villuercas_03VV18_1
IBE_PT_Villuercas_03VV18_2
HIB_IE_Crookstown_170PV10
IBE_PT_Villuercas_03VV18_5
IBE_ES_Fuenteheridos_11VV18_3
IBE_ES_Fuenteheridos_11VV18_10
IBE_ES_Fuenteheridos_11VV18_1
IBE_ES_Fuenteheridos_11VV18_5
IBE_ES_Fuenteheridos_11VV18_7
HIB_IE_TorcWaterfall_171PV10_5
HIB_ES_SierradoCaramulo_336PR
HIB_ES_Penarrubia_4615_7
HIB_ES_Penarrubia_4615_8
HIB_ES_Penarrubia_4615_2
HIB_ES_Penarrubia_4615_6
HIB_ES_SierraEstrela_220OF00
HIB_ES_Penarrubia_4615_4
HIB_IE_Glengarriff_177PV10_5
IBE_ES_MonteradelTorero_10VV18_1
IBE_ES_MonteradelTorero_10VV18_3
IBE_ES_MonteradelTorero_10VV18_5
IBE_ES_MonteradelTorero_10VV18_6
IBE_ES_MonteradelTorero_10VV18_8
IBE_PT_MonchiqueSPicoFoia_07VV18
IBE_PT_CanutoSPicoFoia_08VV18_5
IBE_PT_SaoMArcosAlferce_06VV18
IBE_PT_CanutoSPicoFoia_08VV18_1
IBE_PT_PicoFoia_09VV18_2
IBE_PT_PicoFoia_09VV18_1
IBE_PT_Arrabida_04VV18_9
IBE_PT_Arrabida_04VV18_4
IBE_PT_Arrabida_04VV18_2
IBE_PT_Arrabida_04VV18_7
MAD_PT_SaoJorgeSaoVicente_04VV08_1
MAD_PT_DasQueimadas_03VV08_1
MAD_PT_SaoJorgeSaoVicente_04VV08_2
MAD_PT_SaoJorgeSaoVicente_04VV08_3
MAD_PT_SaoJorgeSaoVicente_04VV08_4
MAD_PT_DasQueimadas_03VV08_2
MAD_PT_DasQueimadas_03VV08_3
MAD_PT_CaboPontadoPargo_2
MAD_PT_CaboPontadoPargo_1
MAD_PT_Santa_05VV08_4
MAD_PT_Santa_05VV08_6
MAD_PT_Santa_05VV08_2
MAD_PT_Funchal_3394CN
MAD_PT_Funchal_9818MV
MAD_PT_CaboPontadoPargo_4
MAD_PT_Santa_05VV08_5
MAD_PT_CaboPontadoPargo_3
MAD_PT_FaialSantana_01VV08_1
MAD_PT_Santa_05VV08_3
MAD_PT_FaialSantana_01VV08_3
MAD_PT_FaialSantana_01VV08_2
c80m4p2r2 (RAxML)
0.005

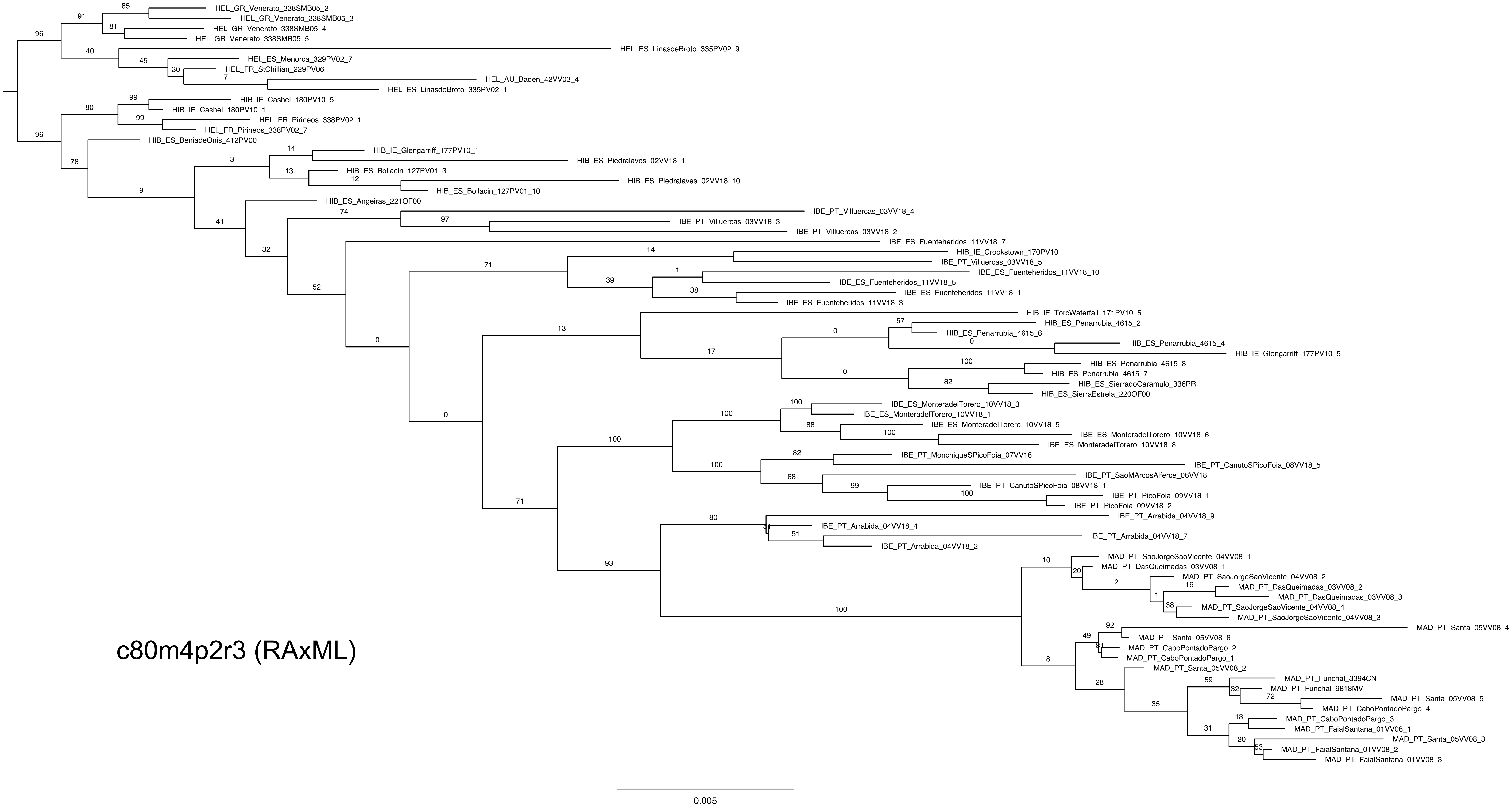


c80m4p2r3 (RAxML)

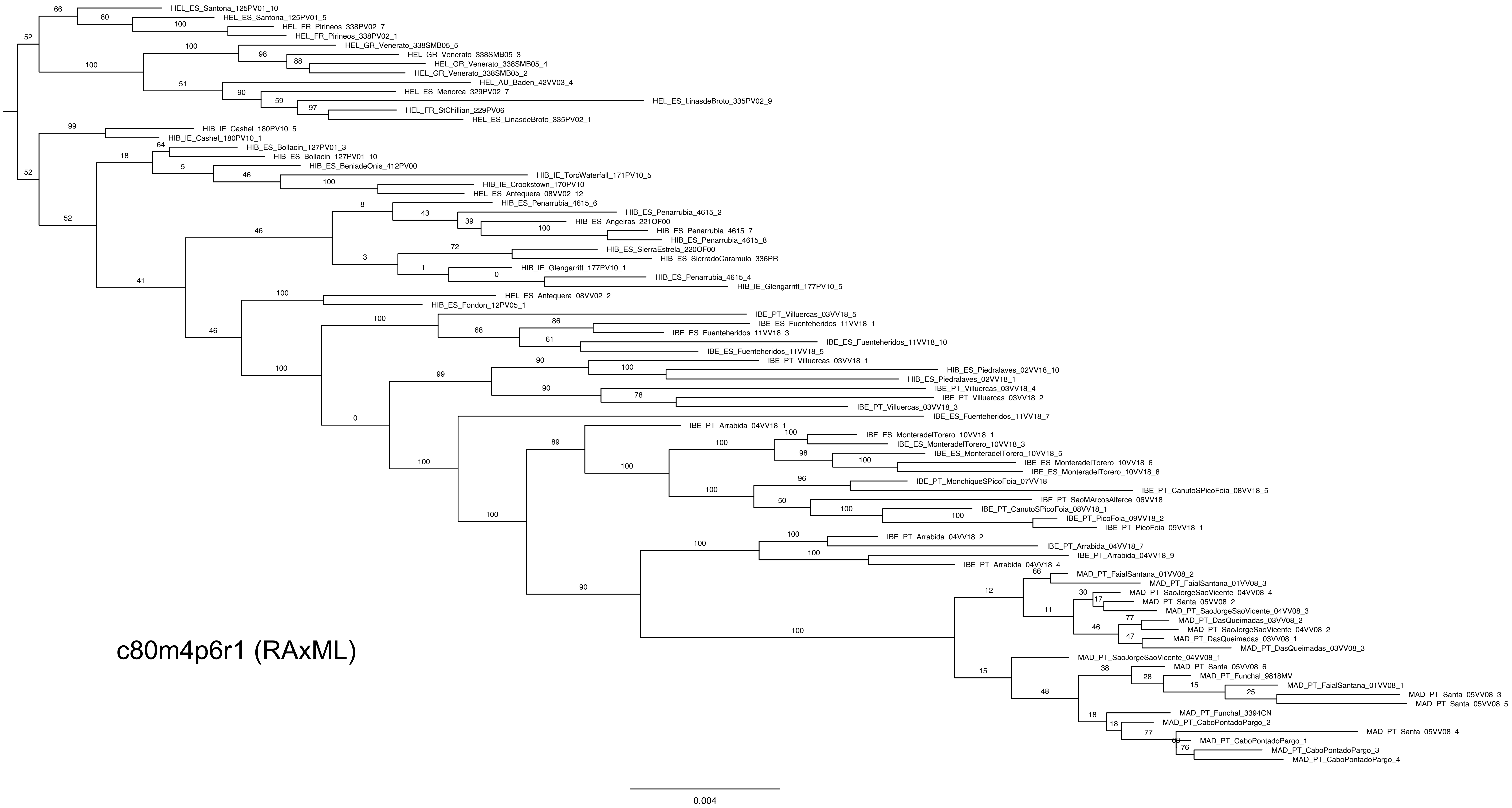
c80m4p6r1 (RAxML)
HEL_ES_Santona_125PV01_10
HEL_ES_Santona_125PV01_5
HEL_FR_Pirineos_338PV02_7
HEL_FR_Pirineos_338PV02_1
HEL_GR_Venerato_338SMB05_5
HEL_GR_Venerato_338SMB05_3
HEL_GR_Venerato_338SMB05_4
HEL_GR_Venerato_338SMB05_2
HEL_AU_Baden_42VV03_4
HEL_ES_Menorca_329PV02_7
HEL_ES_LinasdeBroto_335PV02_9
HEL_FR_StChillian_229PV06
HEL_ES_LinasdeBroto_335PV02_1
HIB_IE_Cashel_180PV10_5
HIB_IE_Cashel_180PV10_1
HIB_ES_Bollacin_127PV01_3
HIB_ES_Bollacin_127PV01_10
HIB_ES_BeniadeOnis_412PV00
HIB_IE_TorcWaterfall_171PV10_5
HIB_IE_Crookstown_170PV10
HEL_ES_Antequera_08VV02_12
HIB_ES_Penarrubia_4615_6
HIB_ES_Penarrubia_4615_2
HIB_ES_Angeiras_221OF00
HIB_ES_Penarrubia_4615_7
HIB_ES_Penarrubia_4615_8
HIB_ES_SierraEstrela_220OF00
HIB_ES_SierradoCaramulo_336PR
HIB_IE_Glengarriff_177PV10_1
HIB_ES_Penarrubia_4615_4
HIB_IE_Glengarriff_177PV10_5
HEL_ES_Antequera_08VV02_2
HIB_ES_Fondon_12PV05_1
IBE_PT_Villuercas_03VV18_5
IBE_ES_Fuenteheridos_11VV18_1
IBE_ES_Fuenteheridos_11VV18_3
IBE_ES_Fuenteheridos_11VV18_10
IBE_ES_Fuenteheridos_11VV18_5
IBE_PT_Villuercas_03VV18_1
HIB_ES_Piedralaves_02VV18_10
HIB_ES_Piedralaves_02VV18_1
IBE_PT_Villuercas_03VV18_4
IBE_PT_Villuercas_03VV18_2
IBE_PT_Villuercas_03VV18_3
IBE_ES_Fuenteheridos_11VV18_7
IBE_PT_Arrabida_04VV18_1
IBE_ES_MonteradelTorero_10VV18_1
IBE_ES_MonteradelTorero_10VV18_3
IBE_ES_MonteradelTorero_10VV18_5
IBE_ES_MonteradelTorero_10VV18_6
IBE_ES_MonteradelTorero_10VV18_8
IBE_PT_MonchiqueSPicoFoia_07VV18
IBE_PT_CanutoSPicoFoia_08VV18_5
IBE_PT_SaoMArcosAlferce_06VV18
IBE_PT_CanutoSPicoFoia_08VV18_1
IBE_PT_PicoFoia_09VV18_2
IBE_PT_PicoFoia_09VV18_1
IBE_PT_Arrabida_04VV18_2
IBE_PT_Arrabida_04VV18_7
IBE_PT_Arrabida_04VV18_9
IBE_PT_Arrabida_04VV18_4
MAD_PT_FaialSantana_01VV08_2
MAD_PT_FaialSantana_01VV08_3
MAD_PT_SaoJorgeSaoVicente_04VV08_4
MAD_PT_Santa_05VV08_2
MAD_PT_SaoJorgeSaoVicente_04VV08_3
MAD_PT_DasQueimadas_03VV08_2
MAD_PT_SaoJorgeSaoVicente_04VV08_2
MAD_PT_DasQueimadas_03VV08_1
MAD_PT_DasQueimadas_03VV08_3
MAD_PT_SaoJorgeSaoVicente_04VV08_1
MAD_PT_Santa_05VV08_6
MAD_PT_Funchal_9818MV
MAD_PT_FaialSantana_01VV08_1
MAD_PT_Santa_05VV08_3
MAD_PT_Santa_05VV08_5
MAD_PT_Funchal_3394CN
MAD_PT_CaboPontadoPargo_2
MAD_PT_Santa_05VV08_4
MAD_PT_CaboPontadoPargo_1
MAD_PT_CaboPontadoPargo_3
MAD_PT_CaboPontadoPargo_4
0.004

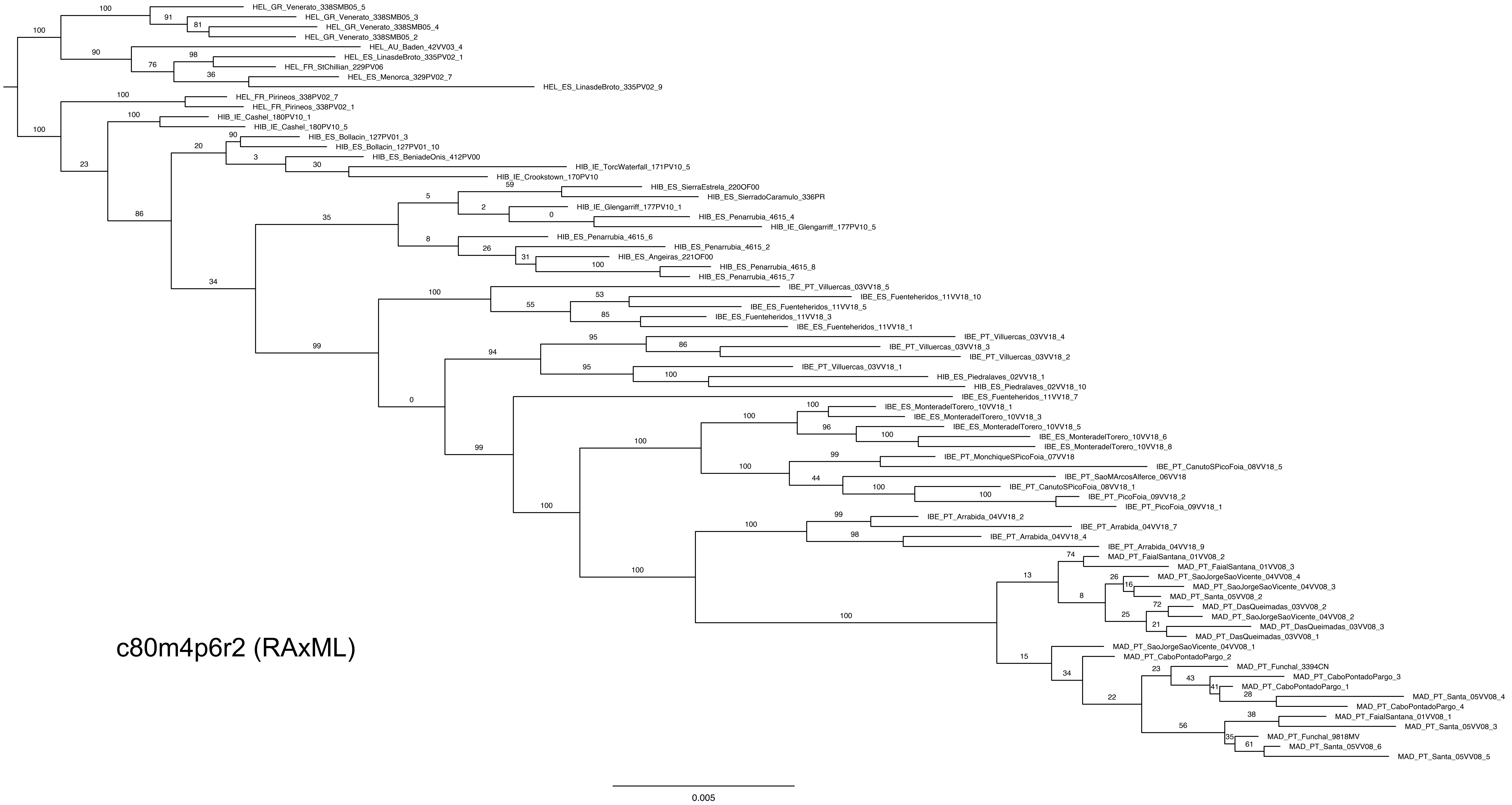
HEL_GR_Venerato_338SMB05_5
HEL_GR_Venerato_338SMB05_3
HEL_GR_Venerato_338SMB05_4
HEL_GR_Venerato_338SMB05_2
HEL_AU_Baden_42VV03_4
HEL_ES_LinasdeBroto_335PV02_1
HEL_FR_StChillian_229PV06
HEL_ES_Menorca_329PV02_7
HEL_ES_LinasdeBroto_335PV02_9
HEL_FR_Pirineos_338PV02_7
HEL_FR_Pirineos_338PV02_1
HIB_IE_Cashel_180PV10_1
HIB_IE_Cashel_180PV10_5
HIB_ES_Bollacin_127PV01_3
HIB_ES_Bollacin_127PV01_10
HIB_ES_BeniadeOnis_412PV00
HIB_IE_TorcWaterfall_171PV10_5
HIB_IE_Crookstown_170PV10
HIB_ES_SierraEstrela_220OF00
HIB_ES_SierradoCaramulo_336PR
HIB_IE_Glengarriff_177PV10_1
HIB_ES_Penarrubia_4615_4
HIB_IE_Glengarriff_177PV10_5
HIB_ES_Penarrubia_4615_6
HIB_ES_Penarrubia_4615_2
HIB_ES_Angeiras_221OF00
HIB_ES_Penarrubia_4615_8
HIB_ES_Penarrubia_4615_7
IBE_PT_Villuercas_03VV18_5
IBE_ES_Fuenteheridos_11VV18_10
IBE_ES_Fuenteheridos_11VV18_5
IBE_ES_Fuenteheridos_11VV18_3
IBE_ES_Fuenteheridos_11VV18_1
IBE_PT_Villuercas_03VV18_4
IBE_PT_Villuercas_03VV18_3
IBE_PT_Villuercas_03VV18_2
IBE_PT_Villuercas_03VV18_1
HIB_ES_Piedralaves_02VV18_1
HIB_ES_Piedralaves_02VV18_10
IBE_ES_Fuenteheridos_11VV18_7
IBE_ES_MonteradelTorero_10VV18_1
IBE_ES_MonteradelTorero_10VV18_3
IBE_ES_MonteradelTorero_10VV18_5
IBE_ES_MonteradelTorero_10VV18_6
IBE_ES_MonteradelTorero_10VV18_8
IBE_PT_MonchiqueSPicoFoia_07VV18
IBE_PT_CanutoSPicoFoia_08VV18_5
IBE_PT_SaoMArcosAlferce_06VV18
IBE_PT_CanutoSPicoFoia_08VV18_1
IBE_PT_PicoFoia_09VV18_2
IBE_PT_PicoFoia_09VV18_1
IBE_PT_Arrabida_04VV18_2
IBE_PT_Arrabida_04VV18_7
IBE_PT_Arrabida_04VV18_4
IBE_PT_Arrabida_04VV18_9
MAD_PT_FaialSantana_01VV08_2
MAD_PT_FaialSantana_01VV08_3
MAD_PT_SaoJorgeSaoVicente_04VV08_4
MAD_PT_SaoJorgeSaoVicente_04VV08_3
MAD_PT_Santa_05VV08_2
MAD_PT_DasQueimadas_03VV08_2
MAD_PT_SaoJorgeSaoVicente_04VV08_2
MAD_PT_DasQueimadas_03VV08_3
MAD_PT_DasQueimadas_03VV08_1
MAD_PT_SaoJorgeSaoVicente_04VV08_1
MAD_PT_CaboPontadoPargo_2
MAD_PT_Funchal_3394CN
MAD_PT_CaboPontadoPargo_3
MAD_PT_CaboPontadoPargo_1
MAD_PT_Santa_05VV08_4
MAD_PT_CaboPontadoPargo_4
MAD_PT_FaialSantana_01VV08_1
MAD_PT_Santa_05VV08_3
MAD_PT_Funchal_9818MV
MAD_PT_Santa_05VV08_6
MAD_PT_Santa_05VV08_5
c80m4p6r2 (RAxML)
0.005

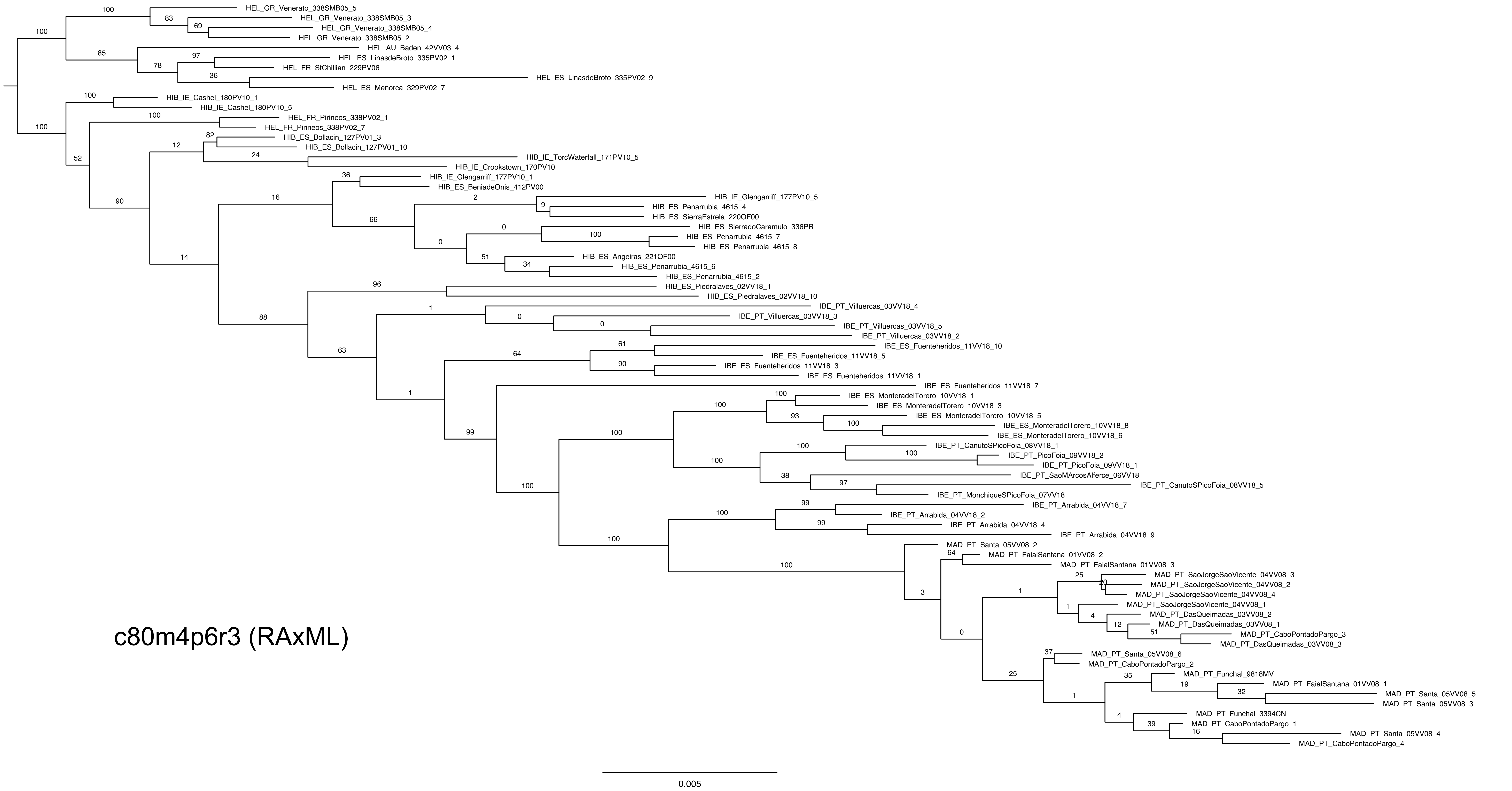
HEL_GR_Venerato_338SMB05_5
HEL_GR_Venerato_338SMB05_3
HEL_GR_Venerato_338SMB05_4
HEL_GR_Venerato_338SMB05_2
HEL_AU_Baden_42VV03_4
HEL_ES_LinasdeBroto_335PV02_1
HEL_FR_StChillian_229PV06
HEL_ES_LinasdeBroto_335PV02_9
HEL_ES_Menorca_329PV02_7
HIB_IE_Cashel_180PV10_1
HIB_IE_Cashel_180PV10_5
HEL_FR_Pirineos_338PV02_1
HEL_FR_Pirineos_338PV02_7
HIB_ES_Bollacin_127PV01_3
HIB_ES_Bollacin_127PV01_10
HIB_IE_TorcWaterfall_171PV10_5
HIB_IE_Crookstown_170PV10
HIB_IE_Glengarriff_177PV10_1
HIB_ES_BeniadeOnis_412PV00
HIB_IE_Glengarriff_177PV10_5
HIB_ES_Penarrubia_4615_4
HIB_ES_SierraEstrela_220OF00
HIB_ES_SierradoCaramulo_336PR
HIB_ES_Penarrubia_4615_7
HIB_ES_Penarrubia_4615_8
HIB_ES_Angeiras_221OF00
HIB_ES_Penarrubia_4615_6
HIB_ES_Penarrubia_4615_2
HIB_ES_Piedralaves_02VV18_1
HIB_ES_Piedralaves_02VV18_10
IBE_PT_Villuercas_03VV18_4
IBE_PT_Villuercas_03VV18_3
IBE_PT_Villuercas_03VV18_5
IBE_PT_Villuercas_03VV18_2
IBE_ES_Fuenteheridos_11VV18_10
IBE_ES_Fuenteheridos_11VV18_5
IBE_ES_Fuenteheridos_11VV18_3
IBE_ES_Fuenteheridos_11VV18_1
IBE_ES_Fuenteheridos_11VV18_7
IBE_ES_MonteradelTorero_10VV18_1
IBE_ES_MonteradelTorero_10VV18_3
IBE_ES_MonteradelTorero_10VV18_5
IBE_ES_MonteradelTorero_10VV18_8
IBE_ES_MonteradelTorero_10VV18_6
IBE_PT_CanutoSPicoFoia_08VV18_1
IBE_PT_PicoFoia_09VV18_2
IBE_PT_PicoFoia_09VV18_1
IBE_PT_SaoMArcosAlferce_06VV18
IBE_PT_CanutoSPicoFoia_08VV18_5
IBE_PT_MonchiqueSPicoFoia_07VV18
IBE_PT_Arrabida_04VV18_7
IBE_PT_Arrabida_04VV18_2
IBE_PT_Arrabida_04VV18_4
IBE_PT_Arrabida_04VV18_9
MAD_PT_Santa_05VV08_2
MAD_PT_FaialSantana_01VV08_2
MAD_PT_FaialSantana_01VV08_3
MAD_PT_SaoJorgeSaoVicente_04VV08_3
MAD_PT_SaoJorgeSaoVicente_04VV08_2
MAD_PT_SaoJorgeSaoVicente_04VV08_4
MAD_PT_SaoJorgeSaoVicente_04VV08_1
MAD_PT_DasQueimadas_03VV08_2
MAD_PT_DasQueimadas_03VV08_1
MAD_PT_CaboPontadoPargo_3
MAD_PT_DasQueimadas_03VV08_3
MAD_PT_Santa_05VV08_6
MAD_PT_CaboPontadoPargo_2
MAD_PT_Funchal_9818MV
MAD_PT_FaialSantana_01VV08_1
MAD_PT_Santa_05VV08_5
MAD_PT_Santa_05VV08_3
MAD_PT_Funchal_3394CN
MAD_PT_CaboPontadoPargo_1
MAD_PT_Santa_05VV08_4
MAD_PT_CaboPontadoPargo_4
c80m4p6r3 (RAxML)
0.005

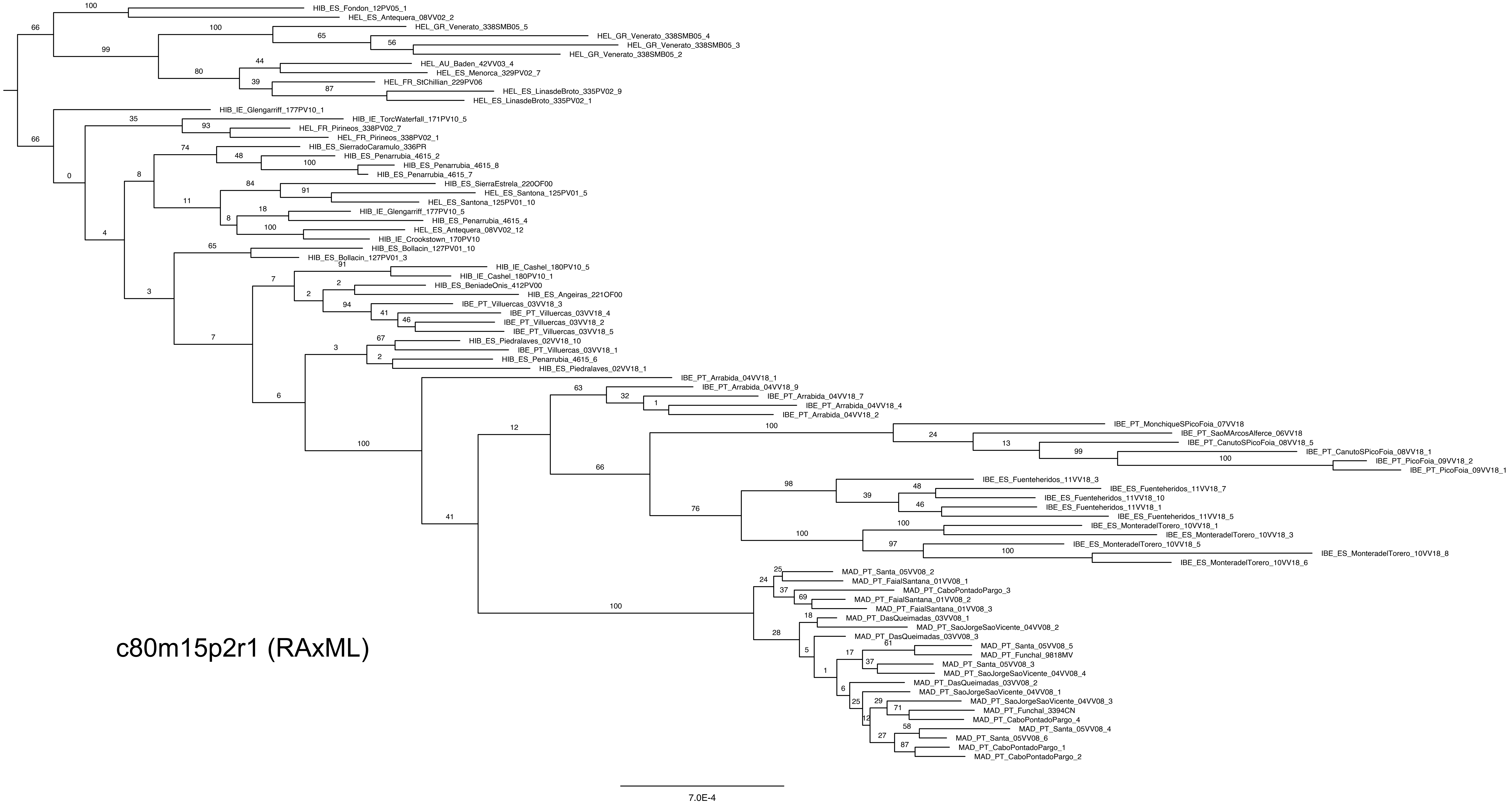
c80m15p2r1 (RAxML)
HIB_ES_Fondon_12PV05_1
HEL_ES_Antequera_08VV02_2
HEL_GR_Venerato_338SMB05_5
HEL_GR_Venerato_338SMB05_4
HEL_GR_Venerato_338SMB05_3
HEL_GR_Venerato_338SMB05_2
HEL_AU_Baden_42VV03_4
HEL_ES_Menorca_329PV02_7
HEL_FR_StChillian_229PV06
HEL_ES_LinasdeBroto_335PV02_9
HEL_ES_LinasdeBroto_335PV02_1
HIB_IE_Glengarriff_177PV10_1
HIB_IE_TorcWaterfall_171PV10_5
HEL_FR_Pirineos_338PV02_7
HEL_FR_Pirineos_338PV02_1
HIB_ES_SierradoCaramulo_336PR
HIB_ES_Penarrubia_4615_2
HIB_ES_Penarrubia_4615_8
HIB_ES_Penarrubia_4615_7
HIB_ES_SierraEstrela_220OF00
HEL_ES_Santona_125PV01_5
HEL_ES_Santona_125PV01_10
HIB_IE_Glengarriff_177PV10_5
HIB_ES_Penarrubia_4615_4
HEL_ES_Antequera_08VV02_12
HIB_IE_Crookstown_170PV10
HIB_ES_Bollacin_127PV01_10
HIB_ES_Bollacin_127PV01_3
HIB_IE_Cashel_180PV10_5
HIB_IE_Cashel_180PV10_1
HIB_ES_BeniadeOnis_412PV00
HIB_ES_Angeiras_221OF00
IBE_PT_Villuercas_03VV18_3
IBE_PT_Villuercas_03VV18_4
IBE_PT_Villuercas_03VV18_2
IBE_PT_Villuercas_03VV18_5
HIB_ES_Piedralaves_02VV18_10
IBE_PT_Villuercas_03VV18_1
HIB_ES_Penarrubia_4615_6
HIB_ES_Piedralaves_02VV18_1
IBE_PT_Arrabida_04VV18_1
IBE_PT_Arrabida_04VV18_9
IBE_PT_Arrabida_04VV18_7
IBE_PT_Arrabida_04VV18_4
IBE_PT_Arrabida_04VV18_2
IBE_PT_MonchiqueSPicoFoia_07VV18
IBE_PT_SaoMArcosAlferce_06VV18
IBE_PT_CanutoSPicoFoia_08VV18_5
IBE_PT_CanutoSPicoFoia_08VV18_1
IBE_PT_PicoFoia_09VV18_2
IBE_PT_PicoFoia_09VV18_1
IBE_ES_Fuenteheridos_11VV18_3
IBE_ES_Fuenteheridos_11VV18_7
IBE_ES_Fuenteheridos_11VV18_10
IBE_ES_Fuenteheridos_11VV18_1
IBE_ES_Fuenteheridos_11VV18_5
IBE_ES_MonteradelTorero_10VV18_1
IBE_ES_MonteradelTorero_10VV18_3
IBE_ES_MonteradelTorero_10VV18_5
IBE_ES_MonteradelTorero_10VV18_8
IBE_ES_MonteradelTorero_10VV18_6
MAD_PT_Santa_05VV08_2
MAD_PT_FaialSantana_01VV08_1
MAD_PT_CaboPontadoPargo_3
MAD_PT_FaialSantana_01VV08_2
MAD_PT_FaialSantana_01VV08_3
MAD_PT_DasQueimadas_03VV08_1
MAD_PT_SaoJorgeSaoVicente_04VV08_2
MAD_PT_DasQueimadas_03VV08_3
MAD_PT_Santa_05VV08_5
MAD_PT_Funchal_9818MV
MAD_PT_Santa_05VV08_3
MAD_PT_SaoJorgeSaoVicente_04VV08_4
MAD_PT_DasQueimadas_03VV08_2
MAD_PT_SaoJorgeSaoVicente_04VV08_1
MAD_PT_SaoJorgeSaoVicente_04VV08_3
MAD_PT_Funchal_3394CN
MAD_PT_CaboPontadoPargo_4
MAD_PT_Santa_05VV08_4
MAD_PT_Santa_05VV08_6
MAD_PT_CaboPontadoPargo_1
MAD_PT_CaboPontadoPargo_2
7.0E-4

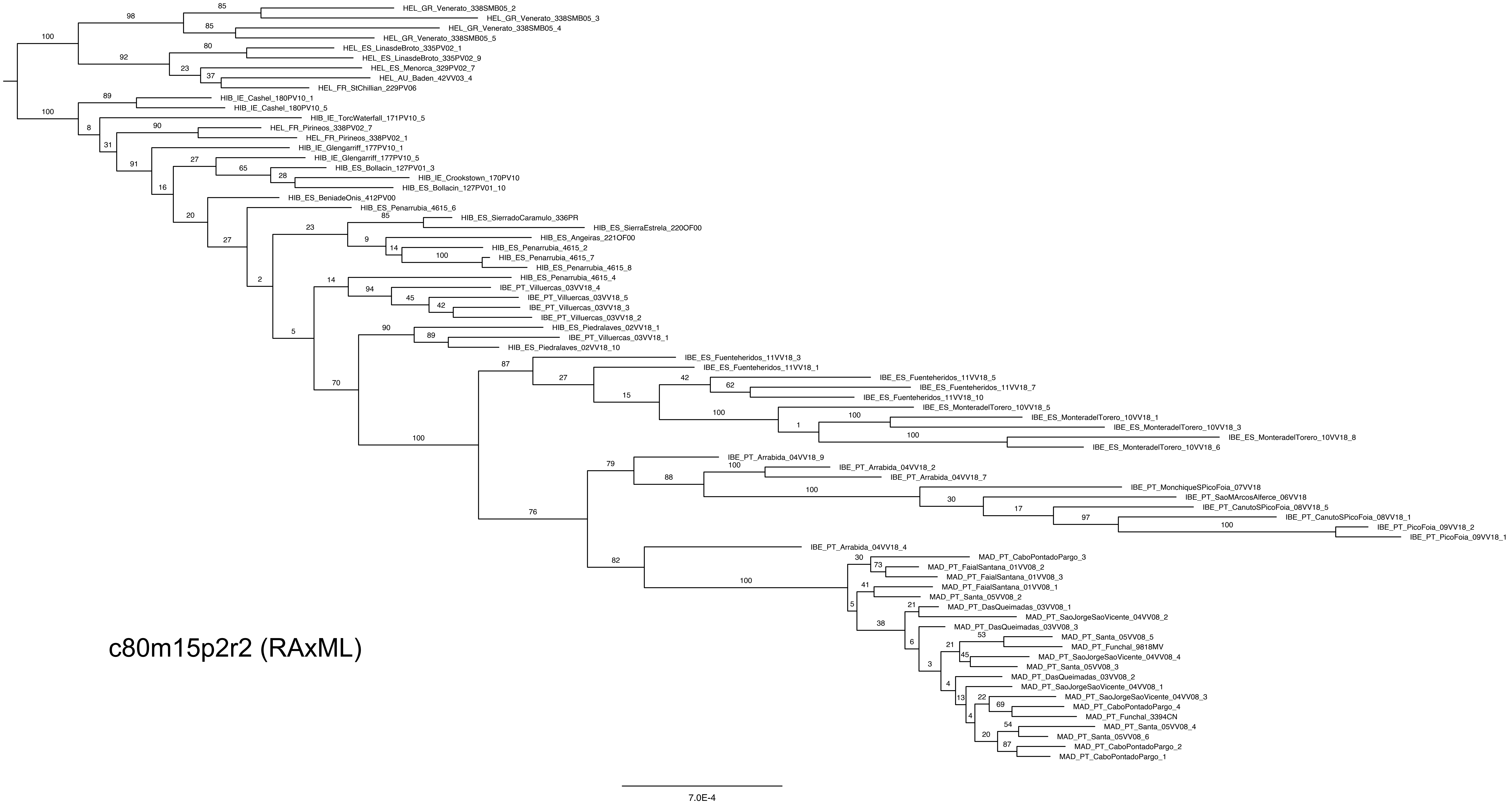

HEL_GR_Venerato_338SMB05_2
HEL_GR_Venerato_338SMB05_3
HEL_GR_Venerato_338SMB05_4
HEL_GR_Venerato_338SMB05_5
HEL_ES_LinasdeBroto_335PV02_1
HEL_ES_LinasdeBroto_335PV02_9
HEL_ES_Menorca_329PV02_7
HEL_AU_Baden_42VV03_4
HEL_FR_StChillian_229PV06
HIB_IE_Cashel_180PV10_1
HIB_IE_Cashel_180PV10_5
HIB_IE_TorcWaterfall_171PV10_5
HEL_FR_Pirineos_338PV02_7
HEL_FR_Pirineos_338PV02_1
HIB_IE_Glengarriff_177PV10_1
HIB_IE_Glengarriff_177PV10_5
HIB_ES_Bollacin_127PV01_3
HIB_IE_Crookstown_170PV10
HIB_ES_Bollacin_127PV01_10
HIB_ES_BeniadeOnis_412PV00
HIB_ES_Penarrubia_4615_6
HIB_ES_SierradoCaramulo_336PR
HIB_ES_SierraEstrela_220OF00
HIB_ES_Angeiras_221OF00
HIB_ES_Penarrubia_4615_2
HIB_ES_Penarrubia_4615_7
HIB_ES_Penarrubia_4615_8
HIB_ES_Penarrubia_4615_4
IBE_PT_Villuercas_03VV18_4
IBE_PT_Villuercas_03VV18_5
IBE_PT_Villuercas_03VV18_3
IBE_PT_Villuercas_03VV18_2
HIB_ES_Piedralaves_02VV18_1
IBE_PT_Villuercas_03VV18_1
HIB_ES_Piedralaves_02VV18_10
IBE_ES_Fuenteheridos_11VV18_3
IBE_ES_Fuenteheridos_11VV18_1
IBE_ES_Fuenteheridos_11VV18_5
IBE_ES_Fuenteheridos_11VV18_7
IBE_ES_Fuenteheridos_11VV18_10
IBE_ES_MonteradelTorero_10VV18_5
IBE_ES_MonteradelTorero_10VV18_1
IBE_ES_MonteradelTorero_10VV18_3
IBE_ES_MonteradelTorero_10VV18_8
IBE_ES_MonteradelTorero_10VV18_6
IBE_PT_Arrabida_04VV18_9
IBE_PT_Arrabida_04VV18_2
IBE_PT_Arrabida_04VV18_7
IBE_PT_MonchiqueSPicoFoia_07VV18
IBE_PT_SaoMArcosAlferce_06VV18
IBE_PT_CanutoSPicoFoia_08VV18_5
IBE_PT_CanutoSPicoFoia_08VV18_1
IBE_PT_PicoFoia_09VV18_2
IBE_PT_PicoFoia_09VV18_1
IBE_PT_Arrabida_04VV18_4
MAD_PT_CaboPontadoPargo_3
MAD_PT_FaialSantana_01VV08_2
MAD_PT_FaialSantana_01VV08_3
MAD_PT_FaialSantana_01VV08_1
MAD_PT_Santa_05VV08_2
MAD_PT_DasQueimadas_03VV08_1
MAD_PT_SaoJorgeSaoVicente_04VV08_2
MAD_PT_DasQueimadas_03VV08_3
MAD_PT_Santa_05VV08_5
MAD_PT_Funchal_9818MV
MAD_PT_SaoJorgeSaoVicente_04VV08_4
MAD_PT_Santa_05VV08_3
MAD_PT_DasQueimadas_03VV08_2
MAD_PT_SaoJorgeSaoVicente_04VV08_1
MAD_PT_SaoJorgeSaoVicente_04VV08_3
MAD_PT_CaboPontadoPargo_4
MAD_PT_Funchal_3394CN
MAD_PT_Santa_05VV08_4
MAD_PT_Santa_05VV08_6
MAD_PT_CaboPontadoPargo_2
MAD_PT_CaboPontadoPargo_1
c80m15p2r2 (RAxML)
7.0E-4

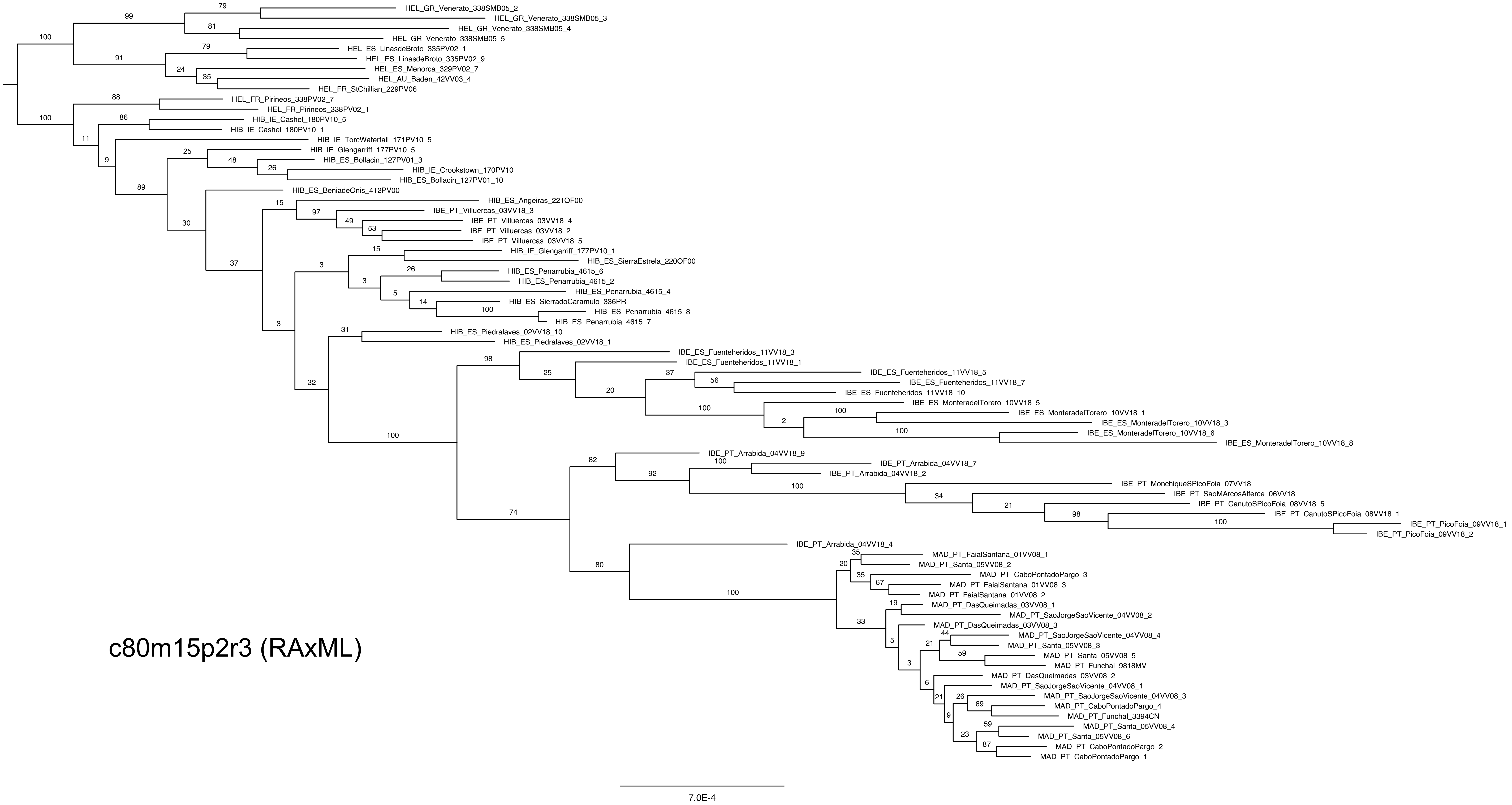
HEL_GR_Venerato_338SMB05_2
HEL_GR_Venerato_338SMB05_3
HEL_GR_Venerato_338SMB05_4
HEL_GR_Venerato_338SMB05_5
HEL_ES_LinasdeBroto_335PV02_1
HEL_ES_LinasdeBroto_335PV02_9
HEL_ES_Menorca_329PV02_7
HEL_AU_Baden_42VV03_4
HEL_FR_StChillian_229PV06
HEL_FR_Pirineos_338PV02_7
HEL_FR_Pirineos_338PV02_1
HIB_IE_Cashel_180PV10_5
HIB_IE_Cashel_180PV10_1
HIB_IE_TorcWaterfall_171PV10_5
HIB_IE_Glengarriff_177PV10_5
HIB_ES_Bollacin_127PV01_3
HIB_IE_Crookstown_170PV10
HIB_ES_Bollacin_127PV01_10
HIB_ES_BeniadeOnis_412PV00
HIB_ES_Angeiras_221OF00
IBE_PT_Villuercas_03VV18_3
IBE_PT_Villuercas_03VV18_4
IBE_PT_Villuercas_03VV18_2
IBE_PT_Villuercas_03VV18_5
HIB_IE_Glengarriff_177PV10_1
HIB_ES_SierraEstrela_220OF00
HIB_ES_Penarrubia_4615_6
HIB_ES_Penarrubia_4615_2
HIB_ES_Penarrubia_4615_4
HIB_ES_SierradoCaramulo_336PR
HIB_ES_Penarrubia_4615_8
HIB_ES_Penarrubia_4615_7
HIB_ES_Piedralaves_02VV18_10
HIB_ES_Piedralaves_02VV18_1
IBE_ES_Fuenteheridos_11VV18_3
IBE_ES_Fuenteheridos_11VV18_1
IBE_ES_Fuenteheridos_11VV18_5
IBE_ES_Fuenteheridos_11VV18_7
IBE_ES_Fuenteheridos_11VV18_10
IBE_ES_MonteradelTorero_10VV18_5
IBE_ES_MonteradelTorero_10VV18_1
IBE_ES_MonteradelTorero_10VV18_3
IBE_ES_MonteradelTorero_10VV18_6
IBE_ES_MonteradelTorero_10VV18_8
IBE_PT_Arrabida_04VV18_9
IBE_PT_Arrabida_04VV18_7
IBE_PT_Arrabida_04VV18_2
IBE_PT_MonchiqueSPicoFoia_07VV18
IBE_PT_SaoMArcosAlferce_06VV18
IBE_PT_CanutoSPicoFoia_08VV18_5
IBE_PT_CanutoSPicoFoia_08VV18_1
IBE_PT_PicoFoia_09VV18_1
IBE_PT_PicoFoia_09VV18_2
IBE_PT_Arrabida_04VV18_4
MAD_PT_FaialSantana_01VV08_1
MAD_PT_Santa_05VV08_2
MAD_PT_CaboPontadoPargo_3
MAD_PT_FaialSantana_01VV08_3
MAD_PT_FaialSantana_01VV08_2
MAD_PT_DasQueimadas_03VV08_1
MAD_PT_SaoJorgeSaoVicente_04VV08_2
MAD_PT_DasQueimadas_03VV08_3
MAD_PT_SaoJorgeSaoVicente_04VV08_4
MAD_PT_Santa_05VV08_3
MAD_PT_Santa_05VV08_5
MAD_PT_Funchal_9818MV
MAD_PT_DasQueimadas_03VV08_2
MAD_PT_SaoJorgeSaoVicente_04VV08_1
MAD_PT_SaoJorgeSaoVicente_04VV08_3
MAD_PT_CaboPontadoPargo_4
MAD_PT_Funchal_3394CN
MAD_PT_Santa_05VV08_4
MAD_PT_Santa_05VV08_6
MAD_PT_CaboPontadoPargo_2
MAD_PT_CaboPontadoPargo_1
c80m15p2r3 (RAxML)
7.0E-4

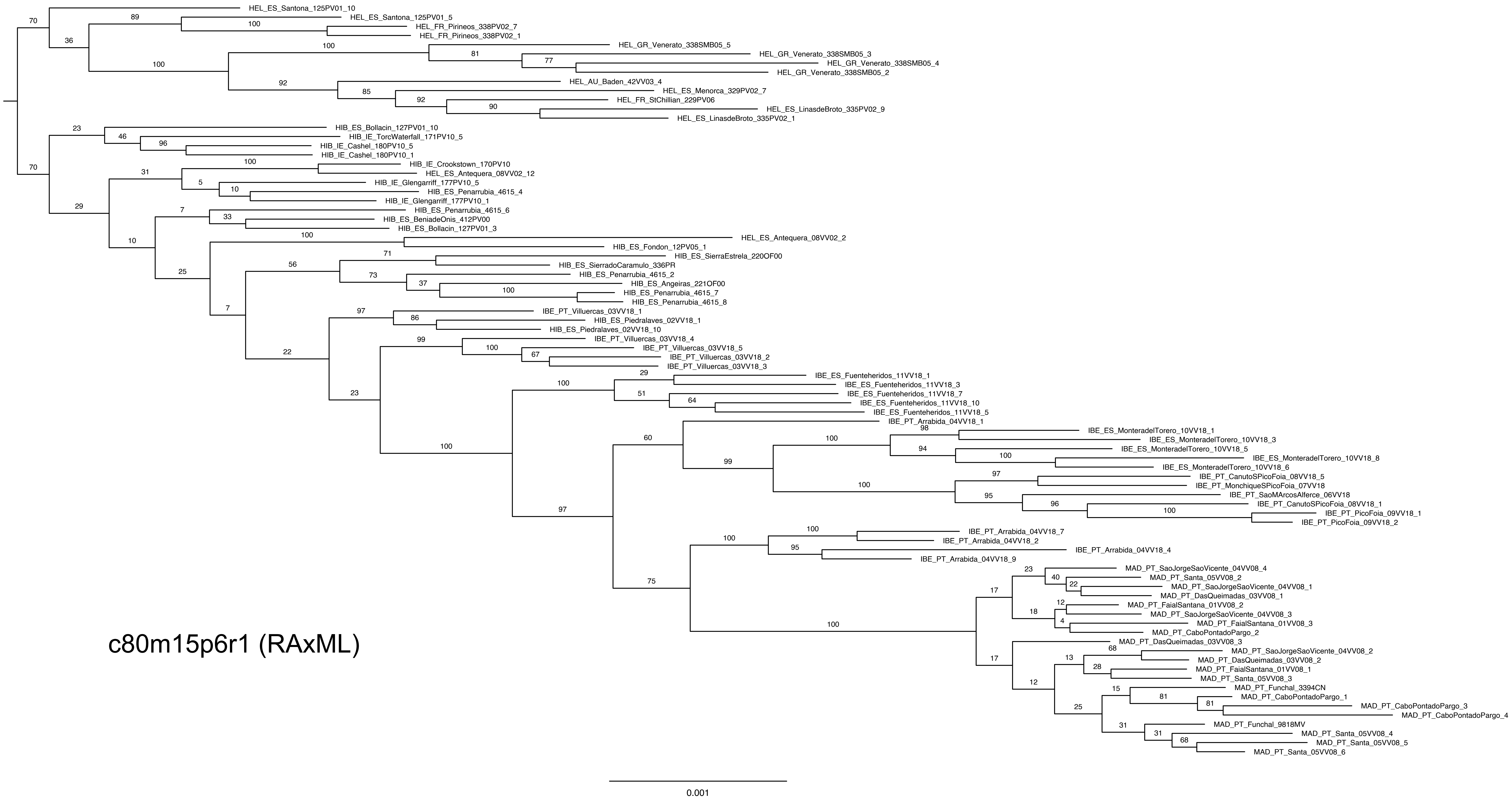
HEL_ES_Santona_125PV01_10
HEL_ES_Santona_125PV01_5
HEL_FR_Pirineos_338PV02_7
HEL_FR_Pirineos_338PV02_1
HEL_GR_Venerato_338SMB05_5
HEL_GR_Venerato_338SMB05_3
HEL_GR_Venerato_338SMB05_4
HEL_GR_Venerato_338SMB05_2
HEL_AU_Baden_42VV03_4
HEL_ES_Menorca_329PV02_7
HEL_FR_StChillian_229PV06
HEL_ES_LinasdeBroto_335PV02_9
HEL_ES_LinasdeBroto_335PV02_1
HIB_ES_Bollacin_127PV01_10
HIB_IE_TorcWaterfall_171PV10_5
HIB_IE_Cashel_180PV10_5
HIB_IE_Cashel_180PV10_1
HIB_IE_Crookstown_170PV10
HEL_ES_Antequera_08VV02_12
HIB_IE_Glengarriff_177PV10_5
HIB_ES_Penarrubia_4615_4
HIB_IE_Glengarriff_177PV10_1
HIB_ES_Penarrubia_4615_6
HIB_ES_BeniadeOnis_412PV00
HIB_ES_Bollacin_127PV01_3
HEL_ES_Antequera_08VV02_2
HIB_ES_Fondon_12PV05_1
HIB_ES_SierraEstrela_220OF00
HIB_ES_SierradoCaramulo_336PR
HIB_ES_Penarrubia_4615_2
HIB_ES_Angeiras_221OF00
HIB_ES_Penarrubia_4615_7
HIB_ES_Penarrubia_4615_8
IBE_PT_Villuercas_03VV18_1
HIB_ES_Piedralaves_02VV18_1
HIB_ES_Piedralaves_02VV18_10
IBE_PT_Villuercas_03VV18_4
IBE_PT_Villuercas_03VV18_5
IBE_PT_Villuercas_03VV18_2
IBE_PT_Villuercas_03VV18_3
IBE_ES_Fuenteheridos_11VV18_1
IBE_ES_Fuenteheridos_11VV18_3
IBE_ES_Fuenteheridos_11VV18_7
IBE_ES_Fuenteheridos_11VV18_10
IBE_ES_Fuenteheridos_11VV18_5
IBE_PT_Arrabida_04VV18_1
IBE_ES_MonteradelTorero_10VV18_1
IBE_ES_MonteradelTorero_10VV18_3
IBE_ES_MonteradelTorero_10VV18_5
IBE_ES_MonteradelTorero_10VV18_8
IBE_ES_MonteradelTorero_10VV18_6
IBE_PT_CanutoSPicoFoia_08VV18_5
IBE_PT_MonchiqueSPicoFoia_07VV18
IBE_PT_SaoMArcosAlferce_06VV18
IBE_PT_CanutoSPicoFoia_08VV18_1
IBE_PT_PicoFoia_09VV18_1
IBE_PT_PicoFoia_09VV18_2
IBE_PT_Arrabida_04VV18_7
IBE_PT_Arrabida_04VV18_2
IBE_PT_Arrabida_04VV18_4
IBE_PT_Arrabida_04VV18_9
MAD_PT_SaoJorgeSaoVicente_04VV08_4
MAD_PT_Santa_05VV08_2
MAD_PT_SaoJorgeSaoVicente_04VV08_1
MAD_PT_DasQueimadas_03VV08_1
MAD_PT_FaialSantana_01VV08_2
MAD_PT_SaoJorgeSaoVicente_04VV08_3
MAD_PT_FaialSantana_01VV08_3
MAD_PT_CaboPontadoPargo_2
MAD_PT_DasQueimadas_03VV08_3
MAD_PT_SaoJorgeSaoVicente_04VV08_2
MAD_PT_DasQueimadas_03VV08_2
MAD_PT_FaialSantana_01VV08_1
MAD_PT_Santa_05VV08_3
MAD_PT_Funchal_3394CN
MAD_PT_CaboPontadoPargo_1
MAD_PT_CaboPontadoPargo_3
MAD_PT_CaboPontadoPargo_4
MAD_PT_Funchal_9818MV
MAD_PT_Santa_05VV08_4
MAD_PT_Santa_05VV08_5
MAD_PT_Santa_05VV08_6
c80m15p6r1 (RAxML)
0.001

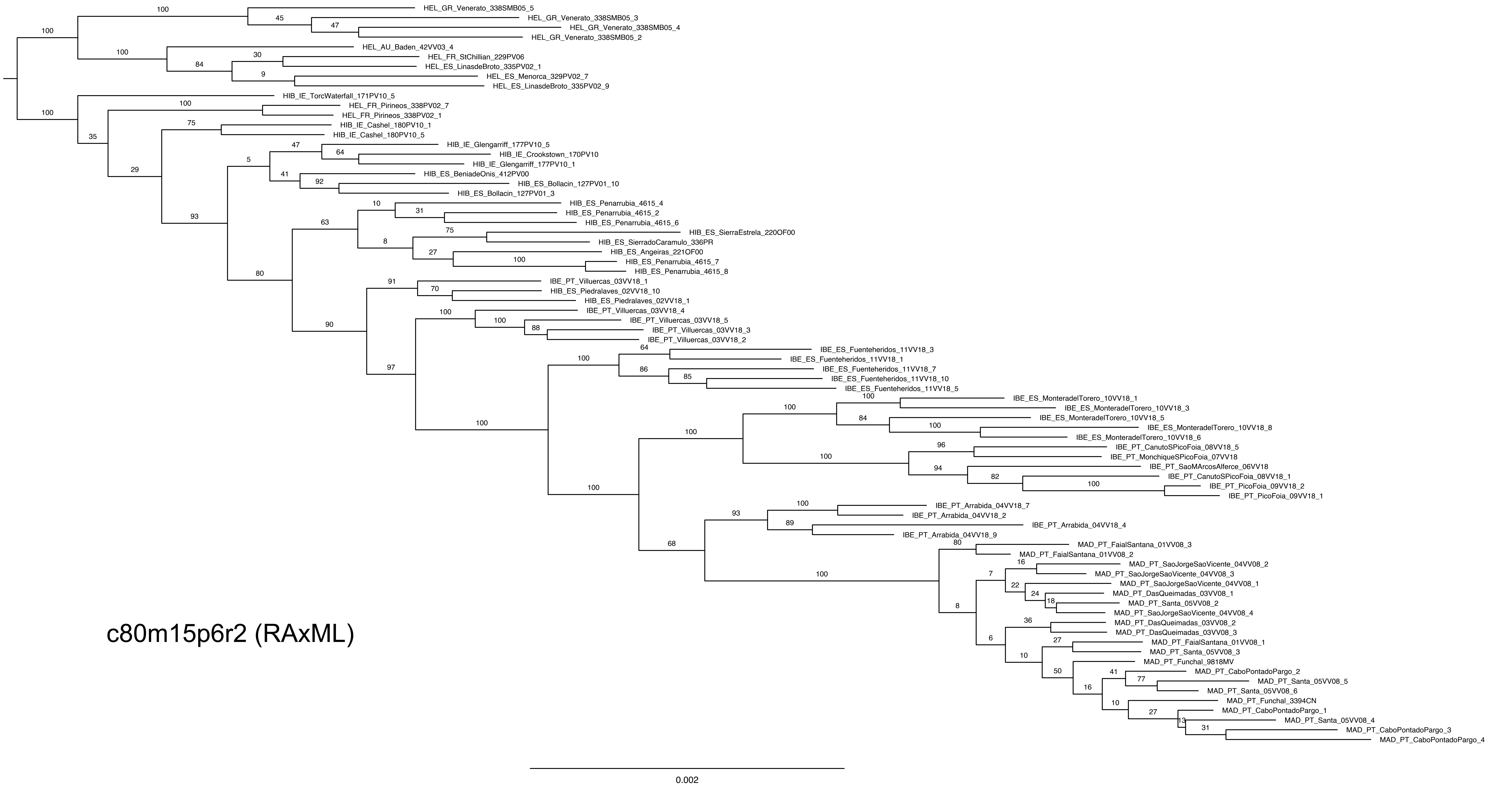

HEL_GR_Venerato_338SMB05_5
HEL_GR_Venerato_338SMB05_3
HEL_GR_Venerato_338SMB05_4
HEL_GR_Venerato_338SMB05_2
HEL_AU_Baden_42VV03_4
HEL_FR_StChillian_229PV06
HEL_ES_LinasdeBroto_335PV02_1
HEL_ES_Menorca_329PV02_7
HEL_ES_LinasdeBroto_335PV02_9
HIB_IE_TorcWaterfall_171PV10_5
HIB_FR_Pirineos_338PV02_7
HIB_FR_Pirineos_338PV02_1
HIB_IE_Cashel_180PV10_1
HIB_IE_Cashel_180PV10_5
HIB_IE_Glengarriff_177PV10_5
HIB_IE_Crookstown_170PV10
HIB_IE_Glengarriff_177PV10_1
HIB_ES_BeniadeOnis_412PV00
HIB_ES_Bollacin_127PV01_10
HIB_ES_Bollacin_127PV01_3
HIB_ES_Penarrubia_4615_4
HIB_ES_Penarrubia_4615_2
HIB_ES_Penarrubia_4615_6
HIB_ES_SierraEstrela_220OF00
HIB_ES_SierradoCaramulo_336PR
HIB_ES_Angeiras_221OF00
HIB_ES_Penarrubia_4615_7
HIB_ES_Penarrubia_4615_8
IBE_PT_Villuercas_03VV18_1
HIB_ES_Piedralaves_02VV18_10
HIB_ES_Piedralaves_02VV18_1
IBE_PT_Villuercas_03VV18_4
IBE_PT_Villuercas_03VV18_5
IBE_PT_Villuercas_03VV18_3
IBE_PT_Villuercas_03VV18_2
IBE_ES_Fuenteheridos_11VV18_3
IBE_ES_Fuenteheridos_11VV18_1
IBE_ES_Fuenteheridos_11VV18_7
IBE_ES_Fuenteheridos_11VV18_10
IBE_ES_Fuenteheridos_11VV18_5
IBE_ES_MonteradelTorero_10VV18_1
IBE_ES_MonteradelTorero_10VV18_3
IBE_ES_MonteradelTorero_10VV18_5
IBE_ES_MonteradelTorero_10VV18_8
IBE_ES_MonteradelTorero_10VV18_6
IBE_PT_CanutoSPicoFoia_08VV18_5
IBE_PT_MonchiqueSPicoFoia_07VV18
IBE_PT_SaoMArcosAlferce_06VV18
IBE_PT_CanutoSPicoFoia_08VV18_1
IBE_PT_PicoFoia_09VV18_2
IBE_PT_PicoFoia_09VV18_1
IBE_PT_Arrabida_04VV18_7
IBE_PT_Arrabida_04VV18_2
IBE_PT_Arrabida_04VV18_4
IBE_PT_Arrabida_04VV18_9
MAD_PT_FaialSantana_01VV08_3
MAD_PT_FaialSantana_01VV08_2
MAD_PT_SaoJorgeSaoVicente_04VV08_2
MAD_PT_SaoJorgeSaoVicente_04VV08_3
MAD_PT_SaoJorgeSaoVicente_04VV08_1
MAD_PT_DasQueimadas_03VV08_1
MAD_PT_Santa_05VV08_2
MAD_PT_SaoJorgeSaoVicente_04VV08_4
MAD_PT_DasQueimadas_03VV08_2
MAD_PT_DasQueimadas_03VV08_3
MAD_PT_FaialSantana_01VV08_1
MAD_PT_Santa_05VV08_3
MAD_PT_Funchal_9818MV
MAD_PT_CaboPontadoPargo_2
MAD_PT_Santa_05VV08_5
MAD_PT_Santa_05VV08_6
MAD_PT_Funchal_3394CN
MAD_PT_CaboPontadoPargo_1
MAD_PT_Santa_05VV08_4
MAD_PT_CaboPontadoPargo_3
MAD_PT_CaboPontadoPargo_4
c80m15p6r2 (RAxML)
0.002

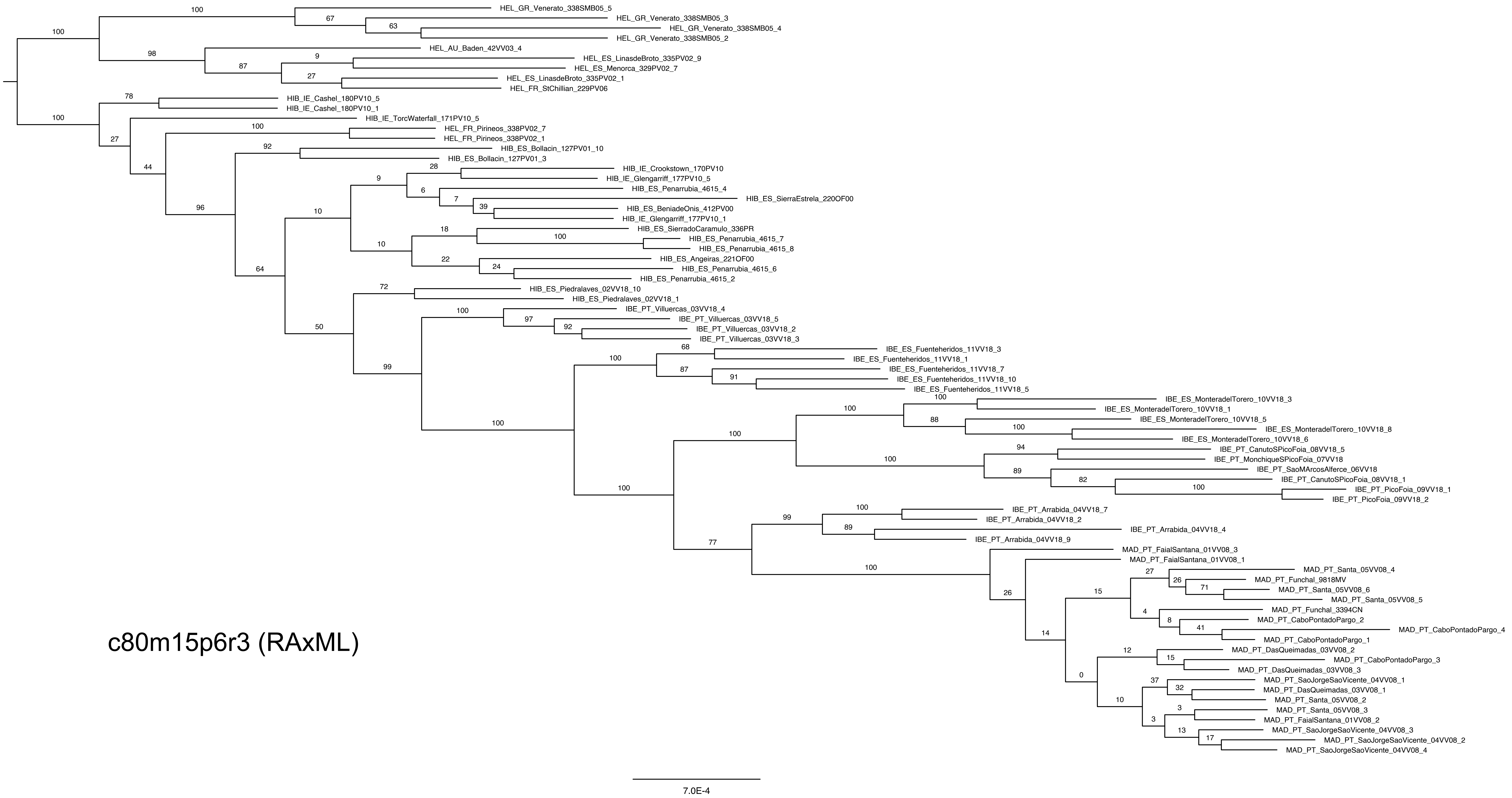
HEL_GR_Venerato_338SMB05_5
HEL_GR_Venerato_338SMB05_3
HEL_GR_Venerato_338SMB05_4
HEL_GR_Venerato_338SMB05_2
HEL_AU_Baden_42VV03_4
HEL_ES_LinasdeBroto_335PV02_9
HEL_ES_Menorca_329PV02_7
HEL_ES_LinasdeBroto_335PV02_1
HEL_FR_StChillian_229PV06
HIB_IE_Cashel_180PV10_5
HIB_IE_Cashel_180PV10_1
HIB_IE_TorcWaterfall_171PV10_5
HEL_FR_Pirineos_338PV02_7
HEL_FR_Pirineos_338PV02_1
HIB_ES_Bollacin_127PV01_10
HIB_ES_Bollacin_127PV01_3
HIB_IE_Crookstown_170PV10
HIB_IE_Glengarriff_177PV10_5
HIB_ES_Penarrubia_4615_4
HIB_ES_SierraEstrela_220OF00
HIB_ES_BeniadeOnis_412PV00
HIB_IE_Glengarriff_177PV10_1
HIB_ES_SierradoCaramulo_336PR
HIB_ES_Penarrubia_4615_7
HIB_ES_Penarrubia_4615_8
HIB_ES_Angeiras_221OF00
HIB_ES_Penarrubia_4615_6
HIB_ES_Penarrubia_4615_2
HIB_ES_Piedralaves_02VV18_10
HIB_ES_Piedralaves_02VV18_1
IBE_PT_Villuercas_03VV18_4
IBE_PT_Villuercas_03VV18_5
IBE_PT_Villuercas_03VV18_2
IBE_PT_Villuercas_03VV18_3
IBE_ES_Fuenteheridos_11VV18_3
IBE_ES_Fuenteheridos_11VV18_1
IBE_ES_Fuenteheridos_11VV18_7
IBE_ES_Fuenteheridos_11VV18_10
IBE_ES_Fuenteheridos_11VV18_5
IBE_ES_MonteradelTorero_10VV18_3
IBE_ES_MonteradelTorero_10VV18_1
IBE_ES_MonteradelTorero_10VV18_5
IBE_ES_MonteradelTorero_10VV18_8
IBE_ES_MonteradelTorero_10VV18_6
IBE_PT_CanutoSPicoFoia_08VV18_5
IBE_PT_MonchiqueSPicoFoia_07VV18
IBE_PT_SaoMArcosAlferce_06VV18
IBE_PT_CanutoSPicoFoia_08VV18_1
IBE_PT_PicoFoia_09VV18_1
IBE_PT_PicoFoia_09VV18_2
IBE_PT_Arrabida_04VV18_7
IBE_PT_Arrabida_04VV18_2
IBE_PT_Arrabida_04VV18_4
IBE_PT_Arrabida_04VV18_9
MAD_PT_FaialSantana_01VV08_3
MAD_PT_FaialSantana_01VV08_1
MAD_PT_Santa_05VV08_4
MAD_PT_Funchal_9818MV
MAD_PT_Santa_05VV08_6
MAD_PT_Santa_05VV08_5
MAD_PT_Funchal_3394CN
MAD_PT_CaboPontadoPargo_2
MAD_PT_CaboPontadoPargo_4
MAD_PT_CaboPontadoPargo_1
MAD_PT_DasQueimadas_03VV08_2
MAD_PT_CaboPontadoPargo_3
MAD_PT_DasQueimadas_03VV08_3
MAD_PT_SaoJorgeSaoVicente_04VV08_1
MAD_PT_DasQueimadas_03VV08_1
MAD_PT_Santa_05VV08_2
MAD_PT_Santa_05VV08_3
MAD_PT_FaialSantana_01VV08_2
MAD_PT_SaoJorgeSaoVicente_04VV08_3
MAD_PT_SaoJorgeSaoVicente_04VV08_2
MAD_PT_SaoJorgeSaoVicente_04VV08_4
c80m15p6r3 (RAxML)
7.0E-4

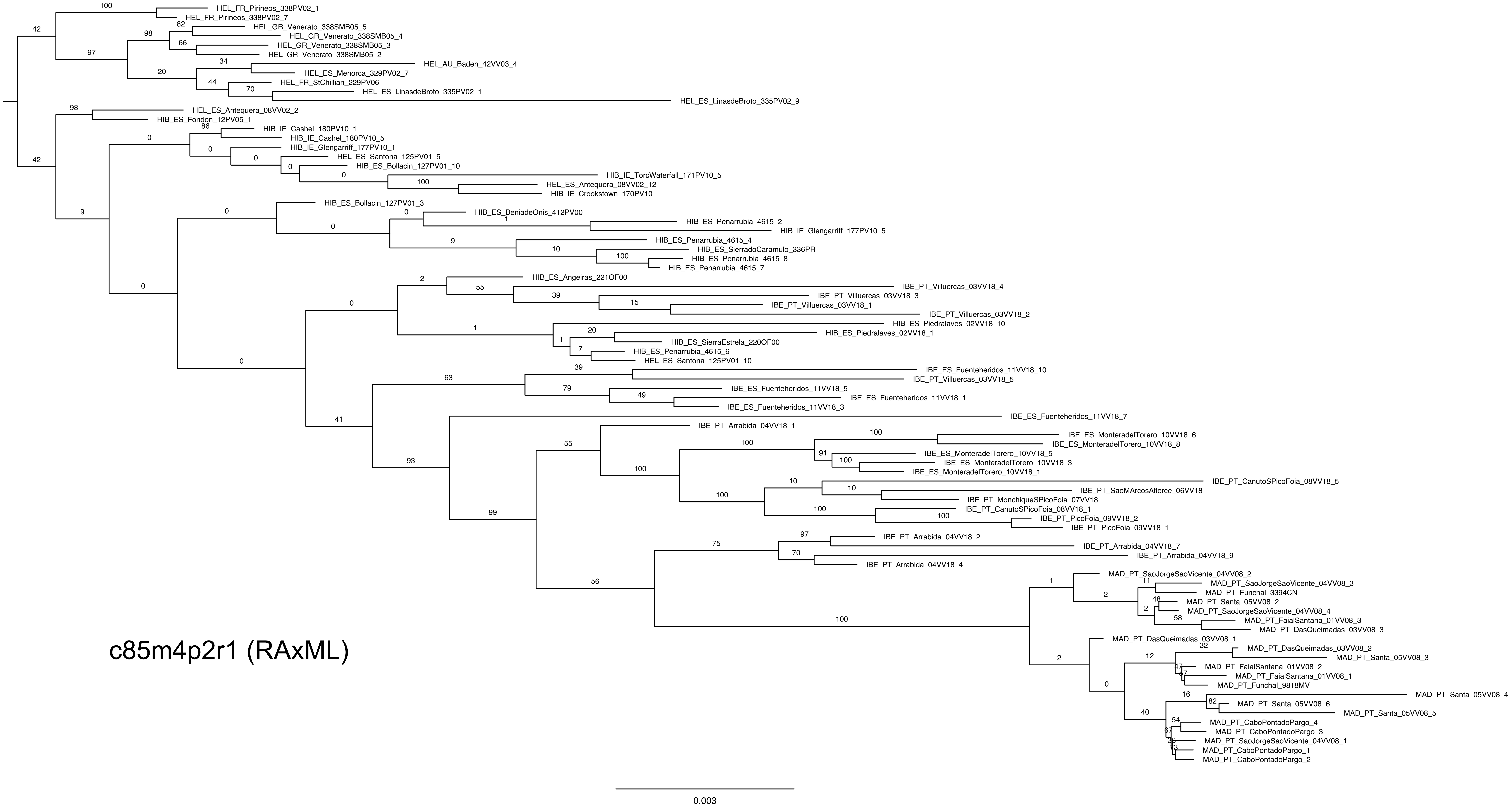

HEL_FR_Pirineos_338PV02_1
HEL_FR_Pirineos_338PV02_7
HEL_GR_Venerato_338SMB05_5
HEL_GR_Venerato_338SMB05_4
HEL_GR_Venerato_338SMB05_3
HEL_GR_Venerato_338SMB05_2
HEL_AU_Baden_42VV03_4
HEL_ES_Menorca_329PV02_7
HEL_FR_StChillian_229PV06
HEL_ES_LinasdeBroto_335PV02_1
HEL_ES_LinasdeBroto_335PV02_9
HEL_ES_Antequera_08VV02_2
HIB_ES_Fondon_12PV05_1
HIB_IE_Cashel_180PV10_1
HIB_IE_Cashel_180PV10_5
HIB_IE_Glengarriff_177PV10_1
HEL_ES_Santona_125PV01_5
HIB_ES_Bollacin_127PV01_10
HIB_IE_TorcWaterfall_171PV10_5
HEL_ES_Antequera_08VV02_12
HIB_IE_Crookstown_170PV10
HIB_ES_Bollacin_127PV01_3
HIB_ES_BeniadeOnis_412PV00
HIB_ES_Penarrubia_4615_2
HIB_IE_Glengarriff_177PV10_5
HIB_ES_Penarrubia_4615_4
HIB_ES_SierradoCaramulo_336PR
HIB_ES_Penarrubia_4615_8
HIB_ES_Penarrubia_4615_7
HIB_ES_Angeiras_221OF00
IBE_PT_Villuercas_03VV18_4
IBE_PT_Villuercas_03VV18_3
IBE_PT_Villuercas_03VV18_1
IBE_PT_Villuercas_03VV18_2
HIB_ES_Piedralaves_02VV18_10
HIB_ES_Piedralaves_02VV18_1
HIB_ES_SierraEstrela_220OF00
HIB_ES_Penarrubia_4615_6
HEL_ES_Santona_125PV01_10
IBE_ES_Fuenteheridos_11VV18_10
IBE_PT_Villuercas_03VV18_5
IBE_ES_Fuenteheridos_11VV18_5
IBE_ES_Fuenteheridos_11VV18_1
IBE_ES_Fuenteheridos_11VV18_3
IBE_ES_Fuenteheridos_11VV18_7
IBE_PT_Arrabida_04VV18_1
IBE_ES_MonteradelTorero_10VV18_6
IBE_ES_MonteradelTorero_10VV18_8
IBE_ES_MonteradelTorero_10VV18_5
IBE_ES_MonteradelTorero_10VV18_3
IBE_ES_MonteradelTorero_10VV18_1
IBE_PT_CanutoSPicoFoia_08VV18_5
IBE_PT_SaoMArcosAlferce_06VV18
IBE_PT_MonchiqueSPicoFoia_07VV18
IBE_PT_CanutoSPicoFoia_08VV18_1
IBE_PT_PicoFoia_09VV18_2
IBE_PT_PicoFoia_09VV18_1
IBE_PT_Arrabida_04VV18_2
IBE_PT_Arrabida_04VV18_7
IBE_PT_Arrabida_04VV18_9
IBE_PT_Arrabida_04VV18_4
MAD_PT_SaoJorgeSaoVicente_04VV08_2
MAD_PT_SaoJorgeSaoVicente_04VV08_3
MAD_PT_Funchal_3394CN
MAD_PT_Santa_05VV08_2
MAD_PT_SaoJorgeSaoVicente_04VV08_4
MAD_PT_FaialSantana_01VV08_3
MAD_PT_DasQueimadas_03VV08_3
MAD_PT_DasQueimadas_03VV08_1
MAD_PT_DasQueimadas_03VV08_2
MAD_PT_Santa_05VV08_3
MAD_PT_FaialSantana_01VV08_2
MAD_PT_FaialSantana_01VV08_1
MAD_PT_Funchal_9818MV
MAD_PT_Santa_05VV08_4
MAD_PT_Santa_05VV08_6
MAD_PT_Santa_05VV08_5
MAD_PT_CaboPontadoPargo_4
MAD_PT_CaboPontadoPargo_3
MAD_PT_SaoJorgeSaoVicente_04VV08_1
MAD_PT_CaboPontadoPargo_1
MAD_PT_CaboPontadoPargo_2
c85m4p2r1 (RAxML)
0.003

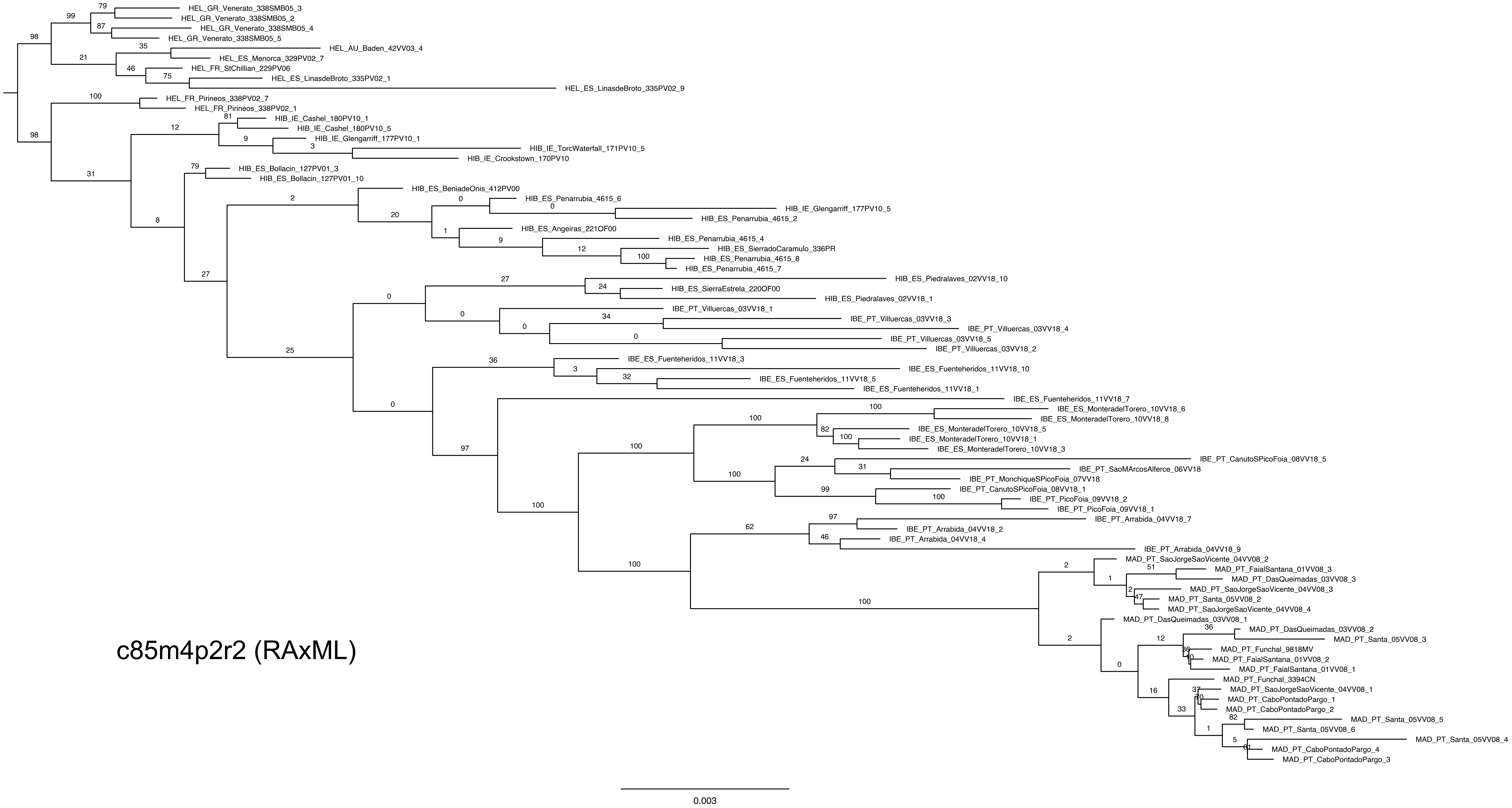
HEL_GR_Venerato_338SMB05_3
HEL_GR_Venerato_338SMB05_2
HEL_GR_Venerato_338SMB05_4
HEL_GR_Venerato_338SMB05_5
HEL_AU_Baden_42VV03_4
HEL_ES_Menorca_329PV02_7
HEL_FR_StChillian_229PV06
HEL_ES_LinasdeBroto_335PV02_1
HEL_ES_LinasdeBroto_335PV02_9
HEL_FR_Pirineos_338PV02_7
HEL_FR_Pirineos_338PV02_1
HIB_IE_Cashel_180PV10_1
HIB_IE_Cashel_180PV10_5
HIB_IE_Glengarriff_177PV10_1
HIB_IE_TorcWaterfall_171PV10_5
HIB_IE_Crookstown_170PV10
HIB_ES_Bollacin_127PV01_3
HIB_ES_Bollacin_127PV01_10
HIB_ES_BeniadeOnis_412PV00
HIB_ES_Penarrubia_4615_6
HIB_IE_Glengarriff_177PV10_5
HIB_ES_Penarrubia_4615_2
HIB_ES_Angeiras_221OF00
HIB_ES_Penarrubia_4615_4
HIB_ES_SierradoCaramulo_336PR
HIB_ES_Penarrubia_4615_8
HIB_ES_Penarrubia_4615_7
HIB_ES_Piedralaves_02VV18_10
HIB_ES_SierraEstrela_220OF00
HIB_ES_Piedralaves_02VV18_1
IBE_PT_Villuercas_03VV18_1
IBE_PT_Villuercas_03VV18_3
IBE_PT_Villuercas_03VV18_4
IBE_PT_Villuercas_03VV18_5
IBE_PT_Villuercas_03VV18_2
IBE_ES_Fuenteheridos_11VV18_3
IBE_ES_Fuenteheridos_11VV18_10
IBE_ES_Fuenteheridos_11VV18_5
IBE_ES_Fuenteheridos_11VV18_1
IBE_ES_Fuenteheridos_11VV18_7
IBE_ES_MonteradelTorero_10VV18_6
IBE_ES_MonteradelTorero_10VV18_8
IBE_ES_MonteradelTorero_10VV18_5
IBE_ES_MonteradelTorero_10VV18_1
IBE_ES_MonteradelTorero_10VV18_3
IBE_PT_CanutoSPicoFoia_08VV18_5
IBE_PT_SaoMArcosAlferce_06VV18
IBE_PT_MonchiqueSPicoFoia_07VV18
IBE_PT_CanutoSPicoFoia_08VV18_1
IBE_PT_PicoFoia_09VV18_2
IBE_PT_PicoFoia_09VV18_1
IBE_PT_Arrabida_04VV18_7
IBE_PT_Arrabida_04VV18_2
IBE_PT_Arrabida_04VV18_4
IBE_PT_Arrabida_04VV18_9
MAD_PT_SaoJorgeSaoVicente_04VV08_2
MAD_PT_FaialSantana_01VV08_3
MAD_PT_DasQueimadas_03VV08_3
MAD_PT_SaoJorgeSaoVicente_04VV08_3
MAD_PT_Santa_05VV08_2
MAD_PT_SaoJorgeSaoVicente_04VV08_4
MAD_PT_DasQueimadas_03VV08_1
MAD_PT_DasQueimadas_03VV08_2
MAD_PT_Santa_05VV08_3
MAD_PT_Funchal_9818MV
MAD_PT_FaialSantana_01VV08_2
MAD_PT_FaialSantana_01VV08_1
MAD_PT_Funchal_3394CN
MAD_PT_SaoJorgeSaoVicente_04VV08_1
MAD_PT_CaboPontadoPargo_1
MAD_PT_CaboPontadoPargo_2
MAD_PT_Santa_05VV08_5
MAD_PT_Santa_05VV08_6
MAD_PT_Santa_05VV08_4
MAD_PT_CaboPontadoPargo_4
MAD_PT_CaboPontadoPargo_3
c85m4p2r2 (RAxML)
0.003

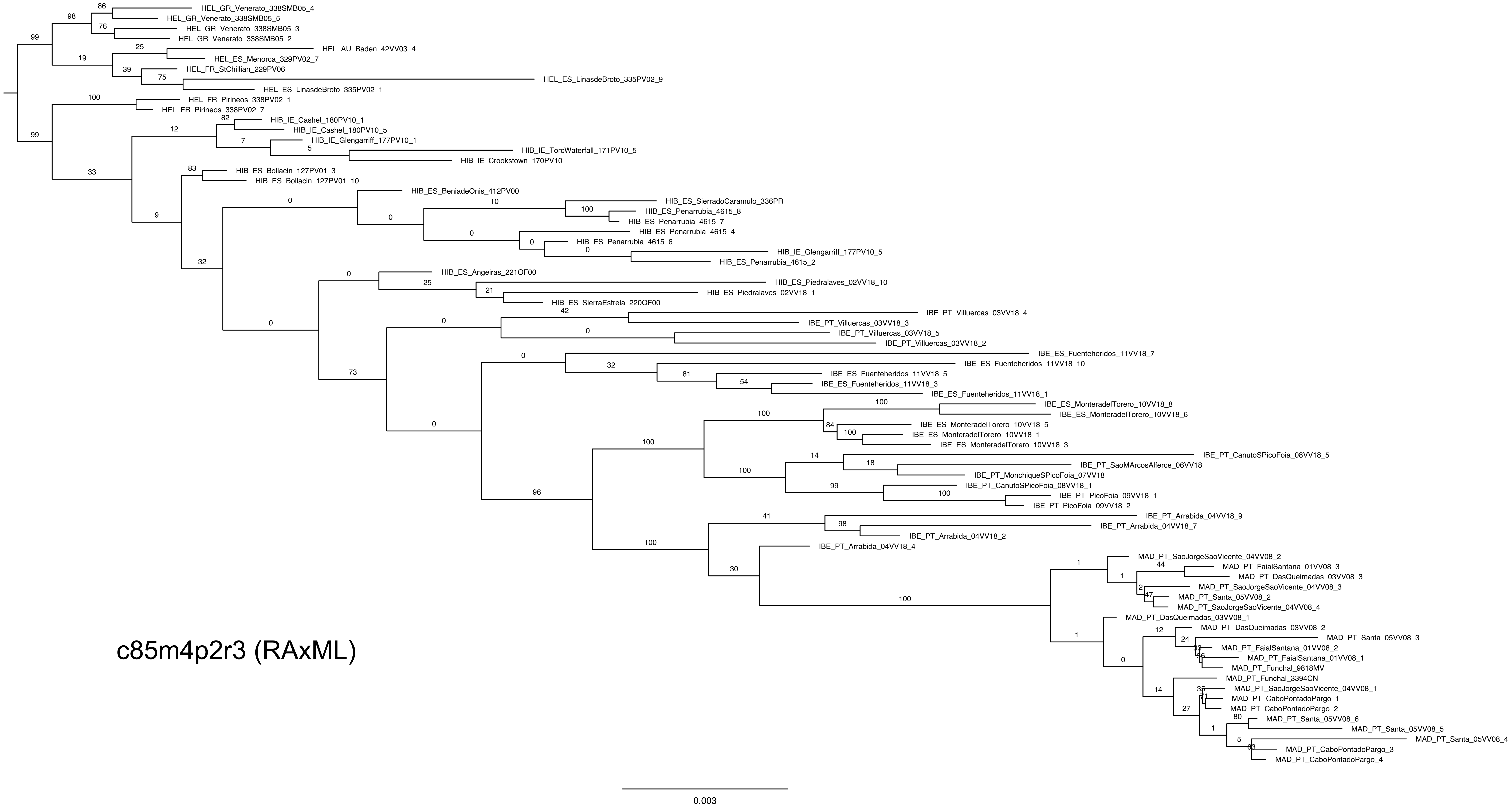
HEL_GR_Venerato_338SMB05_4
HEL_GR_Venerato_338SMB05_5
HEL_GR_Venerato_338SMB05_3
HEL_GR_Venerato_338SMB05_2
HEL_AU_Baden_42VV03_4
HEL_ES_Menorca_329PV02_7
HEL_FR_StChillian_229PV06
HEL_ES_LinasdeBroto_335PV02_9
HEL_ES_LinasdeBroto_335PV02_1
HEL_FR_Pirineos_338PV02_1
HEL_FR_Pirineos_338PV02_7
HIB_IE_Cashel_180PV10_1
HIB_IE_Cashel_180PV10_5
HIB_IE_Glengarriff_177PV10_1
HIB_IE_TorcWaterfall_171PV10_5
HIB_IE_Crookstown_170PV10
HIB_ES_Bollacin_127PV01_3
HIB_ES_Bollacin_127PV01_10
HIB_ES_BeniadeOnis_412PV00
HIB_ES_SierradoCaramulo_336PR
HIB_ES_Penarrubia_4615_8
HIB_ES_Penarrubia_4615_7
HIB_ES_Penarrubia_4615_4
HIB_ES_Penarrubia_4615_6
HIB_IE_Glengarriff_177PV10_5
HIB_ES_Penarrubia_4615_2
HIB_ES_Angeiras_221OF00
HIB_ES_Piedralaves_02VV18_10
HIB_ES_Piedralaves_02VV18_1
HIB_ES_SierraEstrela_220OF00
IBE_PT_Villuercas_03VV18_4
IBE_PT_Villuercas_03VV18_3
IBE_PT_Villuercas_03VV18_5
IBE_PT_Villuercas_03VV18_2
IBE_ES_Fuenteheridos_11VV18_7
IBE_ES_Fuenteheridos_11VV18_10
IBE_ES_Fuenteheridos_11VV18_5
IBE_ES_Fuenteheridos_11VV18_3
IBE_ES_Fuenteheridos_11VV18_1
IBE_ES_MonteradelTorero_10VV18_8
IBE_ES_MonteradelTorero_10VV18_6
IBE_ES_MonteradelTorero_10VV18_5
IBE_ES_MonteradelTorero_10VV18_1
IBE_ES_MonteradelTorero_10VV18_3
IBE_PT_CanutoSPicoFoia_08VV18_5
IBE_PT_SaoMArcosAlferce_06VV18
IBE_PT_MonchiqueSPicoFoia_07VV18
IBE_PT_CanutoSPicoFoia_08VV18_1
IBE_PT_PicoFoia_09VV18_1
IBE_PT_PicoFoia_09VV18_2
IBE_PT_Arrabida_04VV18_9
IBE_PT_Arrabida_04VV18_7
IBE_PT_Arrabida_04VV18_2
IBE_PT_Arrabida_04VV18_4
MAD_PT_SaoJorgeSaoVicente_04VV08_2
MAD_PT_FaialSantana_01VV08_3
MAD_PT_DasQueimadas_03VV08_3
MAD_PT_SaoJorgeSaoVicente_04VV08_3
MAD_PT_Santa_05VV08_2
MAD_PT_SaoJorgeSaoVicente_04VV08_4
MAD_PT_DasQueimadas_03VV08_1
MAD_PT_DasQueimadas_03VV08_2
MAD_PT_Santa_05VV08_3
MAD_PT_FaialSantana_01VV08_2
MAD_PT_FaialSantana_01VV08_1
MAD_PT_Funchal_9818MV
MAD_PT_Funchal_3394CN
MAD_PT_SaoJorgeSaoVicente_04VV08_1
MAD_PT_CaboPontadoPargo_1
MAD_PT_CaboPontadoPargo_2
MAD_PT_Santa_05VV08_6
MAD_PT_Santa_05VV08_5
MAD_PT_Santa_05VV08_4
MAD_PT_CaboPontadoPargo_3
MAD_PT_CaboPontadoPargo_4
c85m4p2r3 (RAxML)
0.003

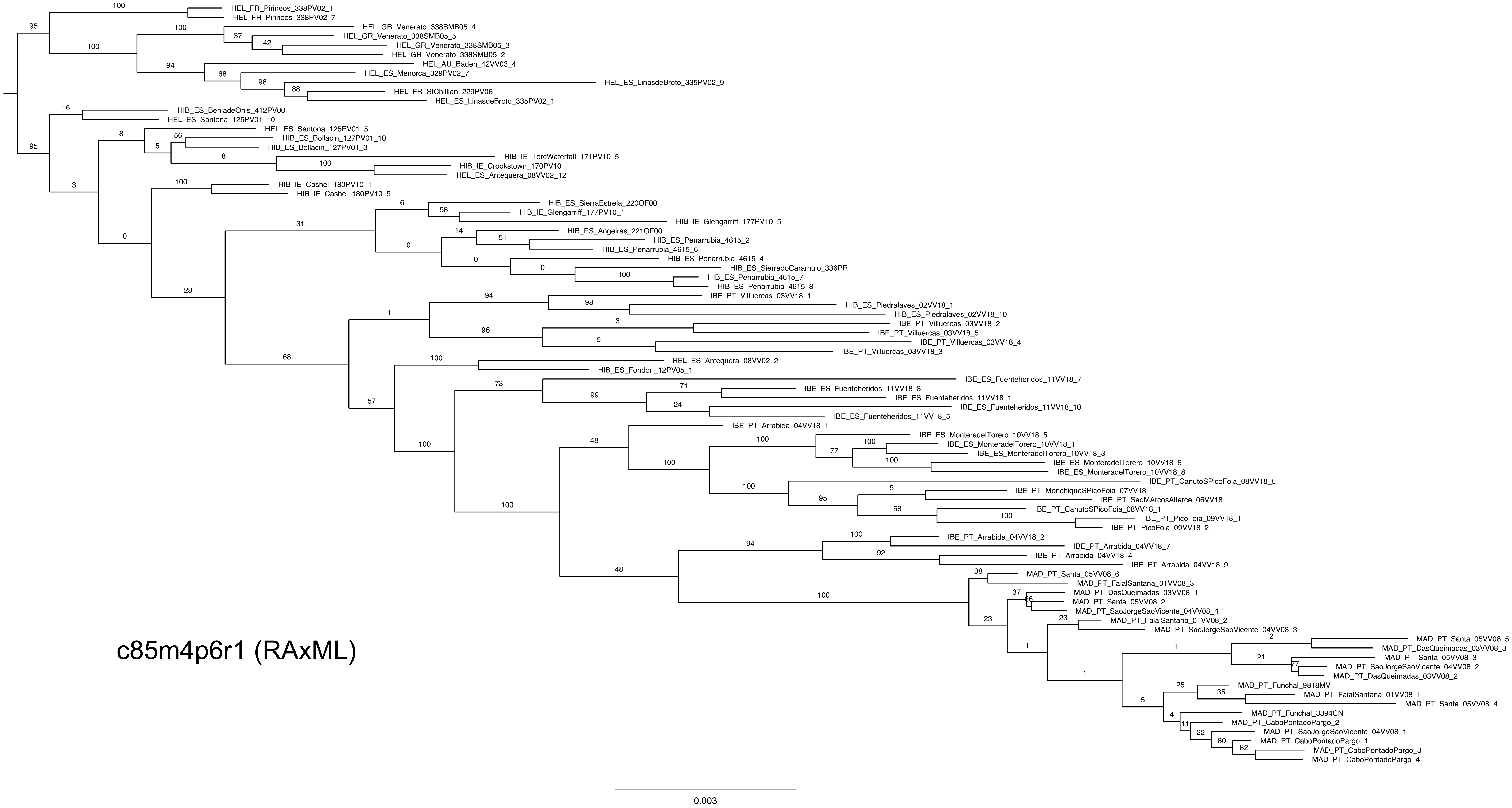
HEL_FR_Pirineos_338PV02_1
HEL_FR_Pirineos_338PV02_7
HEL_GR_Venerato_338SMB05_4
HEL_GR_Venerato_338SMB05_5
HEL_GR_Venerato_338SMB05_3
HEL_GR_Venerato_338SMB05_2
HEL_AU_Baden_42VV03_4
HEL_ES_Menorca_329PV02_7
HEL_ES_LinasdeBroto_335PV02_9
HEL_FR_StChillian_229PV06
HEL_ES_LinasdeBroto_335PV02_1
HIB_ES_BeniadeOnis_412PV00
HEL_ES_Santona_125PV01_10
HEL_ES_Santona_125PV01_5
HIB_ES_Bollacin_127PV01_10
HIB_ES_Bollacin_127PV01_3
HIB_IE_TorcWaterfall_171PV10_5
HIB_IE_Crookstown_170PV10
HEL_ES_Antequera_08VV02_12
HIB_IE_Cashel_180PV10_1
HIB_IE_Cashel_180PV10_5
HIB_ES_SierraEstrela_220OF00
HIB_IE_Glengarriff_177PV10_1
HIB_IE_Glengarriff_177PV10_5
HIB_ES_Angeiras_221OF00
HIB_ES_Penarrubia_4615_2
HIB_ES_Penarrubia_4615_6
HIB_ES_Penarrubia_4615_4
HIB_ES_SierradoCaramulo_336PR
HIB_ES_Penarrubia_4615_7
HIB_ES_Penarrubia_4615_8
IBE_PT_Villuercas_03VV18_1
HIB_ES_Piedralaves_02VV18_1
HIB_ES_Piedralaves_02VV18_10
IBE_PT_Villuercas_03VV18_2
IBE_PT_Villuercas_03VV18_5
IBE_PT_Villuercas_03VV18_4
IBE_PT_Villuercas_03VV18_3
HEL_ES_Antequera_08VV02_2
HIB_ES_Fondon_12PV05_1
IBE_ES_Fuenteheridos_11VV18_7
IBE_ES_Fuenteheridos_11VV18_3
IBE_ES_Fuenteheridos_11VV18_1
IBE_ES_Fuenteheridos_11VV18_10
IBE_ES_Fuenteheridos_11VV18_5
IBE_PT_Arrabida_04VV18_1
IBE_ES_MonteradelTorero_10VV18_5
IBE_ES_MonteradelTorero_10VV18_1
IBE_ES_MonteradelTorero_10VV18_3
IBE_ES_MonteradelTorero_10VV18_6
IBE_ES_MonteradelTorero_10VV18_8
IBE_PT_CanutoSPicoFoia_08VV18_5
IBE_PT_MonchiqueSPicoFoia_07VV18
IBE_PT_SaoMArcosAlferce_06VV18
IBE_PT_CanutoSPicoFoia_08VV18_1
IBE_PT_PicoFoia_09VV18_1
IBE_PT_PicoFoia_09VV18_2
IBE_PT_Arrabida_04VV18_2
IBE_PT_Arrabida_04VV18_7
IBE_PT_Arrabida_04VV18_4
IBE_PT_Arrabida_04VV18_9
MAD_PT_Santa_05VV08_6
MAD_PT_FaialSantana_01VV08_3
MAD_PT_DasQueimadas_03VV08_1
MAD_PT_Santa_05VV08_2
MAD_PT_SaoJorgeSaoVicente_04VV08_4
MAD_PT_FaialSantana_01VV08_2
MAD_PT_SaoJorgeSaoVicente_04VV08_3
MAD_PT_Santa_05VV08_5
MAD_PT_DasQueimadas_03VV08_3
MAD_PT_Santa_05VV08_3
MAD_PT_SaoJorgeSaoVicente_04VV08_2
MAD_PT_DasQueimadas_03VV08_2
MAD_PT_Funchal_9818MV
MAD_PT_FaialSantana_01VV08_1
MAD_PT_Santa_05VV08_4
MAD_PT_Funchal_3394CN
MAD_PT_CaboPontadoPargo_2
MAD_PT_SaoJorgeSaoVicente_04VV08_1
MAD_PT_CaboPontadoPargo_1
MAD_PT_CaboPontadoPargo_3
MAD_PT_CaboPontadoPargo_4
c85m4p6r1 (RAxML)
0.003

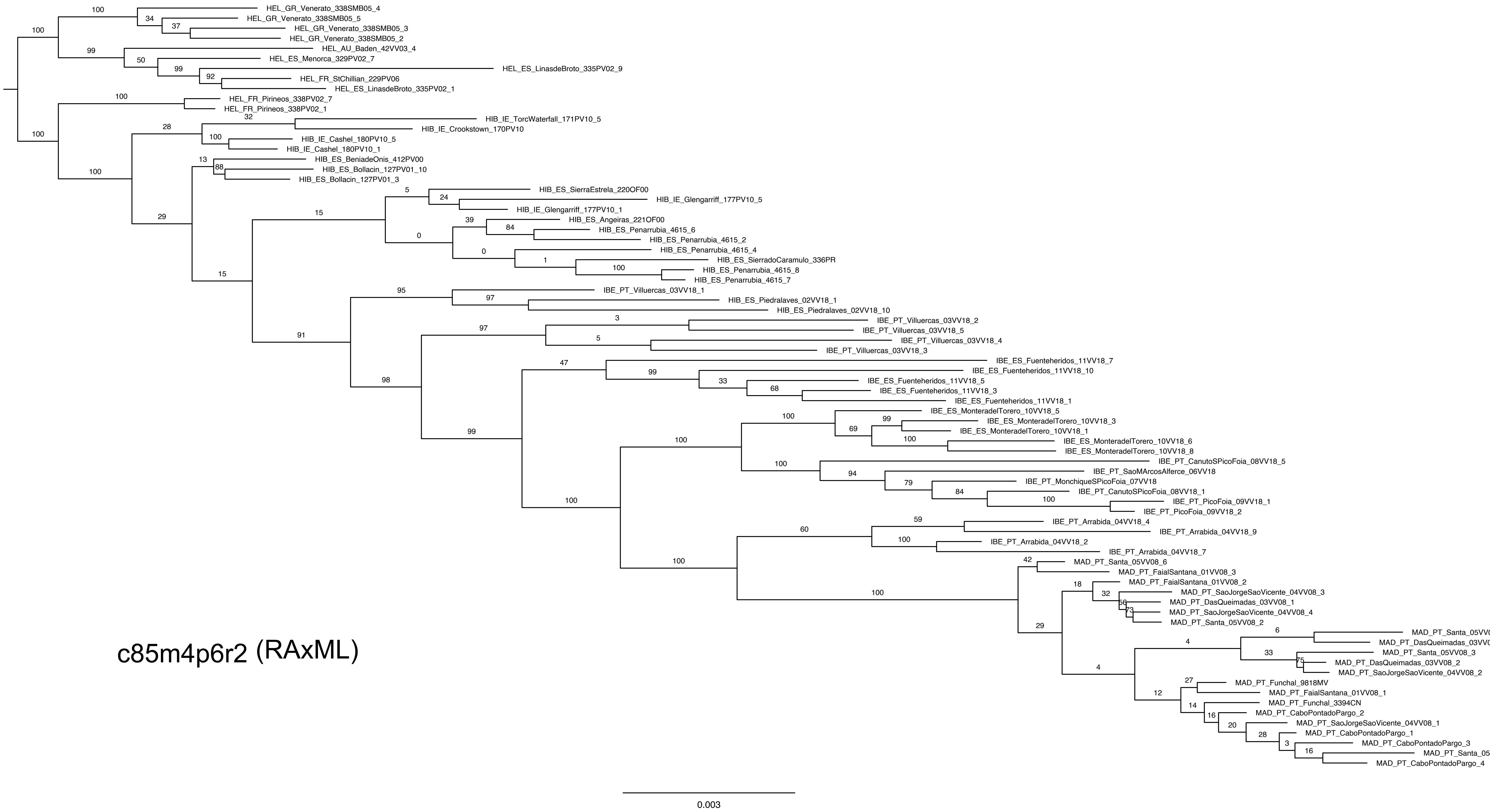
HEL_GR_Venerato_338SMB05_4
HEL_GR_Venerato_338SMB05_5
HEL_GR_Venerato_338SMB05_3
HEL_GR_Venerato_338SMB05_2
HEL_AU_Baden_42VV03_4
HEL_ES_Menorca_329PV02_7
HEL_ES_LinasdeBroto_335PV02_9
HEL_FR_StChillian_229PV06
HEL_ES_LinasdeBroto_335PV02_1
HEL_FR_Pirineos_338PV02_7
HEL_FR_Pirineos_338PV02_1
HIB_IE_TorcWaterfall_171PV10_5
HIB_IE_Crookstown_170PV10
HIB_IE_Cashel_180PV10_5
HIB_IE_Cashel_180PV10_1
HIB_ES_BeniadeOnis_412PV00
HIB_ES_Bollacin_127PV01_10
HIB_ES_Bollacin_127PV01_3
HIB_ES_SierraEstrela_220OF00
HIB_IE_Glengarriff_177PV10_5
HIB_IE_Glengarriff_177PV10_1
HIB_ES_Angeiras_221OF00
HIB_ES_Penarrubia_4615_6
HIB_ES_Penarrubia_4615_2
HIB_ES_Penarrubia_4615_4
HIB_ES_SierradoCaramulo_336PR
HIB_ES_Penarrubia_4615_8
HIB_ES_Penarrubia_4615_7
IBE_PT_Villuercas_03VV18_1
HIB_ES_Piedralaves_02VV18_1
HIB_ES_Piedralaves_02VV18_10
IBE_PT_Villuercas_03VV18_2
IBE_PT_Villuercas_03VV18_5
IBE_PT_Villuercas_03VV18_4
IBE_PT_Villuercas_03VV18_3
IBE_ES_Fuenteheridos_11VV18_7
IBE_ES_Fuenteheridos_11VV18_10
IBE_ES_Fuenteheridos_11VV18_5
IBE_ES_Fuenteheridos_11VV18_3
IBE_ES_Fuenteheridos_11VV18_1
IBE_ES_MonteradelTorero_10VV18_5
IBE_ES_MonteradelTorero_10VV18_3
IBE_ES_MonteradelTorero_10VV18_1
IBE_ES_MonteradelTorero_10VV18_6
IBE_ES_MonteradelTorero_10VV18_8
IBE_PT_CanutoSPicoFoia_08VV18_5
IBE_PT_SaoMArcosAlferce_06VV18
IBE_PT_MonchiqueSPicoFoia_07VV18
IBE_PT_CanutoSPicoFoia_08VV18_1
IBE_PT_PicoFoia_09VV18_1
IBE_PT_PicoFoia_09VV18_2
IBE_PT_Arrabida_04VV18_4
IBE_PT_Arrabida_04VV18_9
IBE_PT_Arrabida_04VV18_2
IBE_PT_Arrabida_04VV18_7
MAD_PT_Santa_05VV08_6
MAD_PT_FaialSantana_01VV08_3
MAD_PT_FaialSantana_01VV08_2
MAD_PT_SaoJorgeSaoVicente_04VV08_3
MAD_PT_DasQueimadas_03VV08_1
MAD_PT_SaoJorgeSaoVicente_04VV08_4
MAD_PT_Santa_05VV08_2
MAD_PT_Santa_05VV08_5
MAD_PT_DasQueimadas_03VV08_3
MAD_PT_Santa_05VV08_3
MAD_PT_DasQueimadas_03VV08_2
MAD_PT_SaoJorgeSaoVicente_04VV08_2
MAD_PT_Funchal_9818MV
MAD_PT_FaialSantana_01VV08_1
MAD_PT_Funchal_3394CN
MAD_PT_CaboPontadoPargo_2
MAD_PT_SaoJorgeSaoVicente_04VV08_1
MAD_PT_CaboPontadoPargo_1
MAD_PT_CaboPontadoPargo_3
MAD_PT_Santa_05VV08_4
MAD_PT_CaboPontadoPargo_4
c85m4p6r2 (RAxML)
0.003

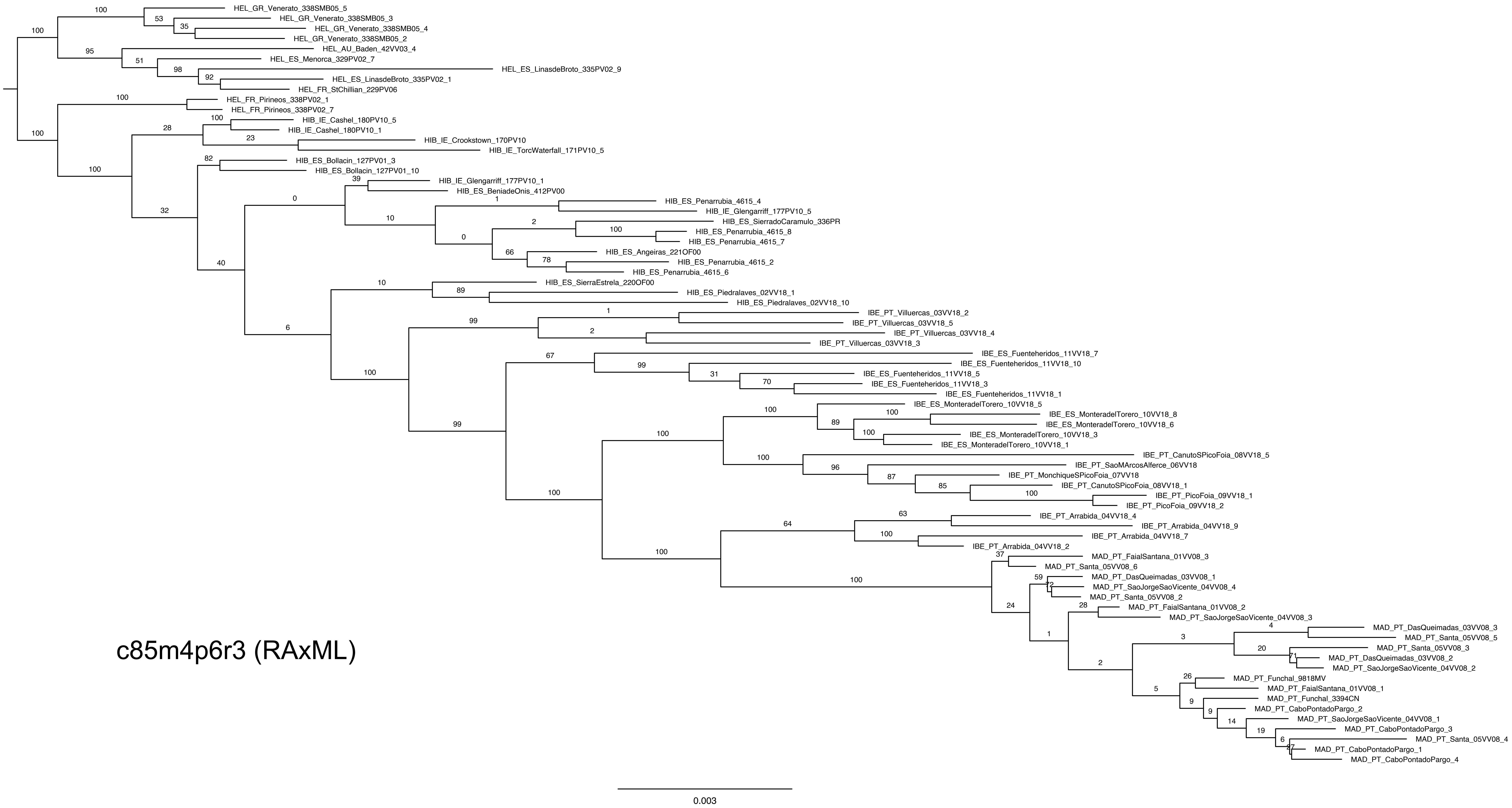
HEL_GR_Venerato_338SMB05_5
HEL_GR_Venerato_338SMB05_3
HEL_GR_Venerato_338SMB05_4
HEL_GR_Venerato_338SMB05_2
HEL_AU_Baden_42VV03_4
HEL_ES_Menorca_329PV02_7
HEL_ES_LinasdeBroto_335PV02_9
HEL_ES_LinasdeBroto_335PV02_1
HEL_FR_StChillian_229PV06
HEL_FR_Pirineos_338PV02_1
HEL_FR_Pirineos_338PV02_7
HIB_IE_Cashel_180PV10_5
HIB_IE_Cashel_180PV10_1
HIB_IE_Crookstown_170PV10
HIB_IE_TorcWaterfall_171PV10_5
HIB_ES_Bollacin_127PV01_3
HIB_ES_Bollacin_127PV01_10
HIB_IE_Glengarriff_177PV10_1
HIB_ES_BeniadeOnis_412PV00
HIB_ES_Penarrubia_4615_4
HIB_IE_Glengarriff_177PV10_5
HIB_ES_SierradoCaramulo_336PR
HIB_ES_Penarrubia_4615_8
HIB_ES_Penarrubia_4615_7
HIB_ES_Angeiras_221OF00
HIB_ES_Penarrubia_4615_2
HIB_ES_Penarrubia_4615_6
HIB_ES_SierraEstrela_220OF00
HIB_ES_Piedralaves_02VV18_1
HIB_ES_Piedralaves_02VV18_10
IBE_PT_Villuercas_03VV18_2
IBE_PT_Villuercas_03VV18_5
IBE_PT_Villuercas_03VV18_4
IBE_PT_Villuercas_03VV18_3
IBE_ES_Fuenteheridos_11VV18_7
IBE_ES_Fuenteheridos_11VV18_10
IBE_ES_Fuenteheridos_11VV18_5
IBE_ES_Fuenteheridos_11VV18_3
IBE_ES_Fuenteheridos_11VV18_1
IBE_ES_MonteradelTorero_10VV18_5
IBE_ES_MonteradelTorero_10VV18_8
IBE_ES_MonteradelTorero_10VV18_6
IBE_ES_MonteradelTorero_10VV18_3
IBE_ES_MonteradelTorero_10VV18_1
IBE_PT_CanutoSPicoFoia_08VV18_5
IBE_PT_SaoMArcosAlferce_06VV18
IBE_PT_MonchiqueSPicoFoia_07VV18
IBE_PT_CanutoSPicoFoia_08VV18_1
IBE_PT_PicoFoia_09VV18_1
IBE_PT_PicoFoia_09VV18_2
IBE_PT_Arrabida_04VV18_4
IBE_PT_Arrabida_04VV18_9
IBE_PT_Arrabida_04VV18_7
IBE_PT_Arrabida_04VV18_2
MAD_PT_FaialSantana_01VV08_3
MAD_PT_Santa_05VV08_6
MAD_PT_DasQueimadas_03VV08_1
MAD_PT_SaoJorgeSaoVicente_04VV08_4
MAD_PT_Santa_05VV08_2
MAD_PT_FaialSantana_01VV08_2
MAD_PT_SaoJorgeSaoVicente_04VV08_3
MAD_PT_DasQueimadas_03VV08_3
MAD_PT_Santa_05VV08_5
MAD_PT_Santa_05VV08_3
MAD_PT_DasQueimadas_03VV08_2
MAD_PT_SaoJorgeSaoVicente_04VV08_2
MAD_PT_Funchal_9818MV
MAD_PT_FaialSantana_01VV08_1
MAD_PT_Funchal_3394CN
MAD_PT_CaboPontadoPargo_2
MAD_PT_SaoJorgeSaoVicente_04VV08_1
MAD_PT_CaboPontadoPargo_3
MAD_PT_Santa_05VV08_4
MAD_PT_CaboPontadoPargo_1
MAD_PT_CaboPontadoPargo_4
c85m4p6r3 (RAxML)
0.003

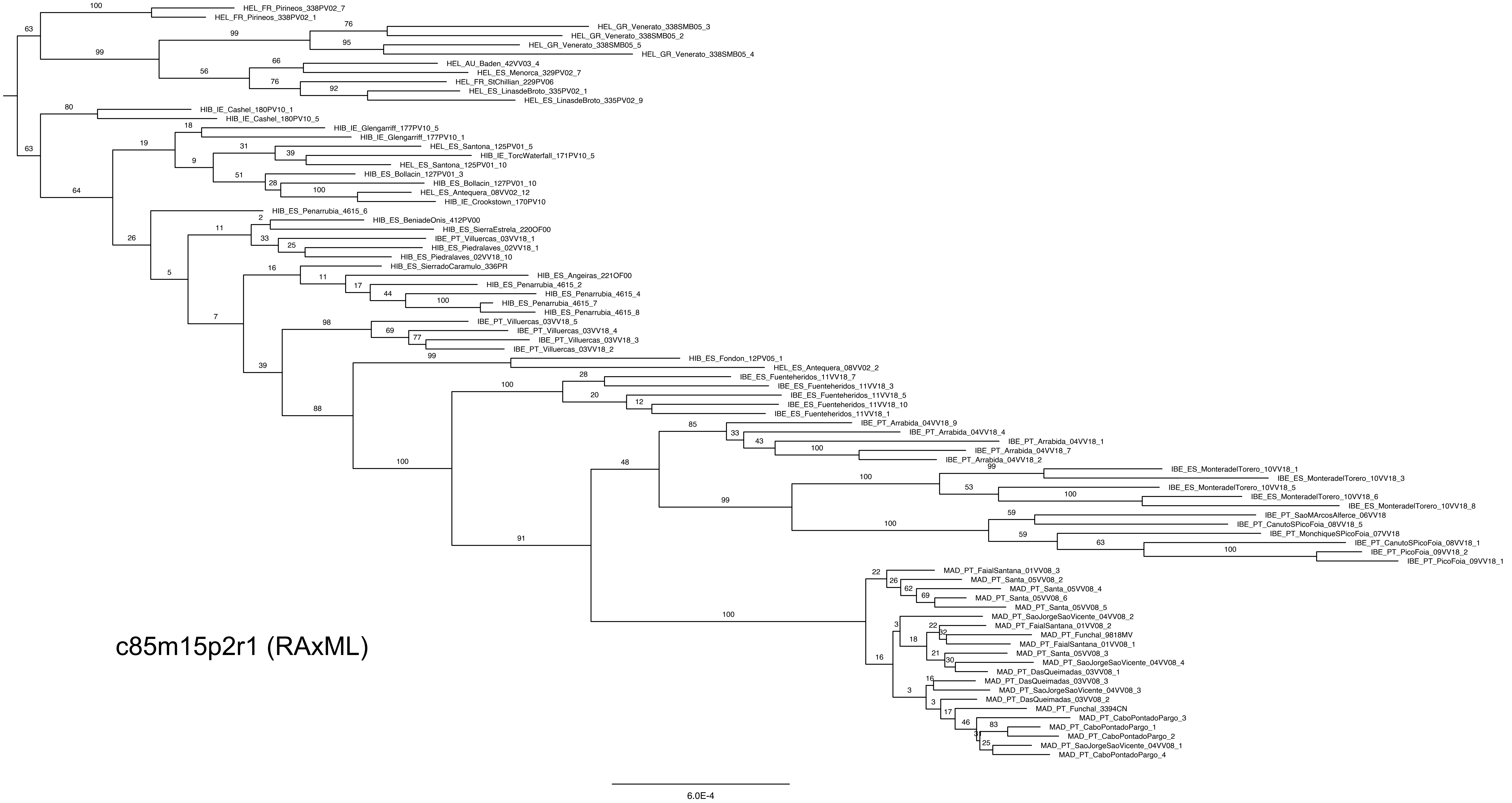
HEL_FR_Pirineos_338PV02_7
HEL_FR_Pirineos_338PV02_1
HEL_GR_Venerato_338SMB05_3
HEL_GR_Venerato_338SMB05_2
HEL_GR_Venerato_338SMB05_5
HEL_GR_Venerato_338SMB05_4
HEL_AU_Baden_42VV03_4
HEL_ES_Menorca_329PV02_7
HEL_FR_StChillian_229PV06
HEL_ES_LinasdeBroto_335PV02_1
HEL_ES_LinasdeBroto_335PV02_9
HIB_IE_Cashel_180PV10_1
HIB_IE_Cashel_180PV10_5
HIB_IE_Glengarriff_177PV10_5
HIB_IE_Glengarriff_177PV10_1
HEL_ES_Santona_125PV01_5
HIB_IE_TorcWaterfall_171PV10_5
HEL_ES_Santona_125PV01_10
HIB_ES_Bollacin_127PV01_3
HIB_ES_Bollacin_127PV01_10
HEL_ES_Antequera_08VV02_12
HIB_IE_Crookstown_170PV10
HIB_ES_Penarrubia_4615_6
HIB_ES_BeniadeOnis_412PV00
HIB_ES_SierraEstrela_220OF00
IBE_PT_Villuercas_03VV18_1
HIB_ES_Piedralaves_02VV18_1
HIB_ES_Piedralaves_02VV18_10
HIB_ES_SierradoCaramulo_336PR
HIB_ES_Angeiras_221OF00
HIB_ES_Penarrubia_4615_2
HIB_ES_Penarrubia_4615_4
HIB_ES_Penarrubia_4615_7
HIB_ES_Penarrubia_4615_8
IBE_PT_Villuercas_03VV18_5
IBE_PT_Villuercas_03VV18_4
IBE_PT_Villuercas_03VV18_3
IBE_PT_Villuercas_03VV18_2
HIB_ES_Fondon_12PV05_1
HEL_ES_Antequera_08VV02_2
IBE_ES_Fuenteheridos_11VV18_7
IBE_ES_Fuenteheridos_11VV18_3
IBE_ES_Fuenteheridos_11VV18_5
IBE_ES_Fuenteheridos_11VV18_10
IBE_ES_Fuenteheridos_11VV18_1
IBE_PT_Arrabida_04VV18_9
IBE_PT_Arrabida_04VV18_4
IBE_PT_Arrabida_04VV18_1
IBE_PT_Arrabida_04VV18_7
IBE_PT_Arrabida_04VV18_2
IBE_ES_MonteradelTorero_10VV18_1
IBE_ES_MonteradelTorero_10VV18_3
IBE_ES_MonteradelTorero_10VV18_5
IBE_ES_MonteradelTorero_10VV18_6
IBE_ES_MonteradelTorero_10VV18_8
IBE_PT_SaoMArcosAlferce_06VV18
IBE_PT_CanutoSPicoFoia_08VV18_5
IBE_PT_MonchiqueSPicoFoia_07VV18
IBE_PT_CanutoSPicoFoia_08VV18_1
IBE_PT_PicoFoia_09VV18_2
IBE_PT_PicoFoia_09VV18_1
MAD_PT_FaialSantana_01VV08_3
MAD_PT_Santa_05VV08_2
MAD_PT_Santa_05VV08_4
MAD_PT_Santa_05VV08_6
MAD_PT_Santa_05VV08_5
MAD_PT_SaoJorgeSaoVicente_04VV08_2
MAD_PT_FaialSantana_01VV08_2
MAD_PT_Funchal_9818MV
MAD_PT_FaialSantana_01VV08_1
MAD_PT_Santa_05VV08_3
MAD_PT_SaoJorgeSaoVicente_04VV08_4
MAD_PT_DasQueimadas_03VV08_1
MAD_PT_DasQueimadas_03VV08_3
MAD_PT_SaoJorgeSaoVicente_04VV08_3
MAD_PT_DasQueimadas_03VV08_2
MAD_PT_Funchal_3394CN
MAD_PT_CaboPontadoPargo_3
MAD_PT_CaboPontadoPargo_1
MAD_PT_CaboPontadoPargo_2
MAD_PT_SaoJorgeSaoVicente_04VV08_1
MAD_PT_CaboPontadoPargo_4
c85m15p2r1 (RAxML)
6.0E-4

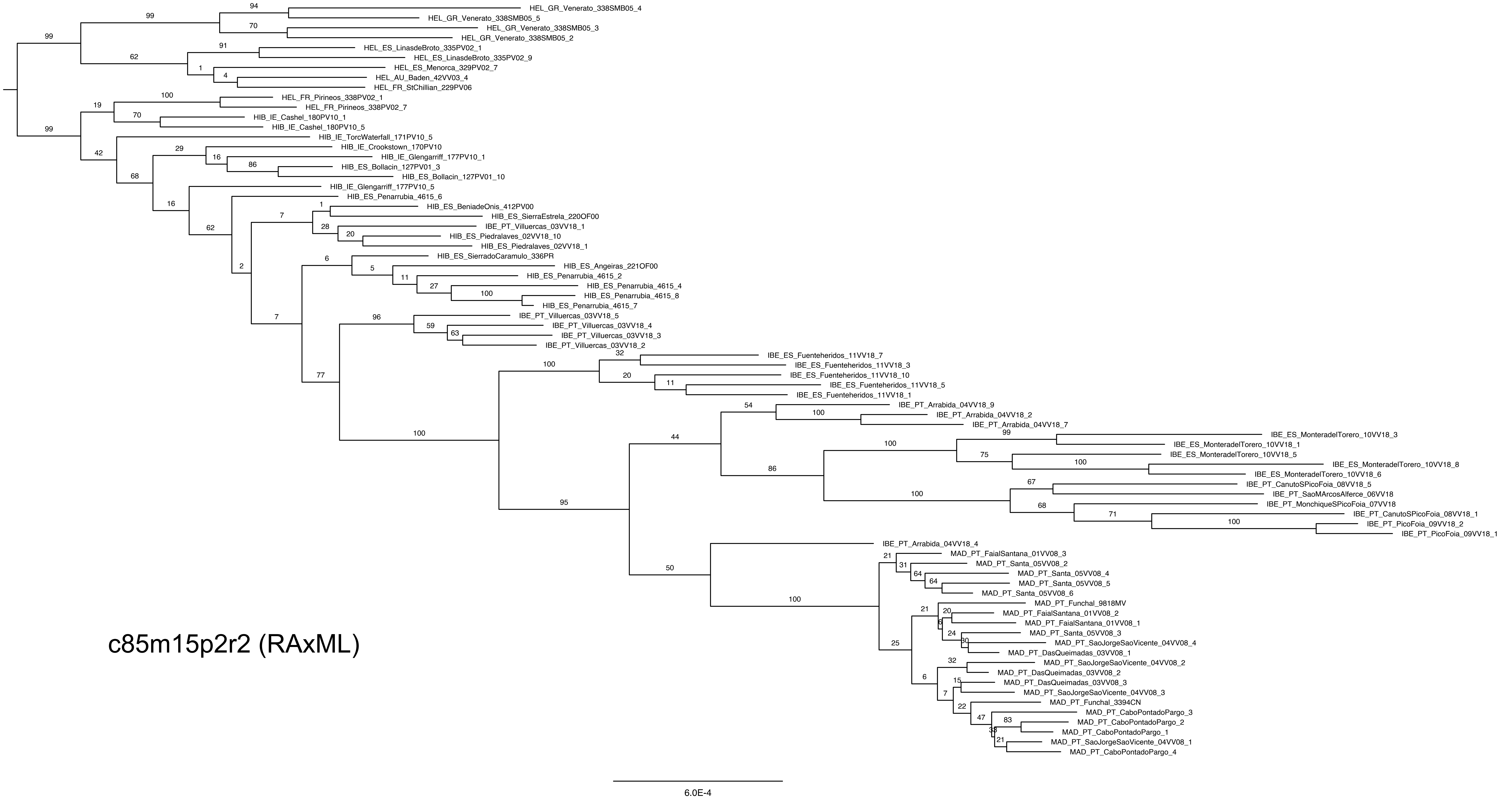

HEL_GR_Venerato_338SMB05_4
HEL_GR_Venerato_338SMB05_5
HEL_GR_Venerato_338SMB05_3
HEL_GR_Venerato_338SMB05_2
HEL_ES_LinasdeBroto_335PV02_1
HEL_ES_LinasdeBroto_335PV02_9
HEL_ES_Menorca_329PV02_7
HEL_AU_Baden_42VV03_4
HEL_FR_StChillian_229PV06
HEL_FR_Pirineos_338PV02_1
HEL_FR_Pirineos_338PV02_7
HIB_IE_Cashel_180PV10_1
HIB_IE_Cashel_180PV10_5
HIB_IE_TorcWaterfall_171PV10_5
HIB_IE_Crookstown_170PV10
HIB_IE_Glengarriff_177PV10_1
HIB_ES_Bollacin_127PV01_3
HIB_ES_Bollacin_127PV01_10
HIB_IE_Glengarriff_177PV10_5
HIB_ES_Penarrubia_4615_6
HIB_ES_BeniadeOnis_412PV00
HIB_ES_SierraEstrela_220OF00
IBE_PT_Villuercas_03VV18_1
HIB_ES_Piedralaves_02VV18_10
HIB_ES_Piedralaves_02VV18_1
HIB_ES_SierradoCaramulo_336PR
HIB_ES_Angeiras_221OF00
HIB_ES_Penarrubia_4615_2
HIB_ES_Penarrubia_4615_4
HIB_ES_Penarrubia_4615_8
HIB_ES_Penarrubia_4615_7
IBE_PT_Villuercas_03VV18_5
IBE_PT_Villuercas_03VV18_4
IBE_PT_Villuercas_03VV18_3
IBE_PT_Villuercas_03VV18_2
IBE_ES_Fuenteheridos_11VV18_7
IBE_ES_Fuenteheridos_11VV18_3
IBE_ES_Fuenteheridos_11VV18_10
IBE_ES_Fuenteheridos_11VV18_5
IBE_ES_Fuenteheridos_11VV18_1
IBE_PT_Arrabida_04VV18_9
IBE_PT_Arrabida_04VV18_2
IBE_PT_Arrabida_04VV18_7
IBE_ES_MonteradelTorero_10VV18_3
IBE_ES_MonteradelTorero_10VV18_1
IBE_ES_MonteradelTorero_10VV18_5
IBE_ES_MonteradelTorero_10VV18_8
IBE_ES_MonteradelTorero_10VV18_6
IBE_PT_CanutoSPicoFoia_08VV18_5
IBE_PT_SaoMArcosAlferce_06VV18
IBE_PT_MonchiqueSPicoFoia_07VV18
IBE_PT_CanutoSPicoFoia_08VV18_1
IBE_PT_PicoFoia_09VV18_2
IBE_PT_PicoFoia_09VV18_1
IBE_PT_Arrabida_04VV18_4
MAD_PT_FaialSantana_01VV08_3
MAD_PT_Santa_05VV08_2
MAD_PT_Santa_05VV08_4
MAD_PT_Santa_05VV08_5
MAD_PT_Santa_05VV08_6
MAD_PT_Funchal_9818MV
MAD_PT_FaialSantana_01VV08_2
MAD_PT_FaialSantana_01VV08_1
MAD_PT_Santa_05VV08_3
MAD_PT_SaoJorgeSaoVicente_04VV08_4
MAD_PT_DasQueimadas_03VV08_1
MAD_PT_SaoJorgeSaoVicente_04VV08_2
MAD_PT_DasQueimadas_03VV08_2
MAD_PT_DasQueimadas_03VV08_3
MAD_PT_SaoJorgeSaoVicente_04VV08_3
MAD_PT_Funchal_3394CN
MAD_PT_CaboPontadoPargo_3
MAD_PT_CaboPontadoPargo_2
MAD_PT_CaboPontadoPargo_1
MAD_PT_SaoJorgeSaoVicente_04VV08_1
MAD_PT_CaboPontadoPargo_4
c85m15p2r2 (RAxML)
6.0E-4

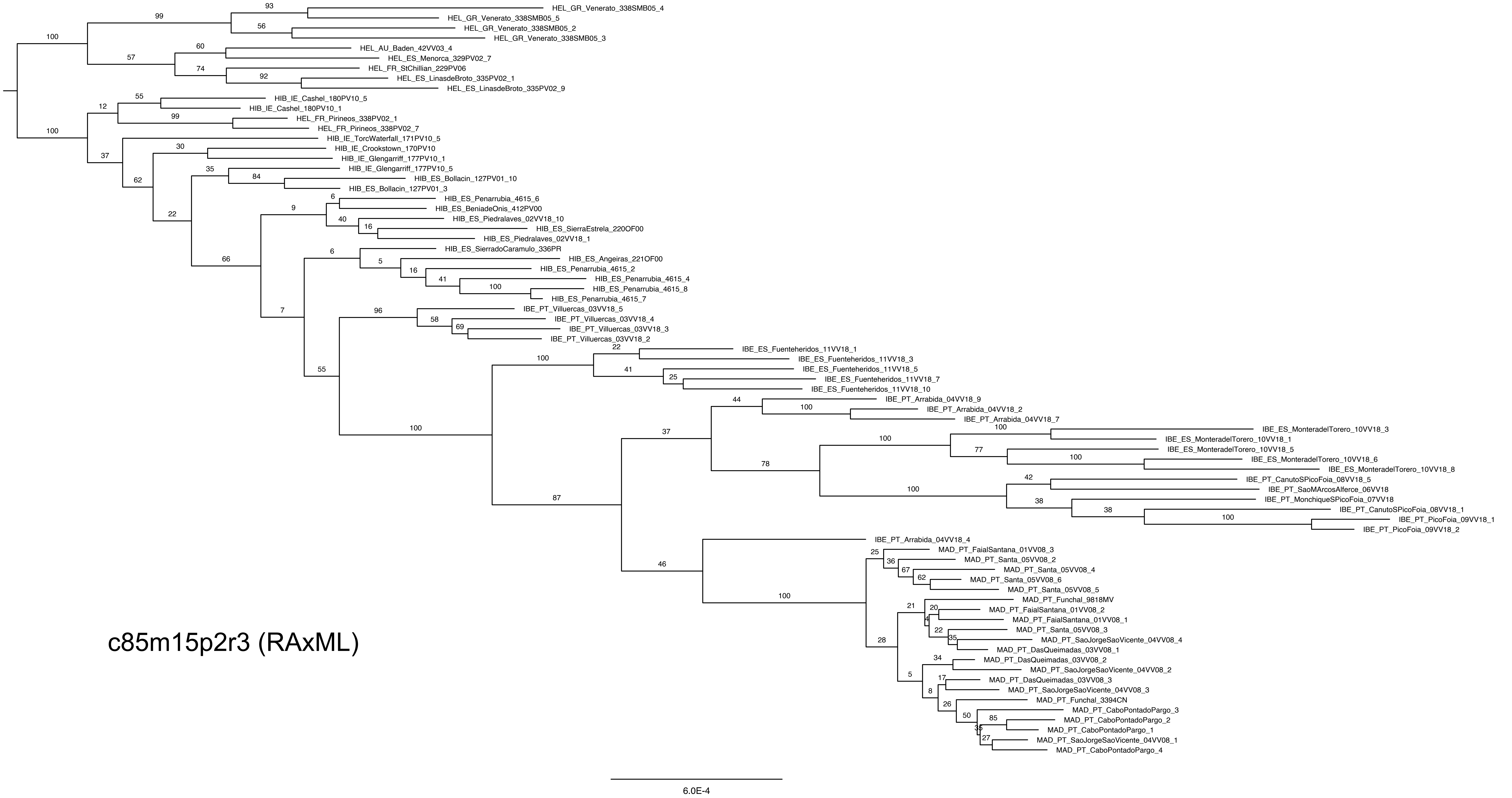


c85m15p2r3 (RAxML)

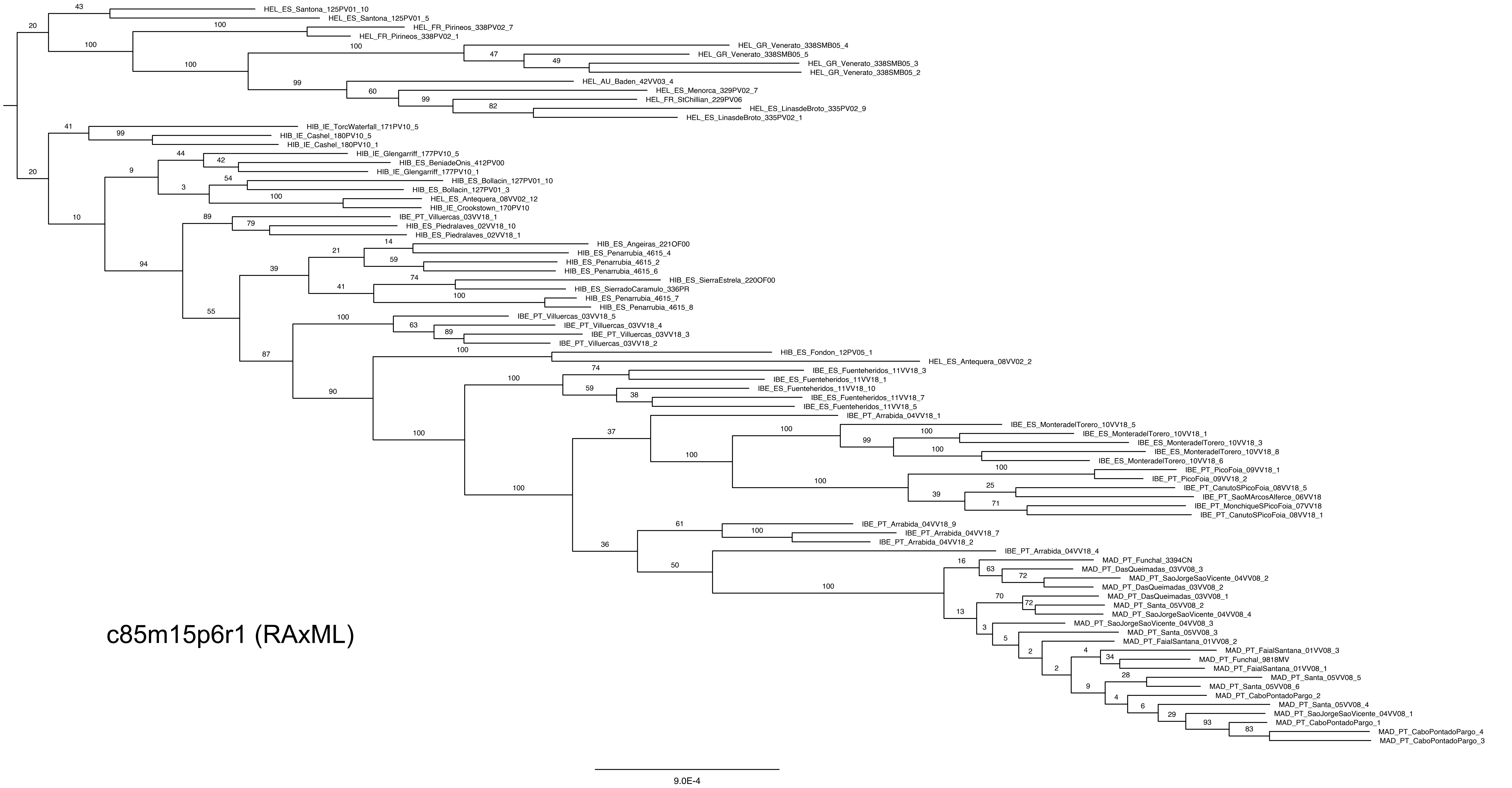
HEL_ES_Santona_125PV01_10
HEL_ES_Santona_125PV01_5
HEL_FR_Pirineos_338PV02_7
HEL_FR_Pirineos_338PV02_1
HEL_GR_Venerato_338SMB05_4
HEL_GR_Venerato_338SMB05_5
HEL_GR_Venerato_338SMB05_3
HEL_GR_Venerato_338SMB05_2
HEL_AU_Baden_42VV03_4
HEL_ES_Menorca_329PV02_7
HEL_FR_StChillian_229PV06
HEL_ES_LinasdeBroto_335PV02_9
HEL_ES_LinasdeBroto_335PV02_1
HIB_IE_TorcWaterfall_171PV10_5
HIB_IE_Cashel_180PV10_5
HIB_IE_Cashel_180PV10_1
HIB_IE_Glengarriff_177PV10_5
HIB_ES_BeniadeOnis_412PV00
HIB_IE_Glengarriff_177PV10_1
HIB_ES_Bollacin_127PV01_10
HIB_ES_Bollacin_127PV01_3
HEL_ES_Antequera_08VV02_12
HIB_IE_Crookstown_170PV10
IBE_PT_Villuercas_03VV18_1
HIB_ES_Piedralaves_02VV18_10
HIB_ES_Piedralaves_02VV18_1
HIB_ES_Angeiras_221OF00
HIB_ES_Penarrubia_4615_4
HIB_ES_Penarrubia_4615_2
HIB_ES_Penarrubia_4615_6
HIB_ES_SierraEstrela_220OF00
HIB_ES_SierradoCaramulo_336PR
HIB_ES_Penarrubia_4615_7
HIB_ES_Penarrubia_4615_8
IBE_PT_Villuercas_03VV18_5
IBE_PT_Villuercas_03VV18_4
IBE_PT_Villuercas_03VV18_3
IBE_PT_Villuercas_03VV18_2
HIB_ES_Fondon_12PV05_1
HEL_ES_Antequera_08VV02_2
IBE_ES_Fuenteheridos_11VV18_3
IBE_ES_Fuenteheridos_11VV18_1
IBE_ES_Fuenteheridos_11VV18_10
IBE_ES_Fuenteheridos_11VV18_7
IBE_ES_Fuenteheridos_11VV18_5
IBE_PT_Arrabida_04VV18_1
IBE_ES_MonteradelTorero_10VV18_5
IBE_ES_MonteradelTorero_10VV18_1
IBE_ES_MonteradelTorero_10VV18_3
IBE_ES_MonteradelTorero_10VV18_8
IBE_ES_MonteradelTorero_10VV18_6
IBE_PT_PicoFoia_09VV18_1
IBE_PT_PicoFoia_09VV18_2
IBE_PT_CanutoSPicoFoia_08VV18_5
IBE_PT_SaoMArcosAlferce_06VV18
IBE_PT_MonchiqueSPicoFoia_07VV18
IBE_PT_CanutoSPicoFoia_08VV18_1
IBE_PT_Arrabida_04VV18_9
IBE_PT_Arrabida_04VV18_7
IBE_PT_Arrabida_04VV18_2
IBE_PT_Arrabida_04VV18_4
MAD_PT_Funchal_3394CN
MAD_PT_DasQueimadas_03VV08_3
MAD_PT_SaoJorgeSaoVicente_04VV08_2
MAD_PT_DasQueimadas_03VV08_2
MAD_PT_DasQueimadas_03VV08_1
MAD_PT_Santa_05VV08_2
MAD_PT_SaoJorgeSaoVicente_04VV08_4
MAD_PT_SaoJorgeSaoVicente_04VV08_3
MAD_PT_Santa_05VV08_3
MAD_PT_FaialSantana_01VV08_2
MAD_PT_FaialSantana_01VV08_3
MAD_PT_Funchal_9818MV
MAD_PT_FaialSantana_01VV08_1
MAD_PT_Santa_05VV08_5
MAD_PT_Santa_05VV08_6
MAD_PT_CaboPontadoPargo_2
MAD_PT_Santa_05VV08_4
MAD_PT_SaoJorgeSaoVicente_04VV08_1
MAD_PT_CaboPontadoPargo_1
MAD_PT_CaboPontadoPargo_4
MAD_PT_CaboPontadoPargo_3
c85m15p6r1 (RAxML)
9.0E-4

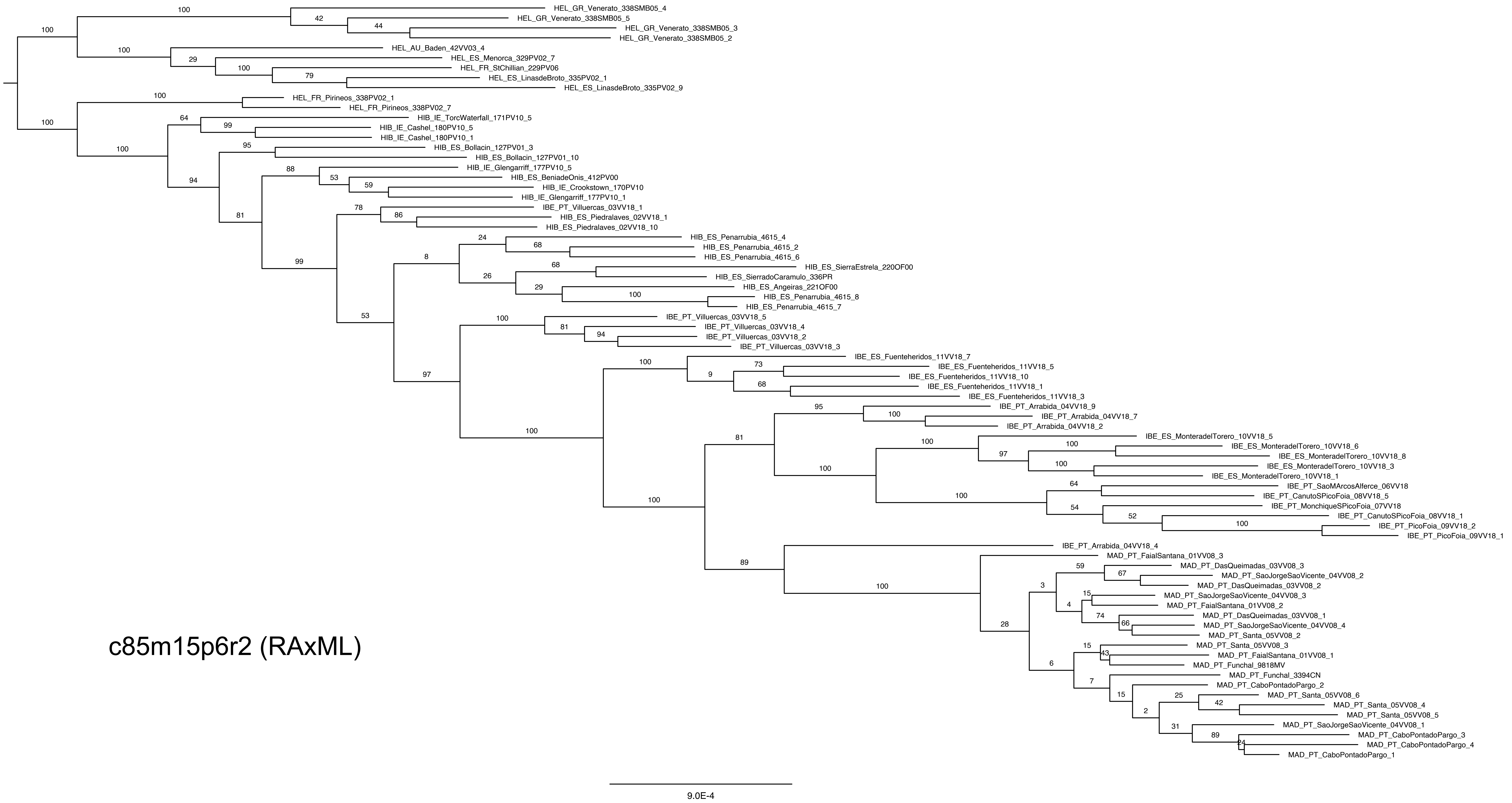
HEL_GR_Venerato_338SMB05_4
HEL_GR_Venerato_338SMB05_5
HEL_GR_Venerato_338SMB05_3
HEL_GR_Venerato_338SMB05_2
HEL_AU_Baden_42VV03_4
HEL_ES_Menorca_329PV02_7
HEL_FR_StChillian_229PV06
HEL_ES_LinasdeBroto_335PV02_1
HEL_ES_LinasdeBroto_335PV02_9
HEL_FR_Pirineos_338PV02_1
HEL_FR_Pirineos_338PV02_7
HIB_IE_TorcWaterfall_171PV10_5
HIB_IE_Cashel_180PV10_5
HIB_IE_Cashel_180PV10_1
HIB_ES_Bollacin_127PV01_3
HIB_ES_Bollacin_127PV01_10
HIB_IE_Glengarriff_177PV10_5
HIB_ES_BeniadeOnis_412PV00
HIB_IE_Crookstown_170PV10
HIB_IE_Glengarriff_177PV10_1
IBE_PT_Villuercas_03VV18_1
HIB_ES_Piedralaves_02VV18_1
HIB_ES_Piedralaves_02VV18_10
HIB_ES_Penarrubia_4615_4
HIB_ES_Penarrubia_4615_2
HIB_ES_Penarrubia_4615_6
HIB_ES_SierraEstrela_220OF00
HIB_ES_SierradoCaramulo_336PR
HIB_ES_Angeiras_221OF00
HIB_ES_Penarrubia_4615_8
HIB_ES_Penarrubia_4615_7
IBE_PT_Villuercas_03VV18_5
IBE_PT_Villuercas_03VV18_4
IBE_PT_Villuercas_03VV18_2
IBE_PT_Villuercas_03VV18_3
IBE_ES_Fuenteheridos_11VV18_7
IBE_ES_Fuenteheridos_11VV18_5
IBE_ES_Fuenteheridos_11VV18_10
IBE_ES_Fuenteheridos_11VV18_1
IBE_ES_Fuenteheridos_11VV18_3
IBE_PT_Arrabida_04VV18_9
IBE_PT_Arrabida_04VV18_7
IBE_PT_Arrabida_04VV18_2
IBE_ES_MonteradelTorero_10VV18_5
IBE_ES_MonteradelTorero_10VV18_6
IBE_ES_MonteradelTorero_10VV18_8
IBE_ES_MonteradelTorero_10VV18_3
IBE_ES_MonteradelTorero_10VV18_1
IBE_PT_SaoMArcosAlferce_06VV18
IBE_PT_CanutoSPicoFoia_08VV18_5
IBE_PT_MonchiqueSPicoFoia_07VV18
IBE_PT_CanutoSPicoFoia_08VV18_1
IBE_PT_PicoFoia_09VV18_2
IBE_PT_PicoFoia_09VV18_1
IBE_PT_Arrabida_04VV18_4
MAD_PT_FaialSantana_01VV08_3
MAD_PT_DasQueimadas_03VV08_3
MAD_PT_SaoJorgeSaoVicente_04VV08_2
MAD_PT_DasQueimadas_03VV08_2
MAD_PT_SaoJorgeSaoVicente_04VV08_3
MAD_PT_FaialSantana_01VV08_2
MAD_PT_DasQueimadas_03VV08_1
MAD_PT_SaoJorgeSaoVicente_04VV08_4
MAD_PT_Santa_05VV08_2
MAD_PT_Santa_05VV08_3
MAD_PT_FaialSantana_01VV08_1
MAD_PT_Funchal_9818MV
MAD_PT_Funchal_3394CN
MAD_PT_CaboPontadoPargo_2
MAD_PT_Santa_05VV08_6
MAD_PT_Santa_05VV08_4
MAD_PT_Santa_05VV08_5
MAD_PT_SaoJorgeSaoVicente_04VV08_1
MAD_PT_CaboPontadoPargo_3
MAD_PT_CaboPontadoPargo_4
MAD_PT_CaboPontadoPargo_1
c85m15p6r2 (RAxML)
9.0E-4

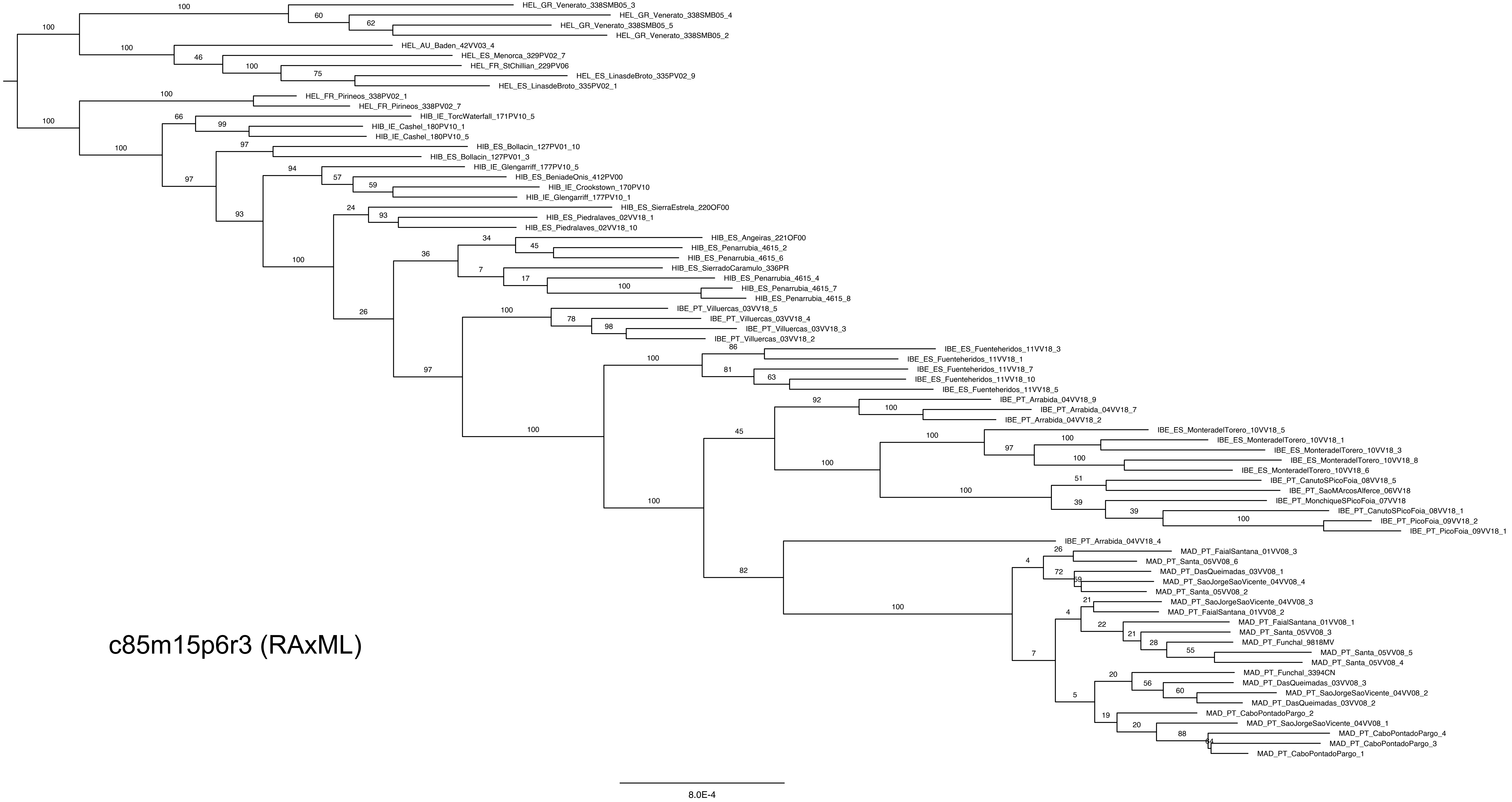
HEL_GR_Venerato_338SMB05_3
HEL_GR_Venerato_338SMB05_4
HEL_GR_Venerato_338SMB05_5
HEL_GR_Venerato_338SMB05_2
HEL_AU_Baden_42VV03_4
HEL_ES_Menorca_329PV02_7
HEL_FR_StChillian_229PV06
HEL_ES_LinasdeBroto_335PV02_9
HEL_ES_LinasdeBroto_335PV02_1
HEL_FR_Pirineos_338PV02_1
HEL_FR_Pirineos_338PV02_7
HIB_IE_TorcWaterfall_171PV10_5
HIB_IE_Cashel_180PV10_1
HIB_IE_Cashel_180PV10_5
HIB_ES_Bollacin_127PV01_10
HIB_ES_Bollacin_127PV01_3
HIB_IE_Glengarriff_177PV10_5
HIB_ES_BeniadeOnis_412PV00
HIB_IE_Crookstown_170PV10
HIB_IE_Glengarriff_177PV10_1
HIB_ES_SierraEstrela_220OF00
HIB_ES_Piedralaves_02VV18_1
HIB_ES_Piedralaves_02VV18_10
HIB_ES_Angeiras_221OF00
HIB_ES_Penarrubia_4615_2
HIB_ES_Penarrubia_4615_6
HIB_ES_SierradoCaramulo_336PR
HIB_ES_Penarrubia_4615_4
HIB_ES_Penarrubia_4615_7
HIB_ES_Penarrubia_4615_8
IBE_PT_Villuercas_03VV18_5
IBE_PT_Villuercas_03VV18_4
IBE_PT_Villuercas_03VV18_3
IBE_PT_Villuercas_03VV18_2
IBE_ES_Fuenteheridos_11VV18_3
IBE_ES_Fuenteheridos_11VV18_1
IBE_ES_Fuenteheridos_11VV18_7
IBE_ES_Fuenteheridos_11VV18_10
IBE_ES_Fuenteheridos_11VV18_5
IBE_PT_Arrabida_04VV18_9
IBE_PT_Arrabida_04VV18_7
IBE_PT_Arrabida_04VV18_2
IBE_ES_MonteradelTorero_10VV18_5
IBE_ES_MonteradelTorero_10VV18_1
IBE_ES_MonteradelTorero_10VV18_3
IBE_ES_MonteradelTorero_10VV18_8
IBE_ES_MonteradelTorero_10VV18_6
IBE_PT_CanutoSPicoFoia_08VV18_5
IBE_PT_SaoMArcosAlferce_06VV18
IBE_PT_MonchiqueSPicoFoia_07VV18
IBE_PT_CanutoSPicoFoia_08VV18_1
IBE_PT_PicoFoia_09VV18_2
IBE_PT_PicoFoia_09VV18_1
IBE_PT_Arrabida_04VV18_4
MAD_PT_FaialSantana_01VV08_3
MAD_PT_Santa_05VV08_6
MAD_PT_DasQueimadas_03VV08_1
MAD_PT_SaoJorgeSaoVicente_04VV08_4
MAD_PT_Santa_05VV08_2
MAD_PT_SaoJorgeSaoVicente_04VV08_3
MAD_PT_FaialSantana_01VV08_2
MAD_PT_FaialSantana_01VV08_1
MAD_PT_Santa_05VV08_3
MAD_PT_Funchal_9818MV
MAD_PT_Santa_05VV08_5
MAD_PT_Santa_05VV08_4
MAD_PT_Funchal_3394CN
MAD_PT_DasQueimadas_03VV08_3
MAD_PT_SaoJorgeSaoVicente_04VV08_2
MAD_PT_DasQueimadas_03VV08_2
MAD_PT_CaboPontadoPargo_2
MAD_PT_SaoJorgeSaoVicente_04VV08_1
MAD_PT_CaboPontadoPargo_4
MAD_PT_CaboPontadoPargo_3
MAD_PT_CaboPontadoPargo_1
c85m15p6r3 (RAxML)
8.0E-4

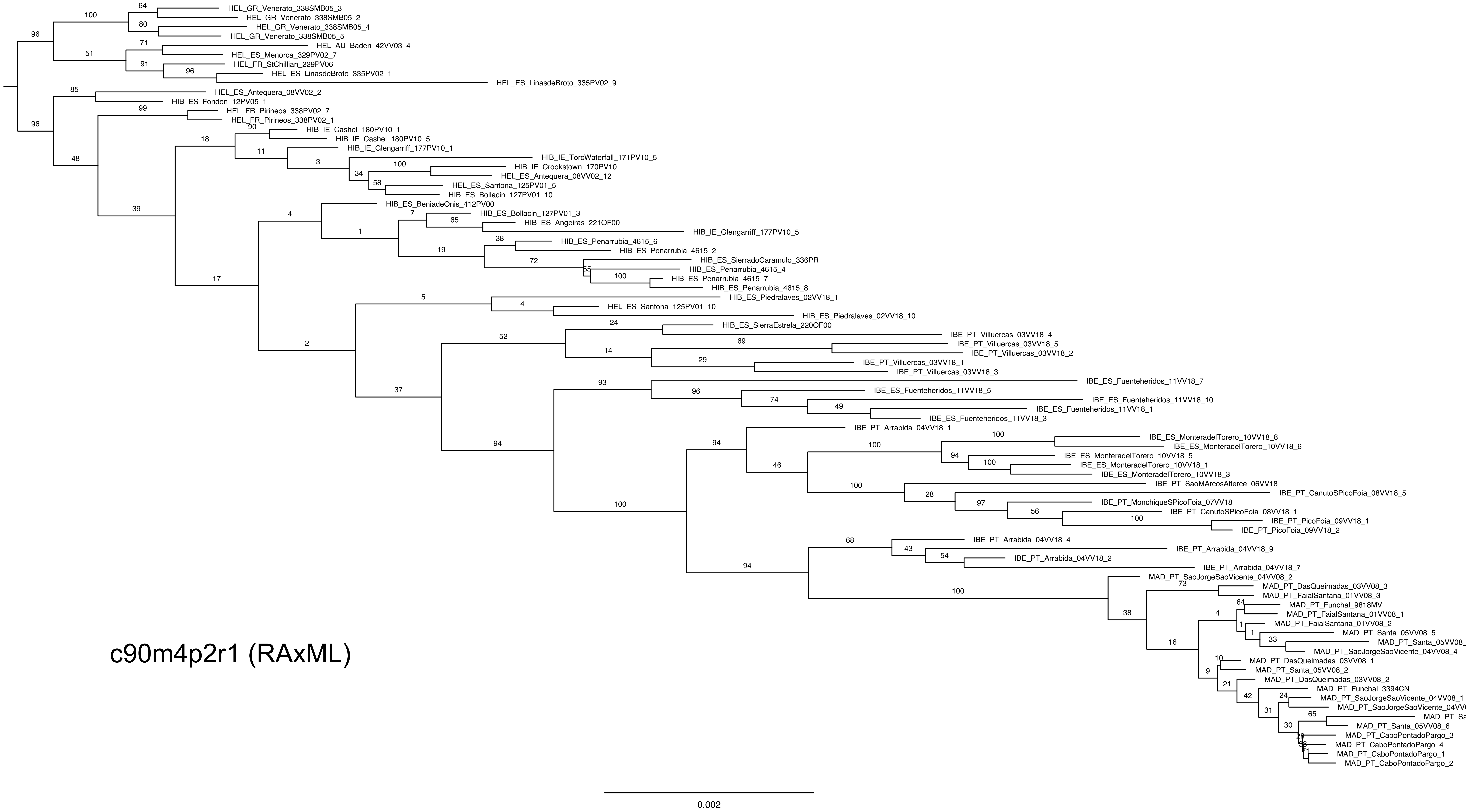
HEL_GR_Venerato_338SMB05_3
HEL_GR_Venerato_338SMB05_2
HEL_GR_Venerato_338SMB05_4
HEL_GR_Venerato_338SMB05_5
HEL_AU_Baden_42VV03_4
HEL_ES_Menorca_329PV02_7
HEL_FR_StChillian_229PV06
HEL_ES_LinasdeBroto_335PV02_1
HEL_ES_LinasdeBroto_335PV02_9
HEL_ES_Antequera_08VV02_2
HIB_ES_Fondon_12PV05_1
HEL_FR_Pirineos_338PV02_7
HEL_FR_Pirineos_338PV02_1
HIB_IE_Cashel_180PV10_1
HIB_IE_Cashel_180PV10_5
HIB_IE_Glengarriff_177PV10_1
HIB_IE_TorcWaterfall_171PV10_5
HIB_IE_Crookstown_170PV10
HEL_ES_Antequera_08VV02_12
HEL_ES_Santona_125PV01_5
HIB_ES_Bollacin_127PV01_10
HIB_ES_BeniadeOnis_412PV00
HIB_ES_Bollacin_127PV01_3
HIB_ES_Angeiras_221OF00
HIB_IE_Glengarriff_177PV10_5
HIB_ES_Penarrubia_4615_6
HIB_ES_Penarrubia_4615_2
HIB_ES_SierradoCaramulo_336PR
HIB_ES_Penarrubia_4615_4
HIB_ES_Penarrubia_4615_7
HIB_ES_Penarrubia_4615_8
HIB_ES_Piedralaves_02VV18_1
HEL_ES_Santona_125PV01_10
HIB_ES_Piedralaves_02VV18_10
HIB_ES_SierraEstrela_220OF00
IBE_PT_Villuercas_03VV18_4
IBE_PT_Villuercas_03VV18_5
IBE_PT_Villuercas_03VV18_2
IBE_PT_Villuercas_03VV18_1
IBE_PT_Villuercas_03VV18_3
IBE_ES_Fuenteheridos_11VV18_7
IBE_ES_Fuenteheridos_11VV18_5
IBE_ES_Fuenteheridos_11VV18_10
IBE_ES_Fuenteheridos_11VV18_1
IBE_ES_Fuenteheridos_11VV18_3
IBE_PT_Arrabida_04VV18_1
IBE_ES_MonteradelTorero_10VV18_8
IBE_ES_MonteradelTorero_10VV18_6
IBE_ES_MonteradelTorero_10VV18_5
IBE_ES_MonteradelTorero_10VV18_1
IBE_ES_MonteradelTorero_10VV18_3
IBE_PT_SaoMArcosAlferce_06VV18
IBE_PT_CanutoSPicoFoia_08VV18_5
IBE_PT_MonchiqueSPicoFoia_07VV18
IBE_PT_CanutoSPicoFoia_08VV18_1
IBE_PT_PicoFoia_09VV18_1
IBE_PT_PicoFoia_09VV18_2
IBE_PT_Arrabida_04VV18_4
IBE_PT_Arrabida_04VV18_9
IBE_PT_Arrabida_04VV18_2
IBE_PT_Arrabida_04VV18_7
MAD_PT_SaoJorgeSaoVicente_04VV08_2
MAD_PT_DasQueimadas_03VV08_3
MAD_PT_FaialSantana_01VV08_3
MAD_PT_Funchal_9818MV
MAD_PT_FaialSantana_01VV08_1
MAD_PT_FaialSantana_01VV08_2
MAD_PT_Santa_05VV08_5
MAD_PT_Santa_05VV08_3
MAD_PT_SaoJorgeSaoVicente_04VV08_4
MAD_PT_DasQueimadas_03VV08_1
MAD_PT_Santa_05VV08_2
MAD_PT_DasQueimadas_03VV08_2
MAD_PT_Funchal_3394CN
MAD_PT_SaoJorgeSaoVicente_04VV08_1
MAD_PT_SaoJorgeSaoVicente_04VV08_3
MAD_PT_Santa_05VV08_4
MAD_PT_Santa_05VV08_6
MAD_PT_CaboPontadoPargo_3
MAD_PT_CaboPontadoPargo_4
MAD_PT_CaboPontadoPargo_1
MAD_PT_CaboPontadoPargo_2
c90m4p2r1 (RAxML)
0.002

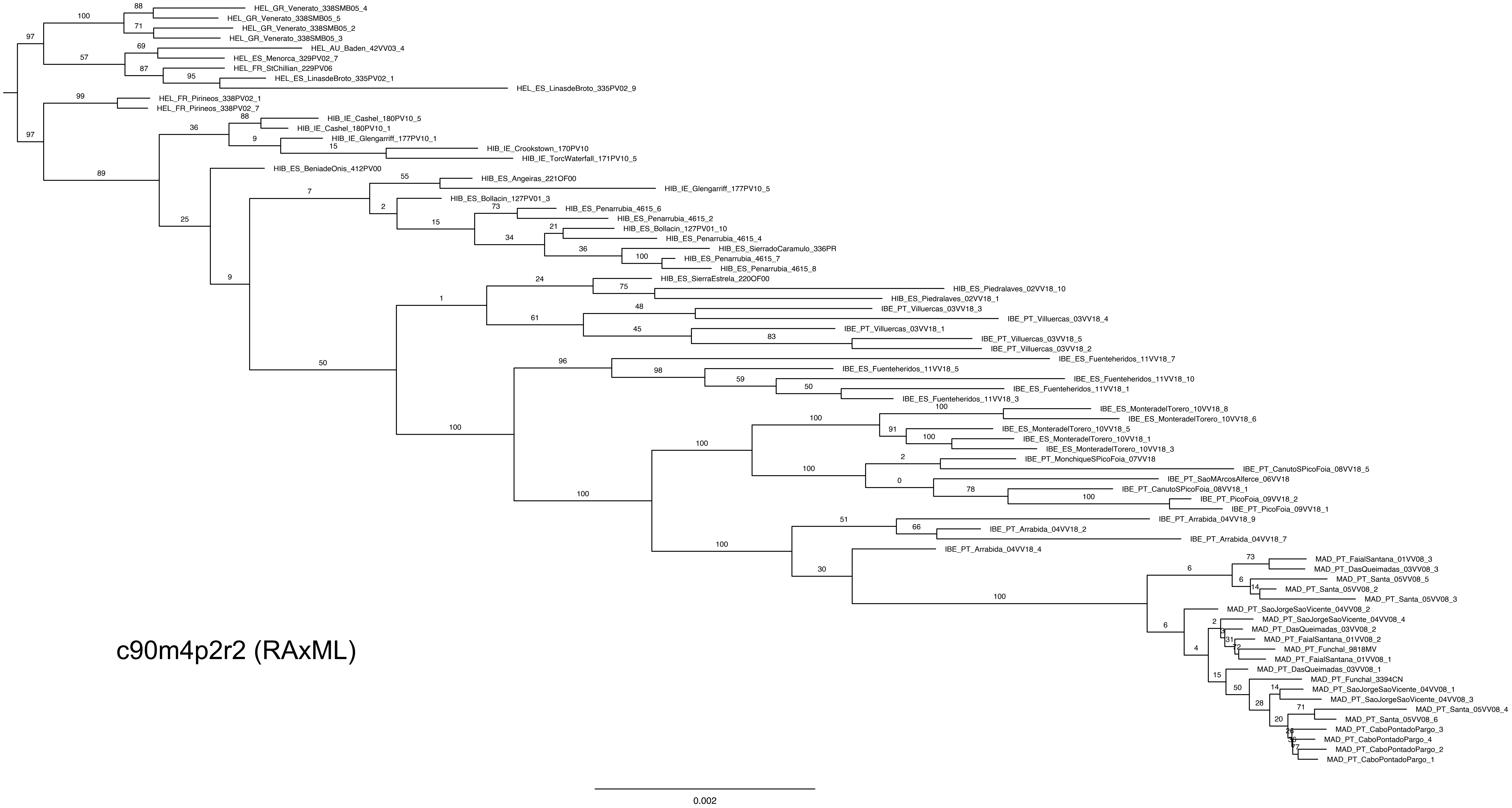

HEL_GR_Venerato_338SMB05_4
HEL_GR_Venerato_338SMB05_5
HEL_GR_Venerato_338SMB05_2
HEL_GR_Venerato_338SMB05_3
HEL_AU_Baden_42VV03_4
HEL_ES_Menorca_329PV02_7
HEL_FR_StChillian_229PV06
HEL_ES_LinasdeBroto_335PV02_1
HEL_ES_LinasdeBroto_335PV02_9
HEL_FR_Pirineos_338PV02_1
HEL_FR_Pirineos_338PV02_7
HIB_IE_Cashel_180PV10_5
HIB_IE_Cashel_180PV10_1
HIB_IE_Glengarriff_177PV10_1
HIB_IE_Crookstown_170PV10
HIB_IE_TorcWaterfall_171PV10_5
HIB_ES_BeniadeOnis_412PV00
HIB_ES_Angeiras_221OF00
HIB_IE_Glengarriff_177PV10_5
HIB_ES_Bollacin_127PV01_3
HIB_ES_Penarrubia_4615_6
HIB_ES_Penarrubia_4615_2
HIB_ES_Bollacin_127PV01_10
HIB_ES_Penarrubia_4615_4
HIB_ES_SierradoCaramulo_336PR
HIB_ES_Penarrubia_4615_7
HIB_ES_Penarrubia_4615_8
HIB_ES_SierraEstrela_220OF00
HIB_ES_Piedralaves_02VV18_10
HIB_ES_Piedralaves_02VV18_1
IBE_PT_Villuercas_03VV18_3
IBE_PT_Villuercas_03VV18_4
IBE_PT_Villuercas_03VV18_1
IBE_PT_Villuercas_03VV18_5
IBE_PT_Villuercas_03VV18_2
IBE_ES_Fuenteheridos_11VV18_7
IBE_ES_Fuenteheridos_11VV18_5
IBE_ES_Fuenteheridos_11VV18_10
IBE_ES_Fuenteheridos_11VV18_1
IBE_ES_Fuenteheridos_11VV18_3
IBE_ES_MonteradelTorero_10VV18_8
IBE_ES_MonteradelTorero_10VV18_6
IBE_ES_MonteradelTorero_10VV18_5
IBE_ES_MonteradelTorero_10VV18_1
IBE_ES_MonteradelTorero_10VV18_3
IBE_PT_MonchiqueSPicoFoia_07VV18
IBE_PT_CanutoSPicoFoia_08VV18_5
IBE_PT_SaoMArcosAlferce_06VV18
IBE_PT_CanutoSPicoFoia_08VV18_1
IBE_PT_PicoFoia_09VV18_2
IBE_PT_PicoFoia_09VV18_1
IBE_PT_Arrabida_04VV18_9
IBE_PT_Arrabida_04VV18_2
IBE_PT_Arrabida_04VV18_7
IBE_PT_Arrabida_04VV18_4
MAD_PT_FaialSantana_01VV08_3
MAD_PT_DasQueimadas_03VV08_3
MAD_PT_Santa_05VV08_5
MAD_PT_Santa_05VV08_2
MAD_PT_Santa_05VV08_3
MAD_PT_SaoJorgeSaoVicente_04VV08_2
MAD_PT_SaoJorgeSaoVicente_04VV08_4
MAD_PT_DasQueimadas_03VV08_2
MAD_PT_FaialSantana_01VV08_2
MAD_PT_Funchal_9818MV
MAD_PT_FaialSantana_01VV08_1
MAD_PT_DasQueimadas_03VV08_1
MAD_PT_Funchal_3394CN
MAD_PT_SaoJorgeSaoVicente_04VV08_1
MAD_PT_SaoJorgeSaoVicente_04VV08_3
MAD_PT_Santa_05VV08_4
MAD_PT_Santa_05VV08_6
MAD_PT_CaboPontadoPargo_3
MAD_PT_CaboPontadoPargo_4
MAD_PT_CaboPontadoPargo_2
MAD_PT_CaboPontadoPargo_1
c90m4p2r2 (RAxML)
0.002

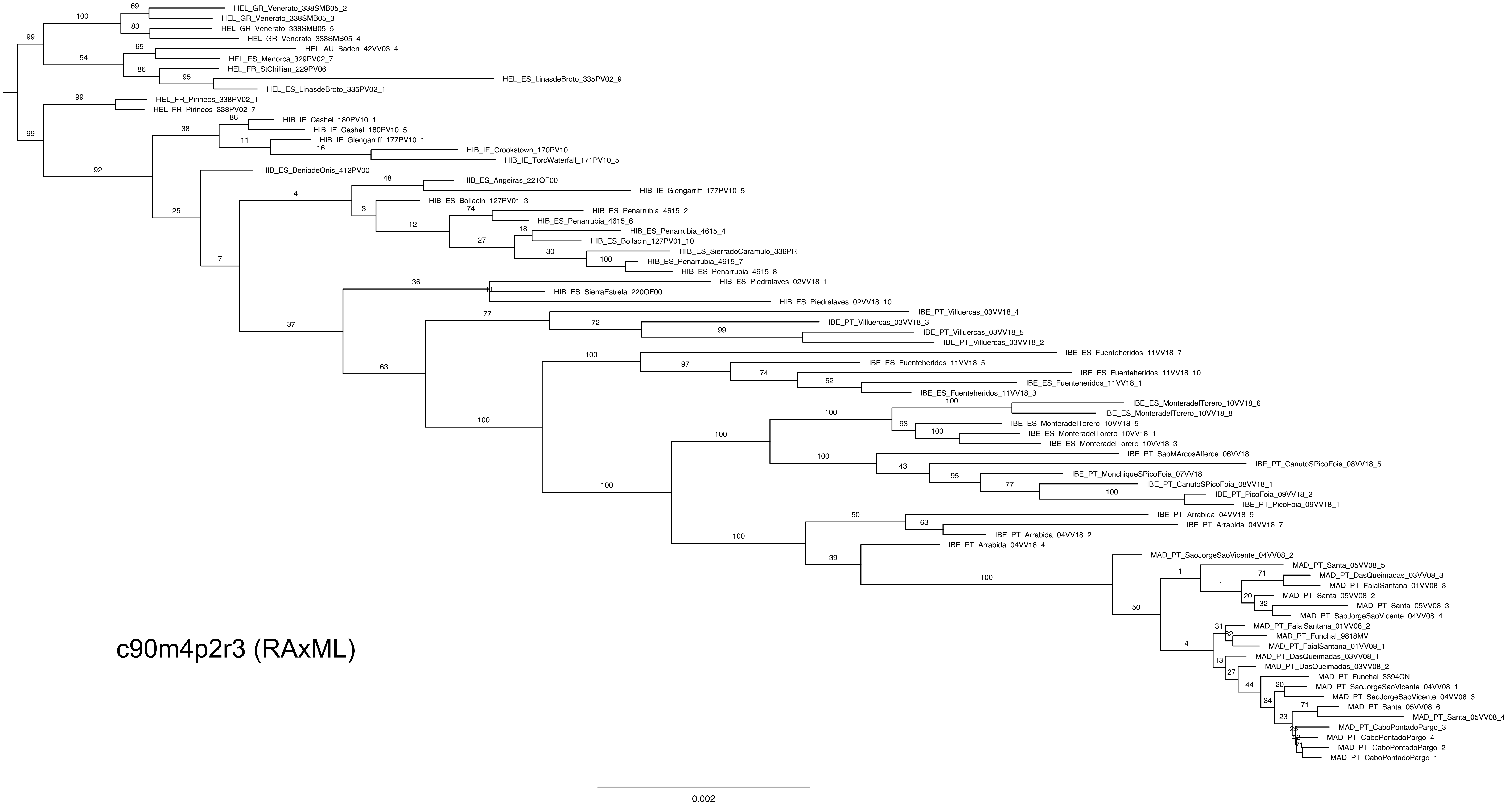
HEL_GR_Venerato_338SMB05_2
HEL_GR_Venerato_338SMB05_3
HEL_GR_Venerato_338SMB05_5
HEL_GR_Venerato_338SMB05_4
HEL_AU_Baden_42VV03_4
HEL_ES_Menorca_329PV02_7
HEL_FR_StChillian_229PV06
HEL_ES_LinasdeBroto_335PV02_9
HEL_ES_LinasdeBroto_335PV02_1
HEL_FR_Pirineos_338PV02_1
HEL_FR_Pirineos_338PV02_7
HIB_IE_Cashel_180PV10_1
HIB_IE_Cashel_180PV10_5
HIB_IE_Glengarriff_177PV10_1
HIB_IE_Crookstown_170PV10
HIB_IE_TorcWaterfall_171PV10_5
HIB_ES_BeniadeOnis_412PV00
HIB_ES_Angeiras_221OF00
HIB_IE_Glengarriff_177PV10_5
HIB_ES_Bollacin_127PV01_3
HIB_ES_Penarrubia_4615_2
HIB_ES_Penarrubia_4615_6
HIB_ES_Penarrubia_4615_4
HIB_ES_Bollacin_127PV01_10
HIB_ES_SierradoCaramulo_336PR
HIB_ES_Penarrubia_4615_7
HIB_ES_Penarrubia_4615_8
HIB_ES_Piedralaves_02VV18_1
HIB_ES_SierraEstrela_220OF00
HIB_ES_Piedralaves_02VV18_10
IBE_PT_Villuercas_03VV18_4
IBE_PT_Villuercas_03VV18_3
IBE_PT_Villuercas_03VV18_5
IBE_PT_Villuercas_03VV18_2
IBE_ES_Fuenteheridos_11VV18_7
IBE_ES_Fuenteheridos_11VV18_5
IBE_ES_Fuenteheridos_11VV18_10
IBE_ES_Fuenteheridos_11VV18_1
IBE_ES_Fuenteheridos_11VV18_3
IBE_ES_MonteradelTorero_10VV18_6
IBE_ES_MonteradelTorero_10VV18_8
IBE_ES_MonteradelTorero_10VV18_5
IBE_ES_MonteradelTorero_10VV18_1
IBE_ES_MonteradelTorero_10VV18_3
IBE_PT_SaoMArcosAlferce_06VV18
IBE_PT_CanutoSPicoFoia_08VV18_5
IBE_PT_MonchiqueSPicoFoia_07VV18
IBE_PT_CanutoSPicoFoia_08VV18_1
IBE_PT_PicoFoia_09VV18_2
IBE_PT_PicoFoia_09VV18_1
IBE_PT_Arrabida_04VV18_9
IBE_PT_Arrabida_04VV18_7
IBE_PT_Arrabida_04VV18_2
IBE_PT_Arrabida_04VV18_4
MAD_PT_SaoJorgeSaoVicente_04VV08_2
MAD_PT_Santa_05VV08_5
MAD_PT_DasQueimadas_03VV08_3
MAD_PT_FaialSantana_01VV08_3
MAD_PT_Santa_05VV08_2
MAD_PT_Santa_05VV08_3
MAD_PT_SaoJorgeSaoVicente_04VV08_4
MAD_PT_FaialSantana_01VV08_2
MAD_PT_Funchal_9818MV
MAD_PT_FaialSantana_01VV08_1
MAD_PT_DasQueimadas_03VV08_1
MAD_PT_DasQueimadas_03VV08_2
MAD_PT_Funchal_3394CN
MAD_PT_SaoJorgeSaoVicente_04VV08_1
MAD_PT_SaoJorgeSaoVicente_04VV08_3
MAD_PT_Santa_05VV08_6
MAD_PT_Santa_05VV08_4
MAD_PT_CaboPontadoPargo_3
MAD_PT_CaboPontadoPargo_4
MAD_PT_CaboPontadoPargo_2
MAD_PT_CaboPontadoPargo_1
c90m4p2r3 (RAxML)
0.002

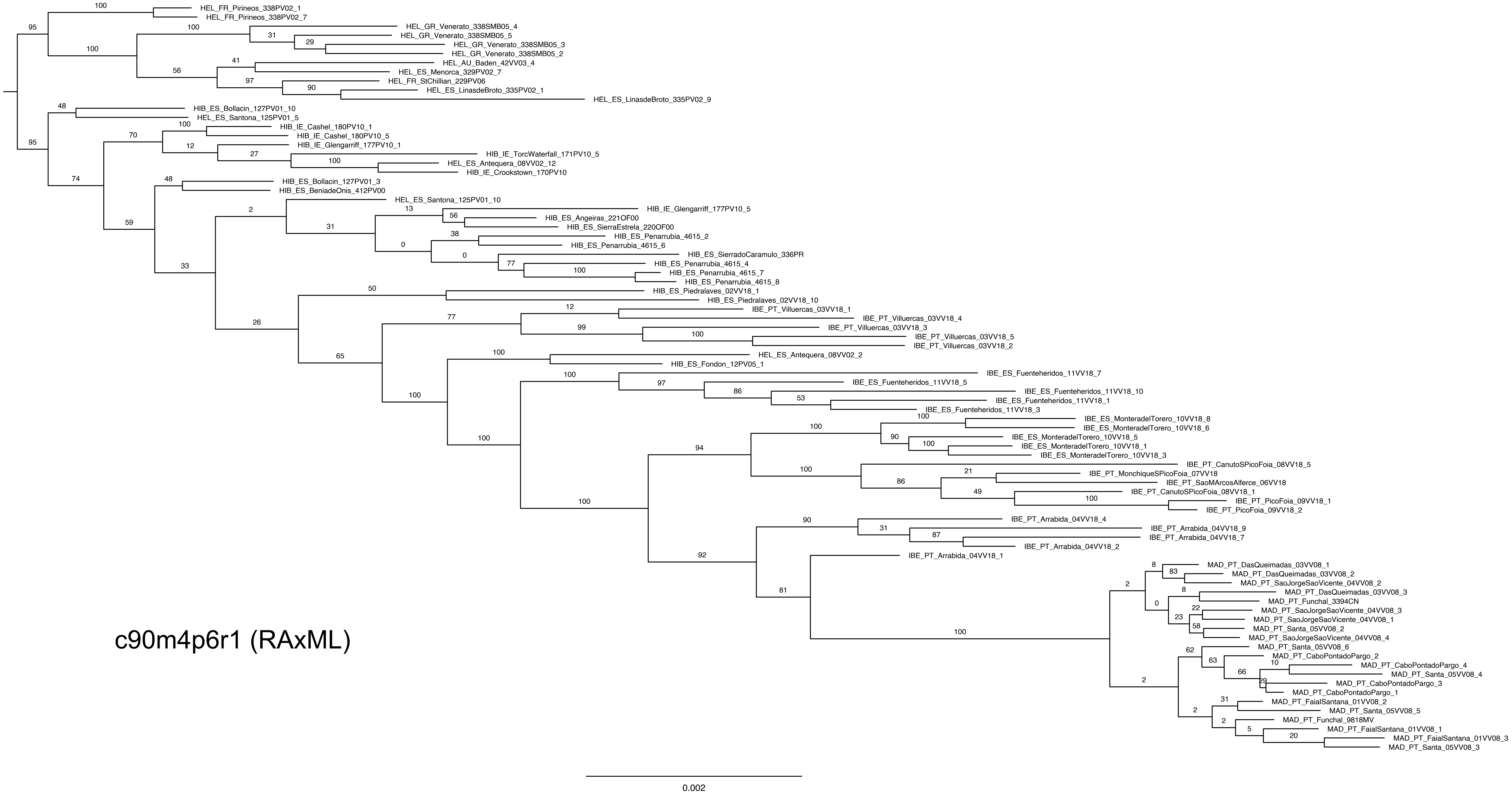
HEL_FR_Pirineos_338PV02_1
HEL_FR_Pirineos_338PV02_7
HEL_GR_Venerato_338SMB05_4
HEL_GR_Venerato_338SMB05_5
HEL_GR_Venerato_338SMB05_3
HEL_GR_Venerato_338SMB05_2
HEL_AU_Baden_42VV03_4
HEL_ES_Menorca_329PV02_7
HEL_FR_StChillian_229PV06
HEL_ES_LinasdeBroto_335PV02_1
HEL_ES_LinasdeBroto_335PV02_9
HIB_ES_Bollacin_127PV01_10
HEL_ES_Santona_125PV01_5
HIB_IE_Cashel_180PV10_1
HIB_IE_Cashel_180PV10_5
HIB_IE_Glengarriff_177PV10_1
HIB_IE_TorcWaterfall_171PV10_5
HEL_ES_Antequera_08VV02_12
HIB_IE_Crookstown_170PV10
HIB_ES_Bollacin_127PV01_3
HIB_ES_BeniadeOnis_412PV00
HEL_ES_Santona_125PV01_10
HIB_IE_Glengarriff_177PV10_5
HIB_ES_Angeiras_221OF00
HIB_ES_SierraEstrela_220OF00
HIB_ES_Penarrubia_4615_2
HIB_ES_Penarrubia_4615_6
HIB_ES_SierradoCaramulo_336PR
HIB_ES_Penarrubia_4615_4
HIB_ES_Penarrubia_4615_7
HIB_ES_Penarrubia_4615_8
HIB_ES_Piedralaves_02VV18_1
HIB_ES_Piedralaves_02VV18_10
IBE_PT_Villuercas_03VV18_1
IBE_PT_Villuercas_03VV18_4
IBE_PT_Villuercas_03VV18_3
IBE_PT_Villuercas_03VV18_5
IBE_PT_Villuercas_03VV18_2
HEL_ES_Antequera_08VV02_2
HIB_ES_Fondon_12PV05_1
IBE_ES_Fuenteheridos_11VV18_7
IBE_ES_Fuenteheridos_11VV18_5
IBE_ES_Fuenteheridos_11VV18_10
IBE_ES_Fuenteheridos_11VV18_1
IBE_ES_Fuenteheridos_11VV18_3
IBE_ES_MonteradelTorero_10VV18_8
IBE_ES_MonteradelTorero_10VV18_6
IBE_ES_MonteradelTorero_10VV18_5
IBE_ES_MonteradelTorero_10VV18_1
IBE_ES_MonteradelTorero_10VV18_3
IBE_PT_CanutoSPicoFoia_08VV18_5
IBE_PT_MonchiqueSPicoFoia_07VV18
IBE_PT_SaoMArcosAlferce_06VV18
IBE_PT_CanutoSPicoFoia_08VV18_1
IBE_PT_PicoFoia_09VV18_1
IBE_PT_PicoFoia_09VV18_2
IBE_PT_Arrabida_04VV18_4
IBE_PT_Arrabida_04VV18_9
IBE_PT_Arrabida_04VV18_7
IBE_PT_Arrabida_04VV18_2
IBE_PT_Arrabida_04VV18_1
MAD_PT_DasQueimadas_03VV08_1
MAD_PT_DasQueimadas_03VV08_2
MAD_PT_SaoJorgeSaoVicente_04VV08_2
MAD_PT_DasQueimadas_03VV08_3
MAD_PT_Funchal_3394CN
MAD_PT_SaoJorgeSaoVicente_04VV08_3
MAD_PT_SaoJorgeSaoVicente_04VV08_1
MAD_PT_Santa_05VV08_2
MAD_PT_SaoJorgeSaoVicente_04VV08_4
MAD_PT_Santa_05VV08_6
MAD_PT_CaboPontadoPargo_2
MAD_PT_CaboPontadoPargo_4
MAD_PT_Santa_05VV08_4
MAD_PT_CaboPontadoPargo_3
MAD_PT_CaboPontadoPargo_1
MAD_PT_FaialSantana_01VV08_2
MAD_PT_Santa_05VV08_5
MAD_PT_Funchal_9818MV
MAD_PT_FaialSantana_01VV08_1
MAD_PT_FaialSantana_01VV08_3
MAD_PT_Santa_05VV08_3
c90m4p6r1 (RAxML)
0.002

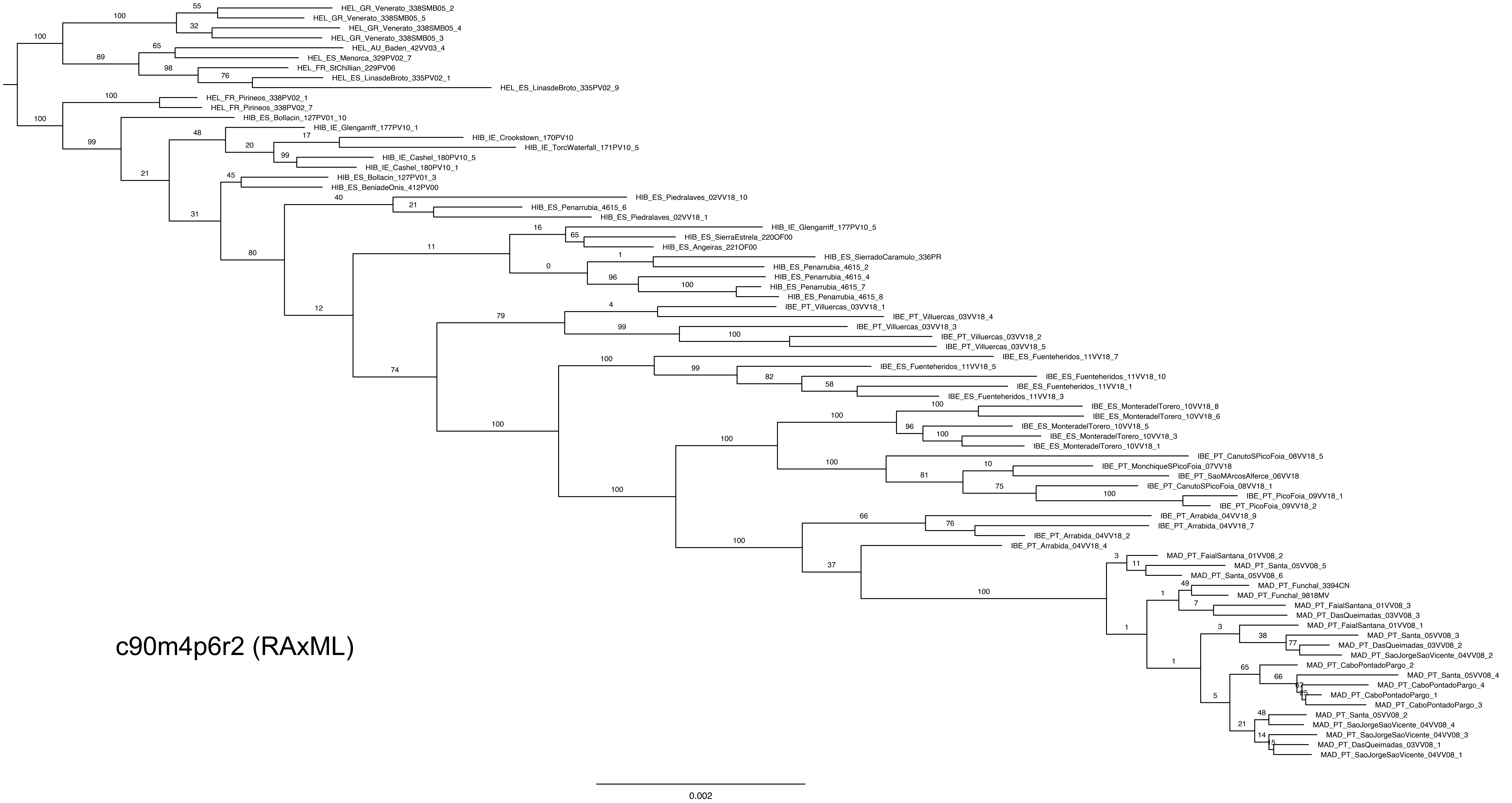
HEL_GR_Venerato_338SMB05_2
HEL_GR_Venerato_338SMB05_5
HEL_GR_Venerato_338SMB05_4
HEL_GR_Venerato_338SMB05_3
HEL_AU_Baden_42VV03_4
HEL_ES_Menorca_329PV02_7
HEL_FR_StChillian_229PV06
HEL_ES_LinasdeBroto_335PV02_1
HEL_ES_LinasdeBroto_335PV02_9
HEL_FR_Pirineos_338PV02_1
HEL_FR_Pirineos_338PV02_7
HIB_ES_Bollacin_127PV01_10
HIB_IE_Glengarriff_177PV10_1
HIB_IE_Crookstown_170PV10
HIB_IE_TorcWaterfall_171PV10_5
HIB_IE_Cashel_180PV10_5
HIB_IE_Cashel_180PV10_1
HIB_ES_Bollacin_127PV01_3
HIB_ES_BeniadeOnis_412PV00
HIB_ES_Piedralaves_02VV18_10
HIB_ES_Penarrubia_4615_6
HIB_ES_Piedralaves_02VV18_1
HIB_IE_Glengarriff_177PV10_5
HIB_ES_SierraEstrela_220OF00
HIB_ES_Angeiras_221OF00
HIB_ES_SierradoCaramulo_336PR
HIB_ES_Penarrubia_4615_2
HIB_ES_Penarrubia_4615_4
HIB_ES_Penarrubia_4615_7
HIB_ES_Penarrubia_4615_8
IBE_PT_Villuercas_03VV18_1
IBE_PT_Villuercas_03VV18_4
IBE_PT_Villuercas_03VV18_3
IBE_PT_Villuercas_03VV18_2
IBE_PT_Villuercas_03VV18_5
IBE_ES_Fuenteheridos_11VV18_7
IBE_ES_Fuenteheridos_11VV18_5
IBE_ES_Fuenteheridos_11VV18_10
IBE_ES_Fuenteheridos_11VV18_1
IBE_ES_Fuenteheridos_11VV18_3
IBE_ES_MonteradelTorero_10VV18_8
IBE_ES_MonteradelTorero_10VV18_6
IBE_ES_MonteradelTorero_10VV18_5
IBE_ES_MonteradelTorero_10VV18_3
IBE_ES_MonteradelTorero_10VV18_1
IBE_PT_CanutoSPicoFoia_08VV18_5
IBE_PT_MonchiqueSPicoFoia_07VV18
IBE_PT_SaoMArcosAlferce_06VV18
IBE_PT_CanutoSPicoFoia_08VV18_1
IBE_PT_PicoFoia_09VV18_1
IBE_PT_PicoFoia_09VV18_2
IBE_PT_Arrabida_04VV18_9
IBE_PT_Arrabida_04VV18_7
IBE_PT_Arrabida_04VV18_2
IBE_PT_Arrabida_04VV18_4
MAD_PT_FaialSantana_01VV08_2
MAD_PT_Santa_05VV08_5
MAD_PT_Santa_05VV08_6
MAD_PT_Funchal_3394CN
MAD_PT_Funchal_9818MV
MAD_PT_FaialSantana_01VV08_3
MAD_PT_DasQueimadas_03VV08_3
MAD_PT_FaialSantana_01VV08_1
MAD_PT_Santa_05VV08_3
MAD_PT_DasQueimadas_03VV08_2
MAD_PT_SaoJorgeSaoVicente_04VV08_2
MAD_PT_CaboPontadoPargo_2
MAD_PT_Santa_05VV08_4
MAD_PT_CaboPontadoPargo_4
MAD_PT_CaboPontadoPargo_1
MAD_PT_CaboPontadoPargo_3
MAD_PT_Santa_05VV08_2
MAD_PT_SaoJorgeSaoVicente_04VV08_4
MAD_PT_SaoJorgeSaoVicente_04VV08_3
MAD_PT_DasQueimadas_03VV08_1
MAD_PT_SaoJorgeSaoVicente_04VV08_1
c90m4p6r2 (RAxML)
0.002

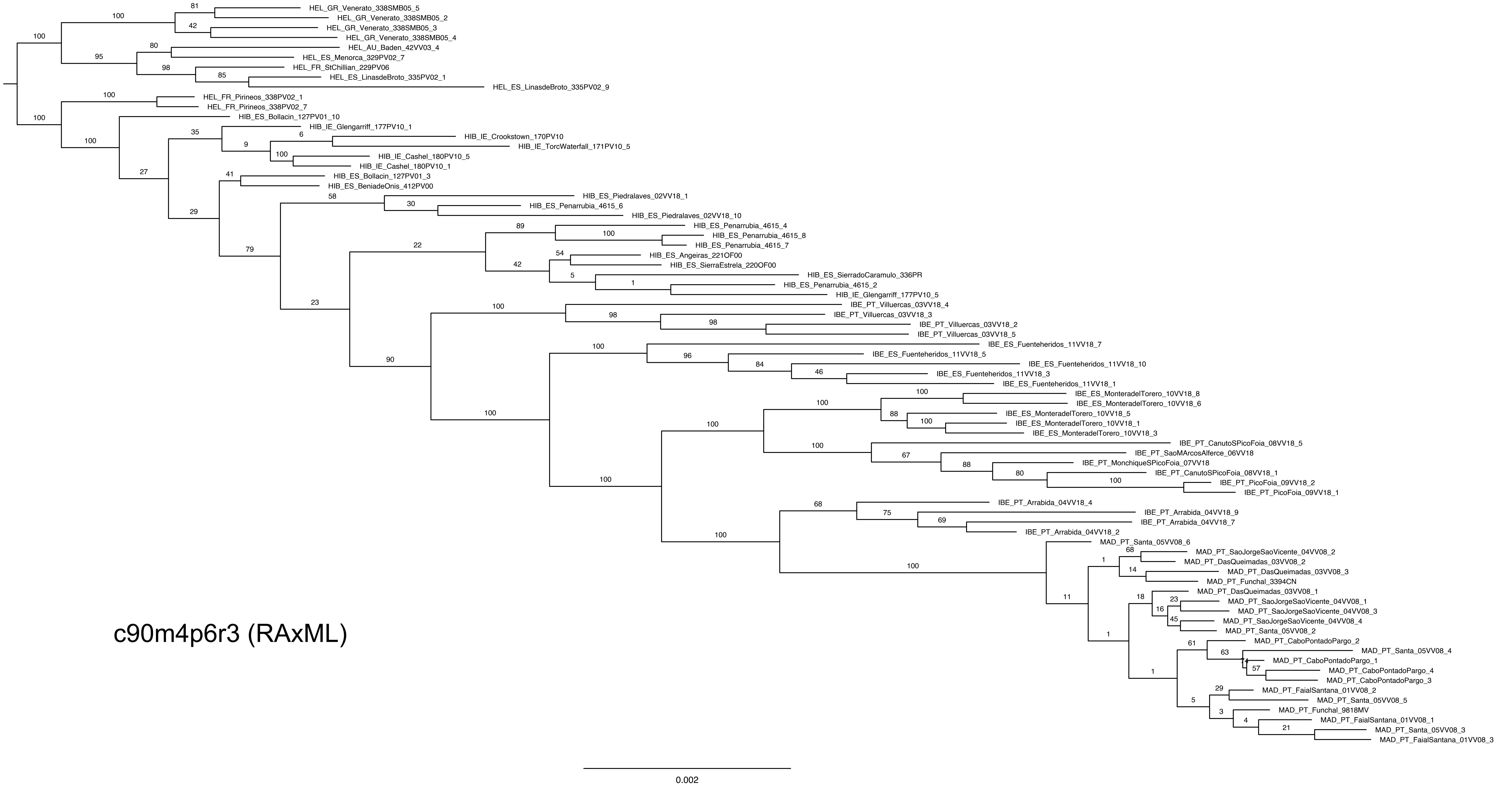
HEL_GR_Venerato_338SMB05_5
HEL_GR_Venerato_338SMB05_2
HEL_GR_Venerato_338SMB05_3
HEL_GR_Venerato_338SMB05_4
HEL_AU_Baden_42VV03_4
HEL_ES_Menorca_329PV02_7
HEL_FR_StChillian_229PV06
HEL_ES_LinasdeBroto_335PV02_1
HEL_ES_LinasdeBroto_335PV02_9
HEL_FR_Pirineos_338PV02_1
HEL_FR_Pirineos_338PV02_7
HIB_ES_Bollacin_127PV01_10
HIB_IE_Glengarriff_177PV10_1
HIB_IE_Crookstown_170PV10
HIB_IE_TorcWaterfall_171PV10_5
HIB_IE_Cashel_180PV10_5
HIB_IE_Cashel_180PV10_1
HIB_ES_Bollacin_127PV01_3
HIB_ES_BeniadeOnis_412PV00
HIB_ES_Piedralaves_02VV18_1
HIB_ES_Penarrubia_4615_6
HIB_ES_Piedralaves_02VV18_10
HIB_ES_Penarrubia_4615_4
HIB_ES_Penarrubia_4615_8
HIB_ES_Penarrubia_4615_7
HIB_ES_Angeiras_221OF00
HIB_ES_SierraEstrela_220OF00
HIB_ES_SierradoCaramulo_336PR
HIB_ES_Penarrubia_4615_2
HIB_IE_Glengarriff_177PV10_5
IBE_PT_Villuercas_03VV18_4
IBE_PT_Villuercas_03VV18_3
IBE_PT_Villuercas_03VV18_2
IBE_PT_Villuercas_03VV18_5
IBE_ES_Fuenteheridos_11VV18_7
IBE_ES_Fuenteheridos_11VV18_5
IBE_ES_Fuenteheridos_11VV18_10
IBE_ES_Fuenteheridos_11VV18_3
IBE_ES_Fuenteheridos_11VV18_1
IBE_ES_MonteradelTorero_10VV18_8
IBE_ES_MonteradelTorero_10VV18_6
IBE_ES_MonteradelTorero_10VV18_5
IBE_ES_MonteradelTorero_10VV18_1
IBE_ES_MonteradelTorero_10VV18_3
IBE_PT_CanutoSPicoFoia_08VV18_5
IBE_PT_SaoMArcosAlferce_06VV18
IBE_PT_MonchiqueSPicoFoia_07VV18
IBE_PT_CanutoSPicoFoia_08VV18_1
IBE_PT_PicoFoia_09VV18_2
IBE_PT_PicoFoia_09VV18_1
IBE_PT_Arrabida_04VV18_4
IBE_PT_Arrabida_04VV18_9
IBE_PT_Arrabida_04VV18_7
IBE_PT_Arrabida_04VV18_2
MAD_PT_Santa_05VV08_6
MAD_PT_SaoJorgeSaoVicente_04VV08_2
MAD_PT_DasQueimadas_03VV08_2
MAD_PT_DasQueimadas_03VV08_3
MAD_PT_Funchal_3394CN
MAD_PT_DasQueimadas_03VV08_1
MAD_PT_SaoJorgeSaoVicente_04VV08_1
MAD_PT_SaoJorgeSaoVicente_04VV08_3
MAD_PT_SaoJorgeSaoVicente_04VV08_4
MAD_PT_Santa_05VV08_2
MAD_PT_CaboPontadoPargo_2
MAD_PT_Santa_05VV08_4
MAD_PT_CaboPontadoPargo_1
MAD_PT_CaboPontadoPargo_4
MAD_PT_CaboPontadoPargo_3
MAD_PT_FaialSantana_01VV08_2
MAD_PT_Santa_05VV08_5
MAD_PT_Funchal_9818MV
MAD_PT_FaialSantana_01VV08_1
MAD_PT_Santa_05VV08_3
MAD_PT_FaialSantana_01VV08_3
c90m4p6r3 (RAxML)
0.002

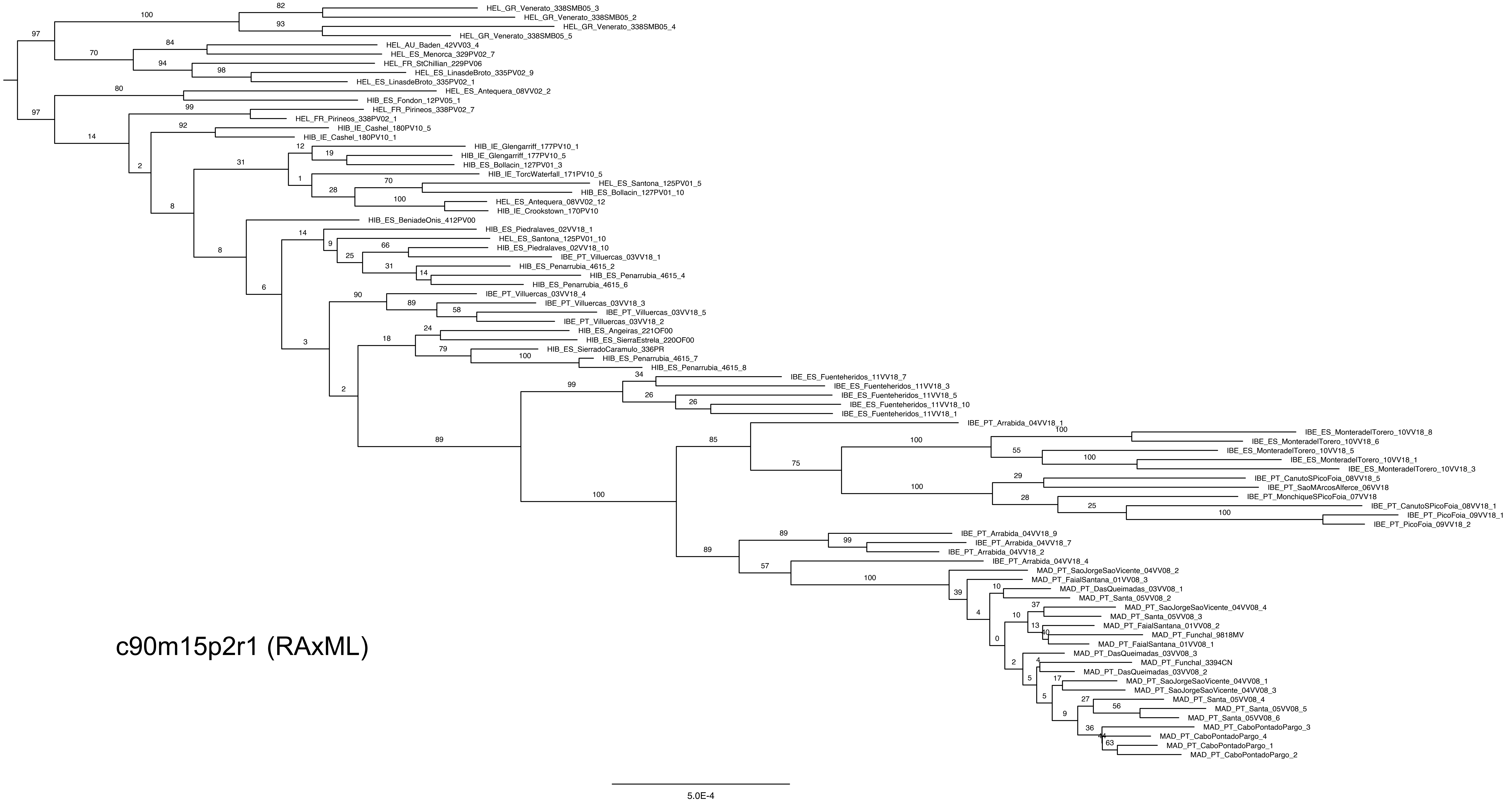
HEL_GR_Venerato_338SMB05_3
HEL_GR_Venerato_338SMB05_2
HEL_GR_Venerato_338SMB05_4
HEL_GR_Venerato_338SMB05_5
HEL_AU_Baden_42VV03_4
HEL_ES_Menorca_329PV02_7
HEL_FR_StChillian_229PV06
HEL_ES_LinasdeBroto_335PV02_9
HEL_ES_LinasdeBroto_335PV02_1
HEL_ES_Antequera_08VV02_2
HIB_ES_Fondon_12PV05_1
HEL_FR_Pirineos_338PV02_7
HEL_FR_Pirineos_338PV02_1
HIB_IE_Cashel_180PV10_5
HIB_IE_Cashel_180PV10_1
HIB_IE_Glengarriff_177PV10_1
HIB_IE_Glengarriff_177PV10_5
HIB_ES_Bollacin_127PV01_3
HIB_IE_TorcWaterfall_171PV10_5
HEL_ES_Santona_125PV01_5
HIB_ES_Bollacin_127PV01_10
HEL_ES_Antequera_08VV02_12
HIB_IE_Crookstown_170PV10
HIB_ES_BeniadeOnis_412PV00
HIB_ES_Piedralaves_02VV18_1
HEL_ES_Santona_125PV01_10
HIB_ES_Piedralaves_02VV18_10
IBE_PT_Villuercas_03VV18_1
HIB_ES_Penarrubia_4615_2
HIB_ES_Penarrubia_4615_4
HIB_ES_Penarrubia_4615_6
IBE_PT_Villuercas_03VV18_4
IBE_PT_Villuercas_03VV18_3
IBE_PT_Villuercas_03VV18_5
IBE_PT_Villuercas_03VV18_2
HIB_ES_Angeiras_221OF00
HIB_ES_SierraEstrela_220OF00
HIB_ES_SierradoCaramulo_336PR
HIB_ES_Penarrubia_4615_7
HIB_ES_Penarrubia_4615_8
IBE_ES_Fuenteheridos_11VV18_7
IBE_ES_Fuenteheridos_11VV18_3
IBE_ES_Fuenteheridos_11VV18_5
IBE_ES_Fuenteheridos_11VV18_10
IBE_ES_Fuenteheridos_11VV18_1
IBE_PT_Arrabida_04VV18_1
IBE_ES_MonteradelTorero_10VV18_8
IBE_ES_MonteradelTorero_10VV18_6
IBE_ES_MonteradelTorero_10VV18_5
IBE_ES_MonteradelTorero_10VV18_1
IBE_ES_MonteradelTorero_10VV18_3
IBE_PT_CanutoSPicoFoia_08VV18_5
IBE_PT_SaoMArcosAlferce_06VV18
IBE_PT_MonchiqueSPicoFoia_07VV18
IBE_PT_CanutoSPicoFoia_08VV18_1
IBE_PT_PicoFoia_09VV18_1
IBE_PT_PicoFoia_09VV18_2
IBE_PT_Arrabida_04VV18_9
IBE_PT_Arrabida_04VV18_7
IBE_PT_Arrabida_04VV18_2
IBE_PT_Arrabida_04VV18_4
MAD_PT_SaoJorgeSaoVicente_04VV08_2
MAD_PT_FaialSantana_01VV08_3
MAD_PT_DasQueimadas_03VV08_1
MAD_PT_Santa_05VV08_2
MAD_PT_SaoJorgeSaoVicente_04VV08_4
MAD_PT_Santa_05VV08_3
MAD_PT_FaialSantana_01VV08_2
MAD_PT_Funchal_9818MV
MAD_PT_FaialSantana_01VV08_1
MAD_PT_DasQueimadas_03VV08_3
MAD_PT_Funchal_3394CN
MAD_PT_DasQueimadas_03VV08_2
MAD_PT_SaoJorgeSaoVicente_04VV08_1
MAD_PT_SaoJorgeSaoVicente_04VV08_3
MAD_PT_Santa_05VV08_4
MAD_PT_Santa_05VV08_5
MAD_PT_Santa_05VV08_6
MAD_PT_CaboPontadoPargo_3
MAD_PT_CaboPontadoPargo_4
MAD_PT_CaboPontadoPargo_1
MAD_PT_CaboPontadoPargo_2
c90m15p2r1 (RAxML)
5.0E-4

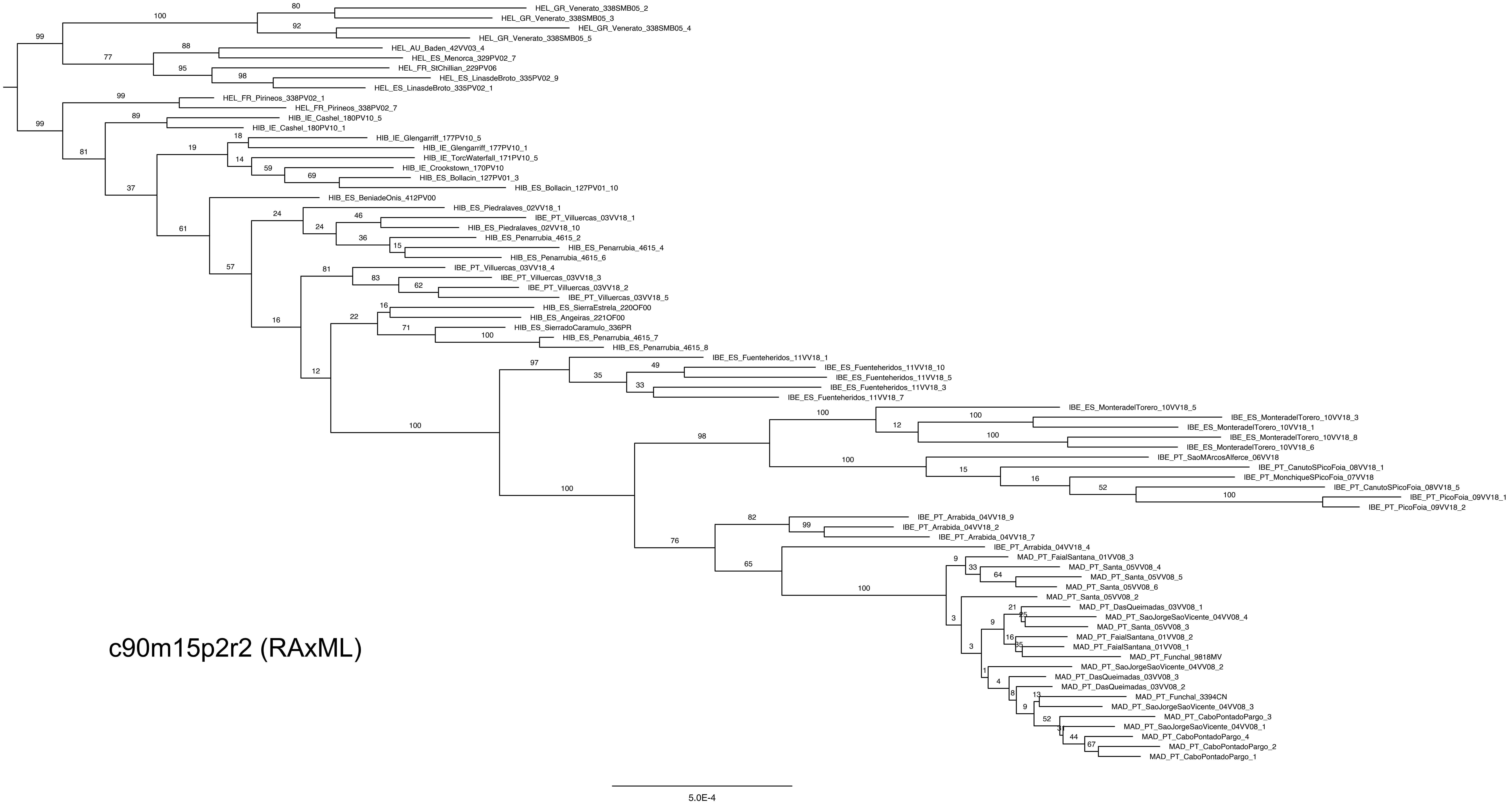
HEL_GR_Venerato_338SMB05_2
HEL_GR_Venerato_338SMB05_3
HEL_GR_Venerato_338SMB05_4
HEL_GR_Venerato_338SMB05_5
HEL_AU_Baden_42VV03_4
HEL_ES_Menorca_329PV02_7
HEL_FR_StChillian_229PV06
HEL_ES_LinasdeBroto_335PV02_9
HEL_ES_LinasdeBroto_335PV02_1
HEL_FR_Pirineos_338PV02_1
HEL_FR_Pirineos_338PV02_7
HIB_IE_Cashel_180PV10_5
HIB_IE_Cashel_180PV10_1
HIB_IE_Glengarriff_177PV10_5
HIB_IE_Glengarriff_177PV10_1
HIB_IE_TorcWaterfall_171PV10_5
HIB_IE_Crookstown_170PV10
HIB_ES_Bollacin_127PV01_3
HIB_ES_Bollacin_127PV01_10
HIB_ES_BeniadeOnis_412PV00
HIB_ES_Piedralaves_02VV18_1
IBE_PT_Villuercas_03VV18_1
HIB_ES_Piedralaves_02VV18_10
HIB_ES_Penarrubia_4615_2
HIB_ES_Penarrubia_4615_4
HIB_ES_Penarrubia_4615_6
IBE_PT_Villuercas_03VV18_4
IBE_PT_Villuercas_03VV18_3
IBE_PT_Villuercas_03VV18_2
IBE_PT_Villuercas_03VV18_5
HIB_ES_SierraEstrela_220OF00
HIB_ES_Angeiras_221OF00
HIB_ES_SierradoCaramulo_336PR
HIB_ES_Penarrubia_4615_7
HIB_ES_Penarrubia_4615_8
IBE_ES_Fuenteheridos_11VV18_1
IBE_ES_Fuenteheridos_11VV18_10
IBE_ES_Fuenteheridos_11VV18_5
IBE_ES_Fuenteheridos_11VV18_3
IBE_ES_Fuenteheridos_11VV18_7
IBE_ES_MonteradelTorero_10VV18_5
IBE_ES_MonteradelTorero_10VV18_3
IBE_ES_MonteradelTorero_10VV18_1
IBE_ES_MonteradelTorero_10VV18_8
IBE_ES_MonteradelTorero_10VV18_6
IBE_PT_SaoMArcosAlferce_06VV18
IBE_PT_CanutoSPicoFoia_08VV18_1
IBE_PT_MonchiqueSPicoFoia_07VV18
IBE_PT_CanutoSPicoFoia_08VV18_5
IBE_PT_PicoFoia_09VV18_1
IBE_PT_PicoFoia_09VV18_2
IBE_PT_Arrabida_04VV18_9
IBE_PT_Arrabida_04VV18_2
IBE_PT_Arrabida_04VV18_7
IBE_PT_Arrabida_04VV18_4
MAD_PT_FaialSantana_01VV08_3
MAD_PT_Santa_05VV08_4
MAD_PT_Santa_05VV08_5
MAD_PT_Santa_05VV08_6
MAD_PT_Santa_05VV08_2
MAD_PT_DasQueimadas_03VV08_1
MAD_PT_SaoJorgeSaoVicente_04VV08_4
MAD_PT_Santa_05VV08_3
MAD_PT_FaialSantana_01VV08_2
MAD_PT_FaialSantana_01VV08_1
MAD_PT_Funchal_9818MV
MAD_PT_SaoJorgeSaoVicente_04VV08_2
MAD_PT_DasQueimadas_03VV08_3
MAD_PT_DasQueimadas_03VV08_2
MAD_PT_Funchal_3394CN
MAD_PT_SaoJorgeSaoVicente_04VV08_3
MAD_PT_CaboPontadoPargo_3
MAD_PT_SaoJorgeSaoVicente_04VV08_1
MAD_PT_CaboPontadoPargo_4
MAD_PT_CaboPontadoPargo_2
MAD_PT_CaboPontadoPargo_1
c90m15p2r2 (RAxML)
5.0E-4

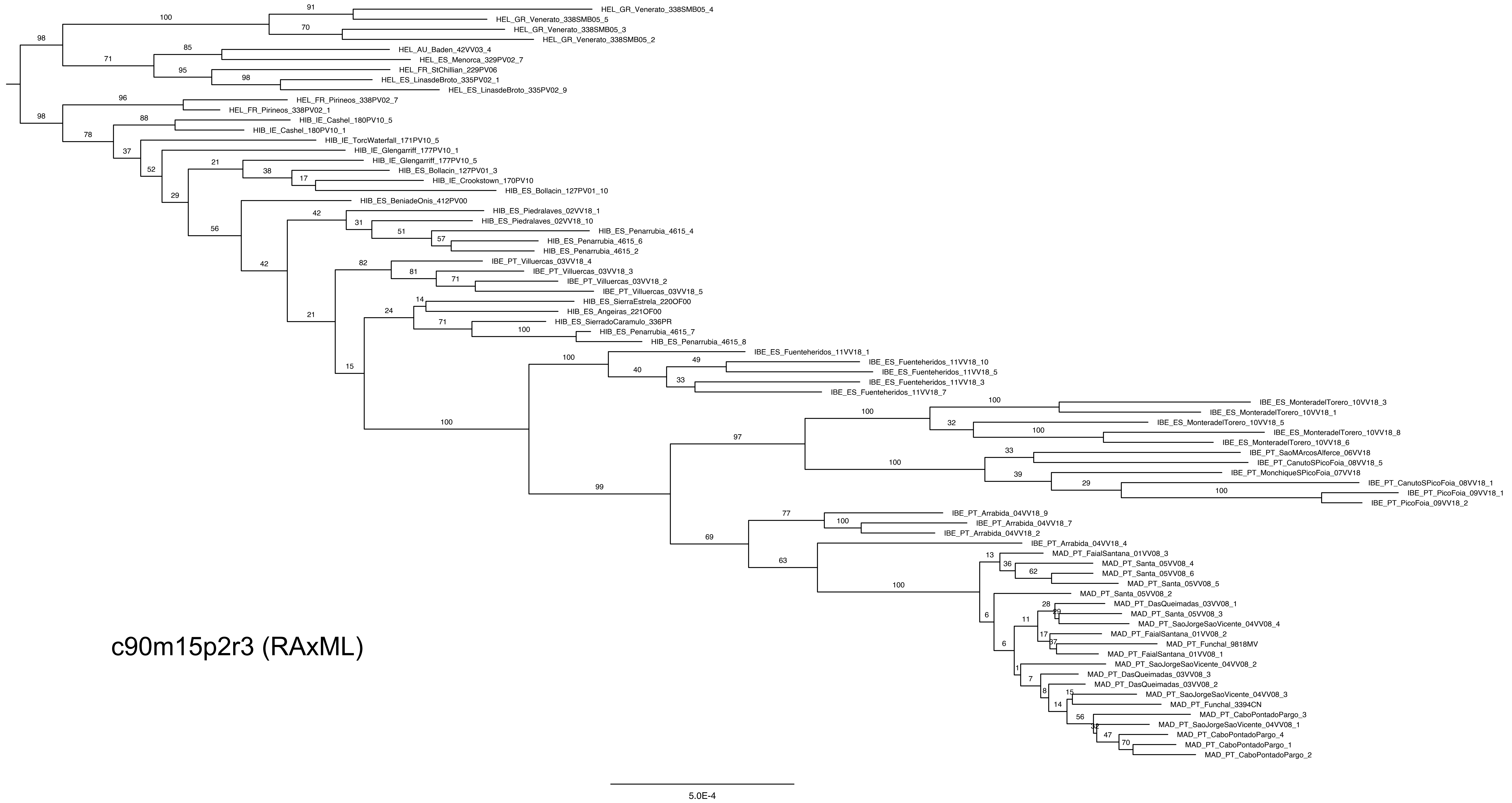

c90m15p2r3 (RAxML)
HEL_GR_Venerato_338SMB05_4
HEL_GR_Venerato_338SMB05_5
HEL_GR_Venerato_338SMB05_3
HEL_GR_Venerato_338SMB05_2
HEL_AU_Baden_42VV03_4
HEL_ES_Menorca_329PV02_7
HEL_FR_StChillian_229PV06
HEL_ES_LinasdeBroto_335PV02_1
HEL_ES_LinasdeBroto_335PV02_9
HEL_FR_Pirineos_338PV02_7
HEL_FR_Pirineos_338PV02_1
HIB_IE_Cashel_180PV10_5
HIB_IE_Cashel_180PV10_1
HIB_IE_TorcWaterfall_171PV10_5
HIB_IE_Glengarriff_177PV10_1
HIB_IE_Glengarriff_177PV10_5
HIB_ES_Bollacin_127PV01_3
HIB_IE_Crookstown_170PV10
HIB_ES_Bollacin_127PV01_10
HIB_ES_BeniadeOnis_412PV00
HIB_ES_Piedralaves_02VV18_1
HIB_ES_Piedralaves_02VV18_10
HIB_ES_Penarrubia_4615_4
HIB_ES_Penarrubia_4615_6
HIB_ES_Penarrubia_4615_2
IBE_PT_Villuercas_03VV18_4
IBE_PT_Villuercas_03VV18_3
IBE_PT_Villuercas_03VV18_2
IBE_PT_Villuercas_03VV18_5
HIB_ES_SierraEstrela_220OF00
HIB_ES_Angeiras_221OF00
HIB_ES_SierradoCaramulo_336PR
HIB_ES_Penarrubia_4615_7
HIB_ES_Penarrubia_4615_8
IBE_ES_Fuenteheridos_11VV18_1
IBE_ES_Fuenteheridos_11VV18_10
IBE_ES_Fuenteheridos_11VV18_5
IBE_ES_Fuenteheridos_11VV18_3
IBE_ES_Fuenteheridos_11VV18_7
IBE_ES_MonteradelTorero_10VV18_3
IBE_ES_MonteradelTorero_10VV18_1
IBE_ES_MonteradelTorero_10VV18_5
IBE_ES_MonteradelTorero_10VV18_8
IBE_ES_MonteradelTorero_10VV18_6
IBE_PT_SaoMArcosAlferce_06VV18
IBE_PT_CanutoSPicoFoia_08VV18_5
IBE_PT_MonchiqueSPicoFoia_07VV18
IBE_PT_CanutoSPicoFoia_08VV18_1
IBE_PT_PicoFoia_09VV18_1
IBE_PT_PicoFoia_09VV18_2
IBE_PT_Arrabida_04VV18_9
IBE_PT_Arrabida_04VV18_7
IBE_PT_Arrabida_04VV18_2
IBE_PT_Arrabida_04VV18_4
MAD_PT_FaialSantana_01VV08_3
MAD_PT_Santa_05VV08_4
MAD_PT_Santa_05VV08_6
MAD_PT_Santa_05VV08_5
MAD_PT_Santa_05VV08_2
MAD_PT_DasQueimadas_03VV08_1
MAD_PT_Santa_05VV08_3
MAD_PT_SaoJorgeSaoVicente_04VV08_4
MAD_PT_FaialSantana_01VV08_2
MAD_PT_Funchal_9818MV
MAD_PT_FaialSantana_01VV08_1
MAD_PT_SaoJorgeSaoVicente_04VV08_2
MAD_PT_DasQueimadas_03VV08_3
MAD_PT_DasQueimadas_03VV08_2
MAD_PT_SaoJorgeSaoVicente_04VV08_3
MAD_PT_Funchal_3394CN
MAD_PT_CaboPontadoPargo_3
MAD_PT_SaoJorgeSaoVicente_04VV08_1
MAD_PT_CaboPontadoPargo_4
MAD_PT_CaboPontadoPargo_1
MAD_PT_CaboPontadoPargo_2
5.0E-4

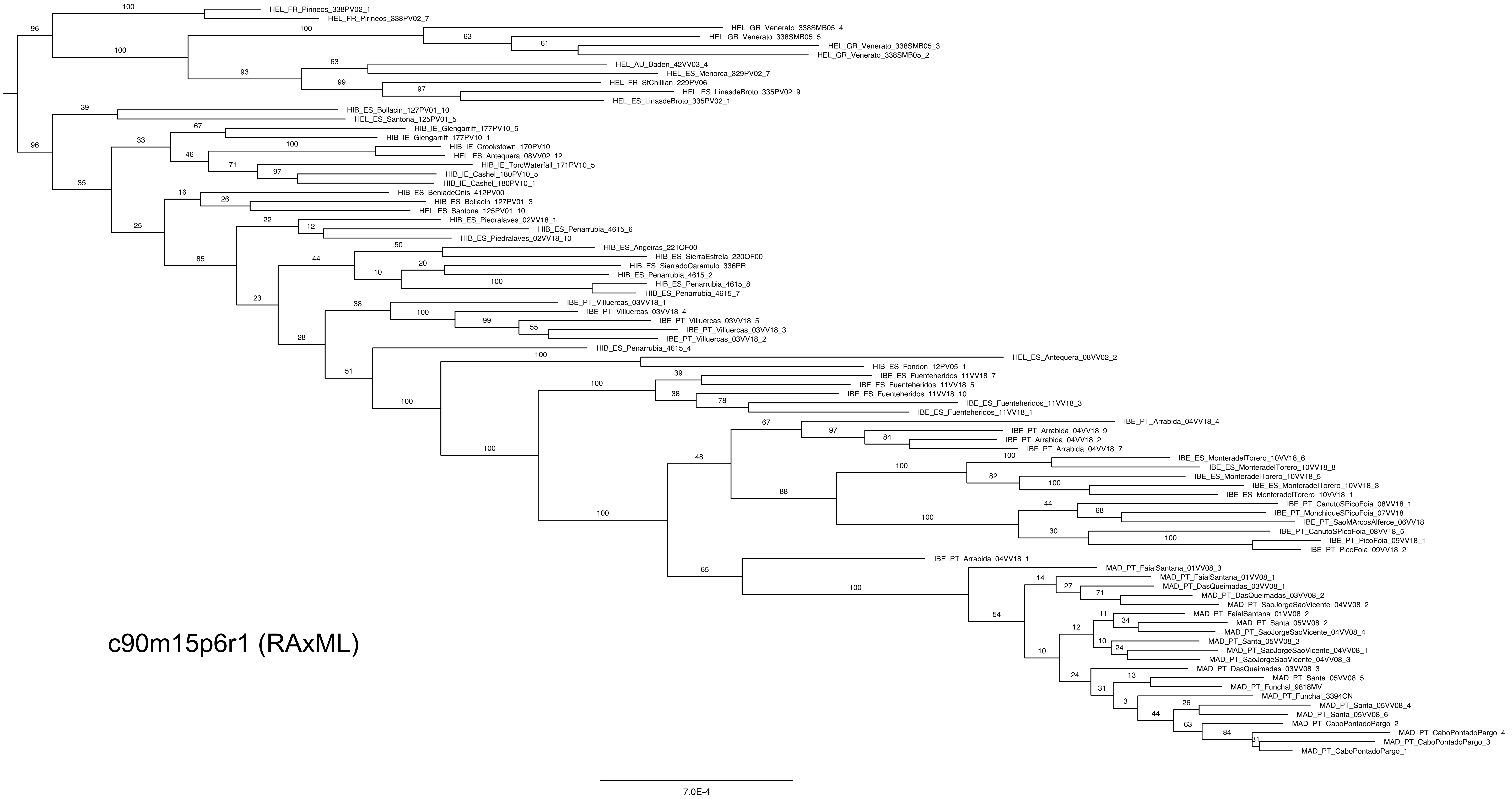

HEL_FR_Pirineos_338PV02_1
HEL_FR_Pirineos_338PV02_7
HEL_GR_Venerato_338SMB05_4
HEL_GR_Venerato_338SMB05_5
HEL_GR_Venerato_338SMB05_3
HEL_GR_Venerato_338SMB05_2
HEL_AU_Baden_42VV03_4
HEL_ES_Menorca_329PV02_7
HEL_FR_StChillian_229PV06
HEL_ES_LinasdeBroto_335PV02_9
HEL_ES_LinasdeBroto_335PV02_1
HIB_ES_Bollacin_127PV01_10
HEL_ES_Santona_125PV01_5
HIB_IE_Glengarriff_177PV10_5
HIB_IE_Glengarriff_177PV10_1
HIB_IE_Crookstown_170PV10
HEL_ES_Antequera_08VV02_12
HIB_IE_TorcWaterfall_171PV10_5
HIB_IE_Cashel_180PV10_5
HIB_IE_Cashel_180PV10_1
HIB_ES_BeniadeOnis_412PV00
HIB_ES_Bollacin_127PV01_3
HEL_ES_Santona_125PV01_10
HIB_ES_Piedralaves_02VV18_1
HIB_ES_Penarrubia_4615_6
HIB_ES_Piedralaves_02VV18_10
HIB_ES_Angeiras_221OF00
HIB_ES_SierraEstrela_220OF00
HIB_ES_SierradoCaramulo_336PR
HIB_ES_Penarrubia_4615_2
HIB_ES_Penarrubia_4615_8
HIB_ES_Penarrubia_4615_7
IBE_PT_Villuercas_03VV18_1
IBE_PT_Villuercas_03VV18_4
IBE_PT_Villuercas_03VV18_5
IBE_PT_Villuercas_03VV18_3
IBE_PT_Villuercas_03VV18_2
HIB_ES_Penarrubia_4615_4
HEL_ES_Antequera_08VV02_2
HIB_ES_Fondon_12PV05_1
IBE_ES_Fuenteheridos_11VV18_7
IBE_ES_Fuenteheridos_11VV18_5
IBE_ES_Fuenteheridos_11VV18_10
IBE_ES_Fuenteheridos_11VV18_3
IBE_ES_Fuenteheridos_11VV18_1
IBE_PT_Arrabida_04VV18_4
IBE_PT_Arrabida_04VV18_9
IBE_PT_Arrabida_04VV18_2
IBE_PT_Arrabida_04VV18_7
IBE_ES_MonteradelTorero_10VV18_6
IBE_ES_MonteradelTorero_10VV18_8
IBE_ES_MonteradelTorero_10VV18_5
IBE_ES_MonteradelTorero_10VV18_3
IBE_ES_MonteradelTorero_10VV18_1
IBE_PT_CanutoSPicoFoia_08VV18_1
IBE_PT_MonchiqueSPicoFoia_07VV18
IBE_PT_SaoMArcosAlferce_06VV18
IBE_PT_CanutoSPicoFoia_08VV18_5
IBE_PT_PicoFoia_09VV18_1
IBE_PT_PicoFoia_09VV18_2
IBE_PT_Arrabida_04VV18_1
MAD_PT_FaialSantana_01VV08_3
MAD_PT_FaialSantana_01VV08_1
MAD_PT_DasQueimadas_03VV08_1
MAD_PT_DasQueimadas_03VV08_2
MAD_PT_SaoJorgeSaoVicente_04VV08_2
MAD_PT_FaialSantana_01VV08_2
MAD_PT_Santa_05VV08_2
MAD_PT_SaoJorgeSaoVicente_04VV08_4
MAD_PT_Santa_05VV08_3
MAD_PT_SaoJorgeSaoVicente_04VV08_1
MAD_PT_SaoJorgeSaoVicente_04VV08_3
MAD_PT_DasQueimadas_03VV08_3
MAD_PT_Santa_05VV08_5
MAD_PT_Funchal_9818MV
MAD_PT_Funchal_3394CN
MAD_PT_Santa_05VV08_4
MAD_PT_Santa_05VV08_6
MAD_PT_CaboPontadoPargo_2
MAD_PT_CaboPontadoPargo_4
MAD_PT_CaboPontadoPargo_3
MAD_PT_CaboPontadoPargo_1
c90m15p6r1 (RAxML)
7.0E-4

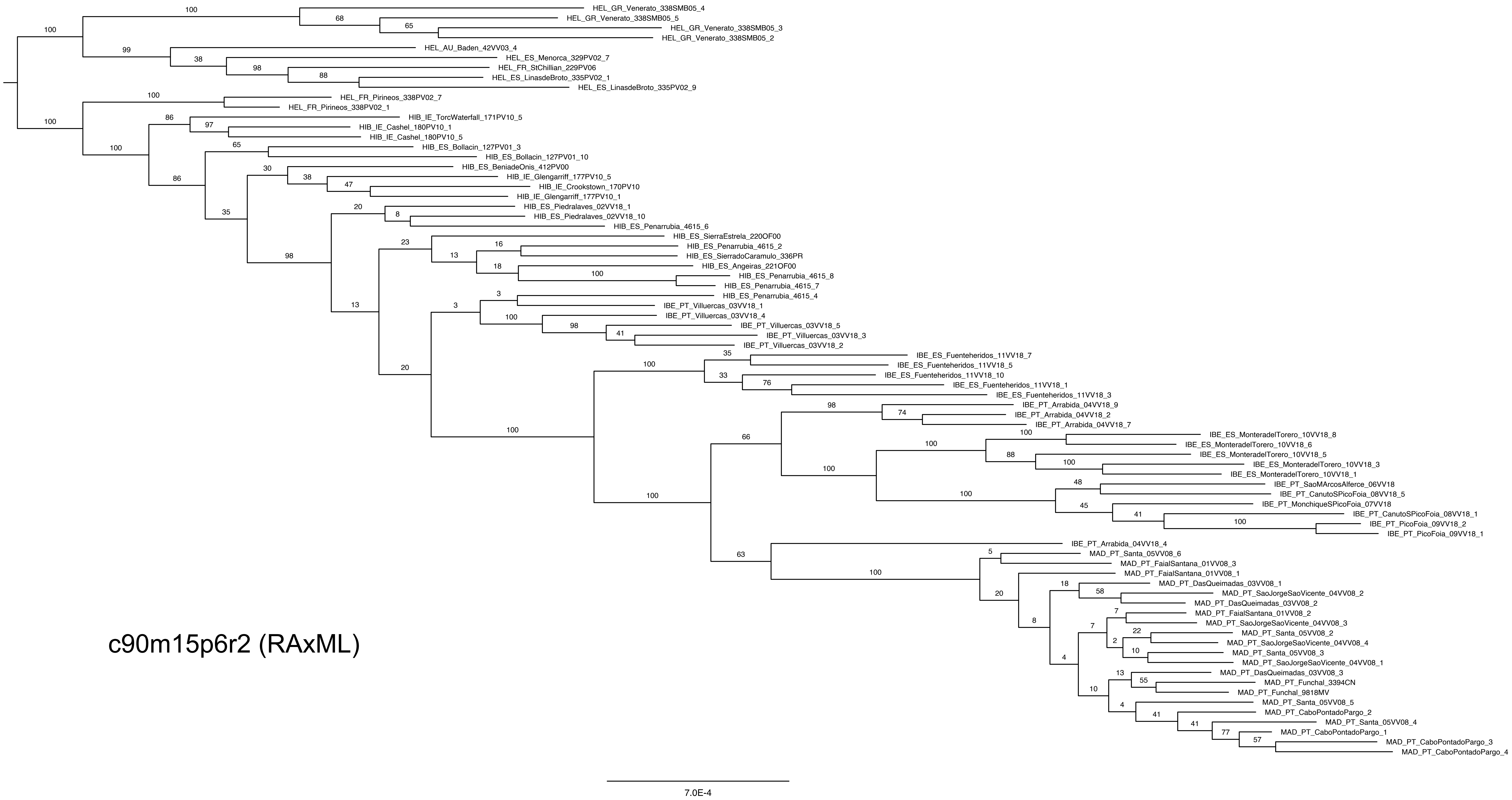
HEL_GR_Venerato_338SMB05_4
HEL_GR_Venerato_338SMB05_5
HEL_GR_Venerato_338SMB05_3
HEL_GR_Venerato_338SMB05_2
HEL_AU_Baden_42VV03_4
HEL_ES_Menorca_329PV02_7
HEL_FR_StChillian_229PV06
HEL_ES_LinasdeBroto_335PV02_1
HEL_ES_LinasdeBroto_335PV02_9
HEL_FR_Pirineos_338PV02_7
HEL_FR_Pirineos_338PV02_1
HIB_IE_TorcWaterfall_171PV10_5
HIB_IE_Cashel_180PV10_1
HIB_IE_Cashel_180PV10_5
HIB_ES_Bollacin_127PV01_3
HIB_ES_Bollacin_127PV01_10
HIB_ES_BeniadeOnis_412PV00
HIB_IE_Glengarriff_177PV10_5
HIB_IE_Crookstown_170PV10
HIB_IE_Glengarriff_177PV10_1
HIB_ES_Piedralaves_02VV18_1
HIB_ES_Piedralaves_02VV18_10
HIB_ES_Penarrubia_4615_6
HIB_ES_SierraEstrela_220OF00
HIB_ES_Penarrubia_4615_2
HIB_ES_SierradoCaramulo_336PR
HIB_ES_Angeiras_221OF00
HIB_ES_Penarrubia_4615_8
HIB_ES_Penarrubia_4615_7
HIB_ES_Penarrubia_4615_4
IBE_PT_Villuercas_03VV18_1
IBE_PT_Villuercas_03VV18_4
IBE_PT_Villuercas_03VV18_5
IBE_PT_Villuercas_03VV18_3
IBE_PT_Villuercas_03VV18_2
IBE_ES_Fuenteheridos_11VV18_7
IBE_ES_Fuenteheridos_11VV18_5
IBE_ES_Fuenteheridos_11VV18_10
IBE_ES_Fuenteheridos_11VV18_1
IBE_ES_Fuenteheridos_11VV18_3
IBE_PT_Arrabida_04VV18_9
IBE_PT_Arrabida_04VV18_2
IBE_PT_Arrabida_04VV18_7
IBE_ES_MonteradelTorero_10VV18_8
IBE_ES_MonteradelTorero_10VV18_6
IBE_ES_MonteradelTorero_10VV18_5
IBE_ES_MonteradelTorero_10VV18_3
IBE_ES_MonteradelTorero_10VV18_1
IBE_PT_SaoMArcosAlferce_06VV18
IBE_PT_CanutoSPicoFoia_08VV18_5
IBE_PT_MonchiqueSPicoFoia_07VV18
IBE_PT_CanutoSPicoFoia_08VV18_1
IBE_PT_PicoFoia_09VV18_2
IBE_PT_PicoFoia_09VV18_1
IBE_PT_Arrabida_04VV18_4
MAD_PT_Santa_05VV08_6
MAD_PT_FaialSantana_01VV08_3
MAD_PT_FaialSantana_01VV08_1
MAD_PT_DasQueimadas_03VV08_1
MAD_PT_SaoJorgeSaoVicente_04VV08_2
MAD_PT_DasQueimadas_03VV08_2
MAD_PT_FaialSantana_01VV08_2
MAD_PT_SaoJorgeSaoVicente_04VV08_3
MAD_PT_Santa_05VV08_2
MAD_PT_SaoJorgeSaoVicente_04VV08_4
MAD_PT_Santa_05VV08_3
MAD_PT_SaoJorgeSaoVicente_04VV08_1
MAD_PT_DasQueimadas_03VV08_3
MAD_PT_Funchal_3394CN
MAD_PT_Funchal_9818MV
MAD_PT_Santa_05VV08_5
MAD_PT_CaboPontadoPargo_2
MAD_PT_Santa_05VV08_4
MAD_PT_CaboPontadoPargo_1
MAD_PT_CaboPontadoPargo_3
MAD_PT_CaboPontadoPargo_4
c90m15p6r2 (RAxML)
7.0E-4

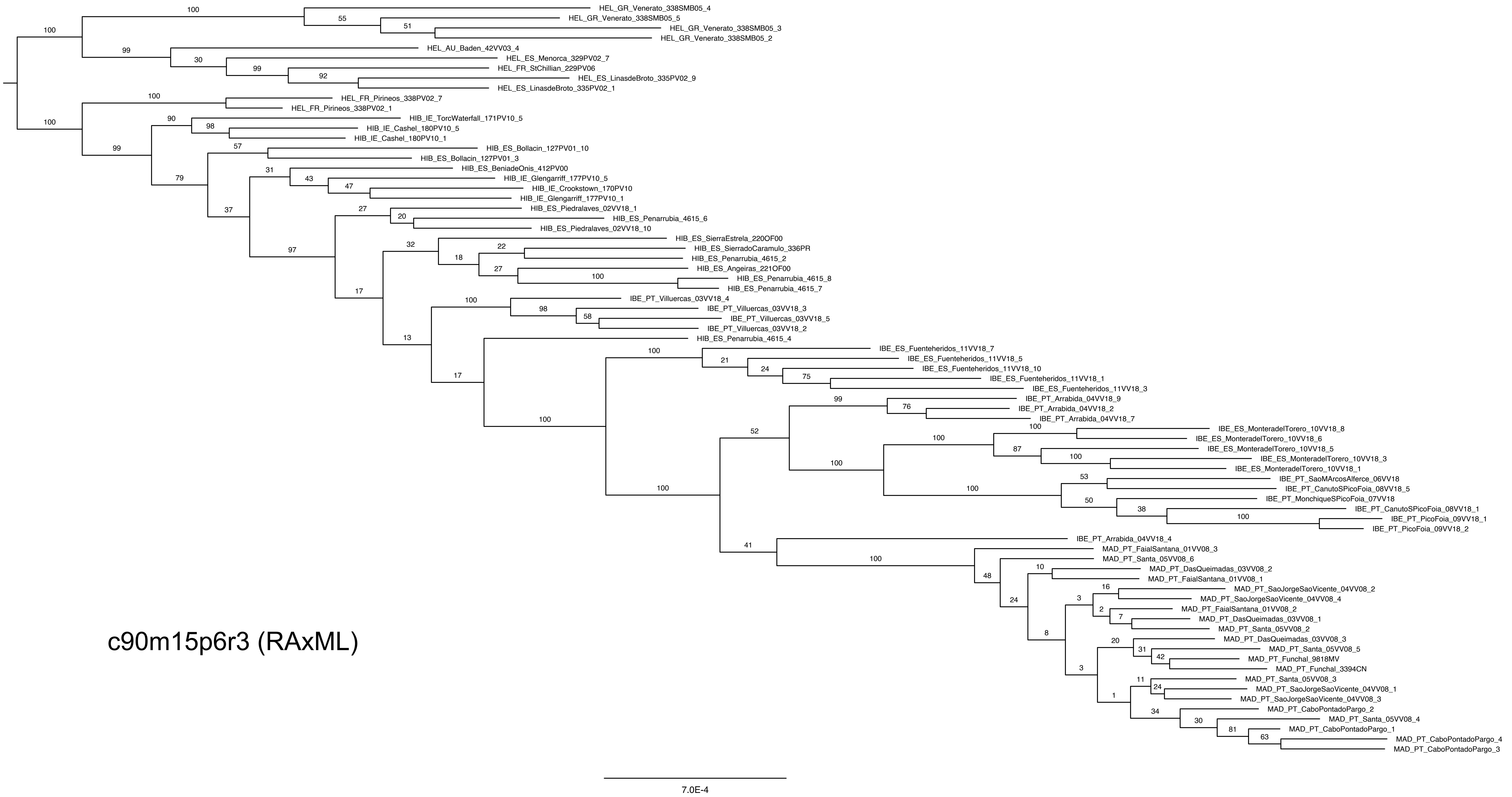


c90m15p6r3 (RAxML)

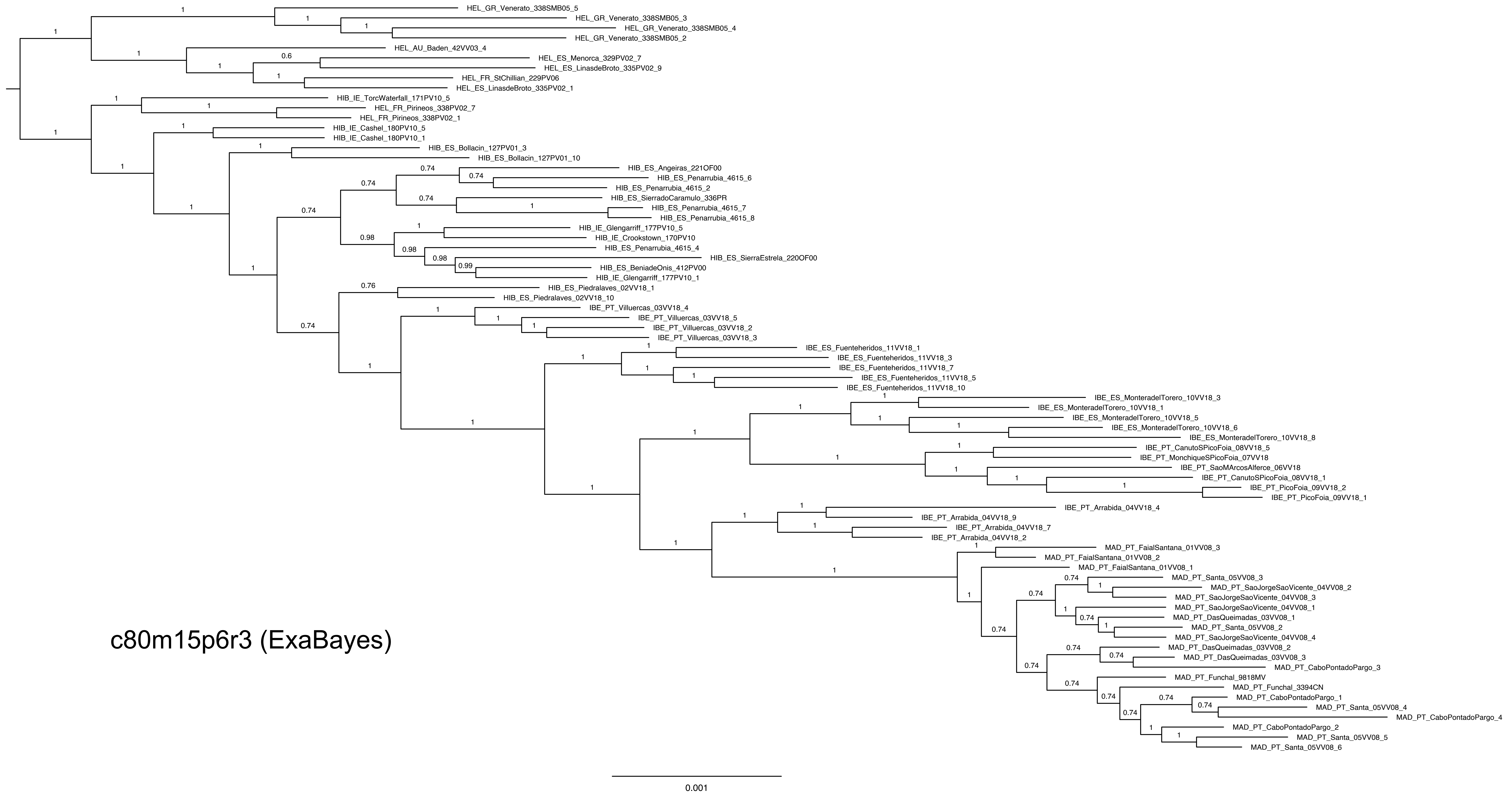
HEL_GR_Venerato_338SMB05_5
HEL_GR_Venerato_338SMB05_3
HEL_GR_Venerato_338SMB05_4
HEL_GR_Venerato_338SMB05_2
HEL_AU_Baden_42VV03_4
HEL_ES_Menorca_329PV02_7
HEL_ES_LinasdeBroto_335PV02_9
HEL_FR_StChillian_229PV06
HEL_ES_LinasdeBroto_335PV02_1
HIB_IE_TorcWaterfall_171PV10_5
HEL_FR_Pirineos_338PV02_7
HEL_FR_Pirineos_338PV02_1
HIB_IE_Cashel_180PV10_5
HIB_IE_Cashel_180PV10_1
HIB_ES_Bollacin_127PV01_3
HIB_ES_Bollacin_127PV01_10
HIB_ES_Angeiras_221OF00
HIB_ES_Penarrubia_4615_6
HIB_ES_Penarrubia_4615_2
HIB_ES_SierradoCaramulo_336PR
HIB_ES_Penarrubia_4615_7
HIB_ES_Penarrubia_4615_8
HIB_IE_Glengarriff_177PV10_5
HIB_IE_Crookstown_170PV10
HIB_ES_Penarrubia_4615_4
HIB_ES_SierraEstrela_220OF00
HIB_ES_BeniadeOnis_412PV00
HIB_IE_Glengarriff_177PV10_1
HIB_ES_Piedralaves_02VV18_1
HIB_ES_Piedralaves_02VV18_10
IBE_PT_Villuercas_03VV18_4
IBE_PT_Villuercas_03VV18_5
IBE_PT_Villuercas_03VV18_2
IBE_PT_Villuercas_03VV18_3
IBE_ES_Fuenteheridos_11VV18_1
IBE_ES_Fuenteheridos_11VV18_3
IBE_ES_Fuenteheridos_11VV18_7
IBE_ES_Fuenteheridos_11VV18_5
IBE_ES_Fuenteheridos_11VV18_10
IBE_ES_MonteradelTorero_10VV18_3
IBE_ES_MonteradelTorero_10VV18_1
IBE_ES_MonteradelTorero_10VV18_5
IBE_ES_MonteradelTorero_10VV18_6
IBE_ES_MonteradelTorero_10VV18_8
IBE_PT_CanutoSPicoFoia_08VV18_5
IBE_PT_MonchiqueSPicoFoia_07VV18
IBE_PT_SaoMArcosAlferce_06VV18
IBE_PT_CanutoSPicoFoia_08VV18_1
IBE_PT_PicoFoia_09VV18_2
IBE_PT_PicoFoia_09VV18_1
IBE_PT_Arrabida_04VV18_4
IBE_PT_Arrabida_04VV18_9
IBE_PT_Arrabida_04VV18_7
IBE_PT_Arrabida_04VV18_2
MAD_PT_FaialSantana_01VV08_3
MAD_PT_FaialSantana_01VV08_2
MAD_PT_FaialSantana_01VV08_1
MAD_PT_Santa_05VV08_3
MAD_PT_SaoJorgeSaoVicente_04VV08_2
MAD_PT_SaoJorgeSaoVicente_04VV08_3
MAD_PT_SaoJorgeSaoVicente_04VV08_1
MAD_PT_DasQueimadas_03VV08_1
MAD_PT_Santa_05VV08_2
MAD_PT_SaoJorgeSaoVicente_04VV08_4
MAD_PT_DasQueimadas_03VV08_2
MAD_PT_DasQueimadas_03VV08_3
MAD_PT_CaboPontadoPargo_3
MAD_PT_Funchal_9818MV
MAD_PT_Funchal_3394CN
MAD_PT_CaboPontadoPargo_1
MAD_PT_Santa_05VV08_4
MAD_PT_CaboPontadoPargo_4
MAD_PT_CaboPontadoPargo_2
MAD_PT_Santa_05VV08_5
MAD_PT_Santa_05VV08_6
c80m15p6r3 (ExaBayes)
0.001

Supplementary figure 3. Results from population genetic structure analyses. (A) Three-dimensional representation of the genetic PCA (72% of variance accounted) from two different points of view. Envelopes encompass 95% of the observations for each species. (B) Genetic clustering of individuals using two different parameters in BAPS (K= 3, K= 4 ). Colours represent clusters. Each column represents an individual and its contribution to each cluster.

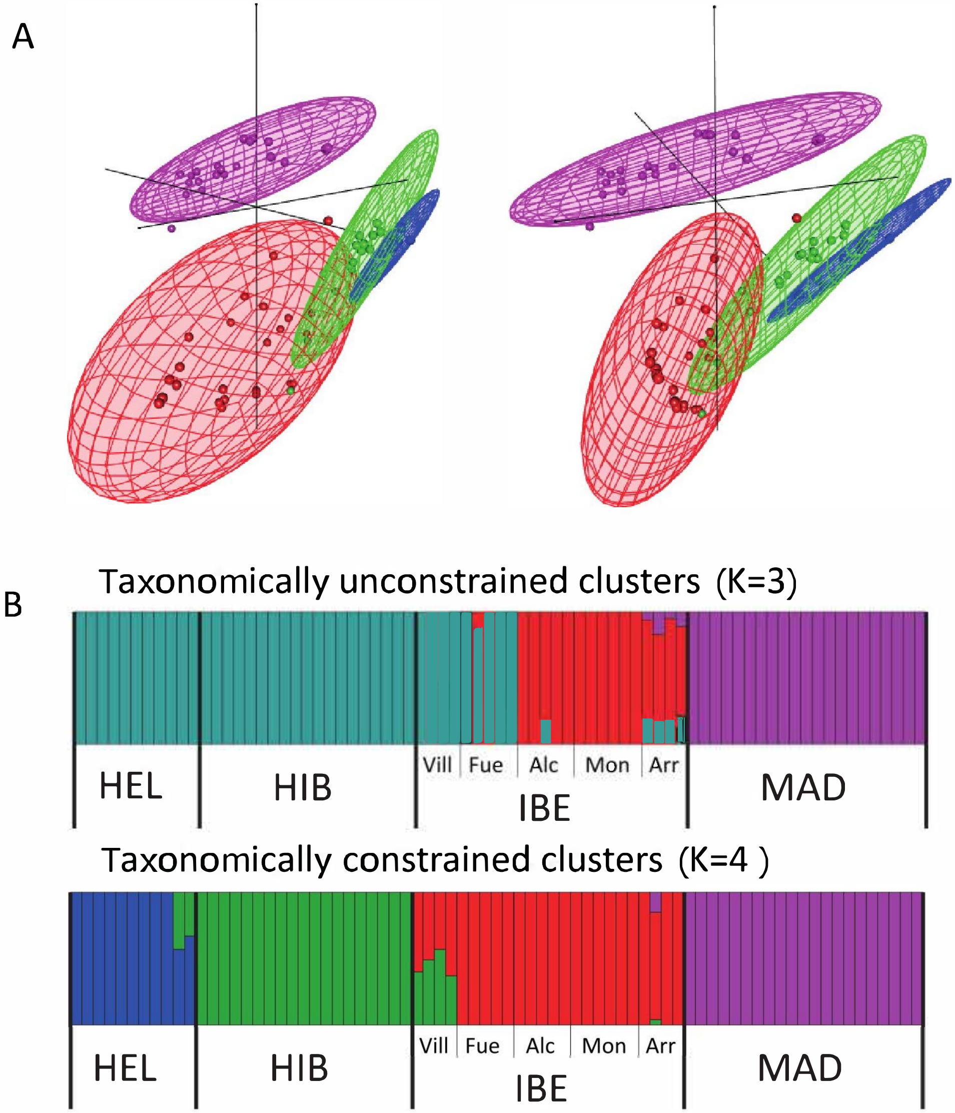

Supplementary figure 4. Differences in the principal components of the climatic PCA among the species of the western polyploid clade of *Hedera*: *H. hibernica* (HIB), *H. iberica* (IBE), and *H. maderensis* (MAD). The crossbar within the boxplot shows the median of the PCs, and the length of the box indicates the interquartile range. Shape of the violin plot reflects the kernel density plot. Pairwise comparisons are indicated with the correspondent p.value. Level of significance is shown. ****: $p \leq 0.0001$, ***: $p \leq 0.001$, **: $p \leq 0.01$, *: $p \leq 0.05$, ns: $p > 0.05$. (A) Principal Component 1, (B) Principal Component 2.

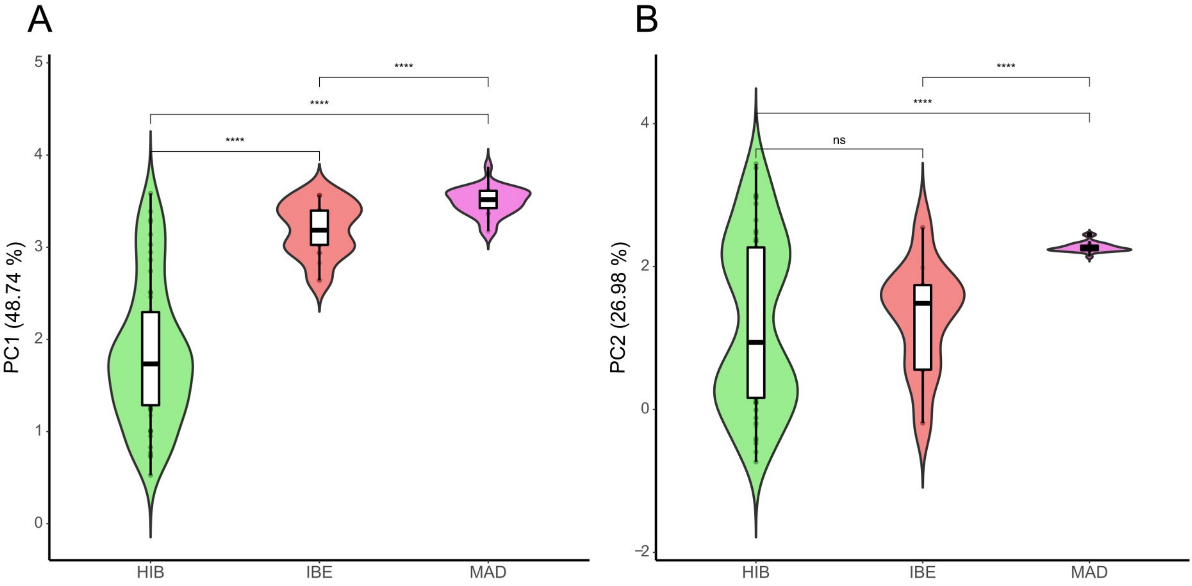

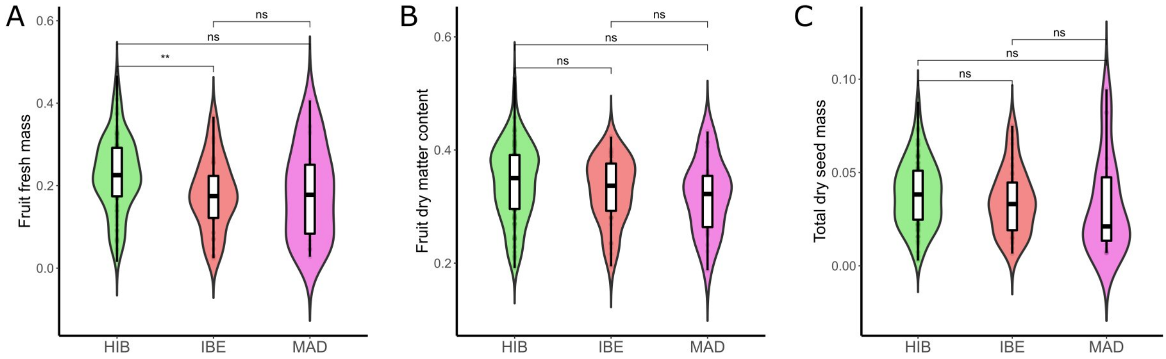


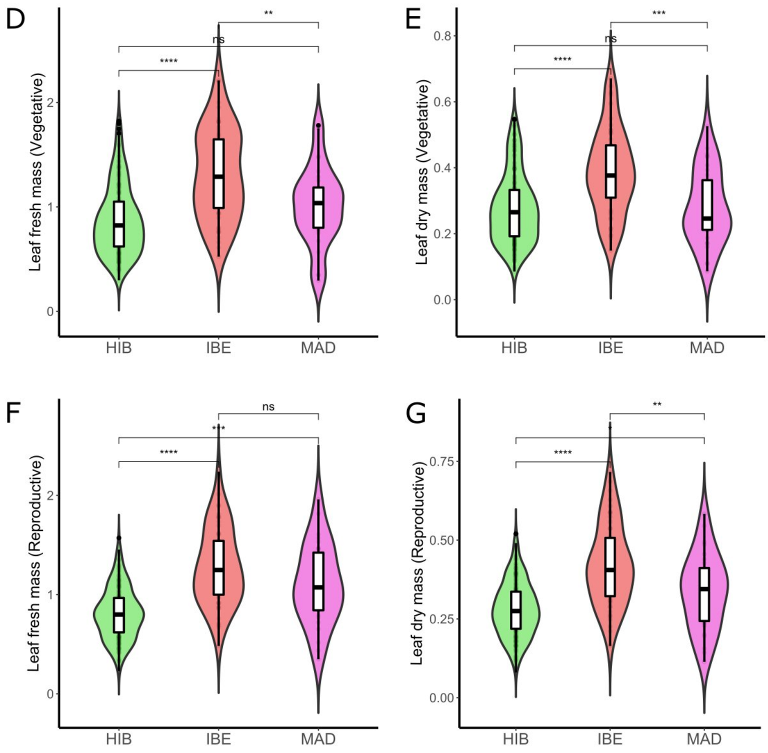


Supplementary figure 5. Vegetative and regenerative functional trait differences among the species of the western polyploid clade *Hedera*: *H. hibernica* (HIB), *H. iberica* (IBE), and *H. maderensis* (MAD). The crossbar within the boxplot shows the median, and the length of the box indicates the interquartile range. Shape of the violin plot reflects the kernel density plot of data. Pairwise comparisons are indicated with the correspondent p.value. Level of significance is shown. ****: $p \leq 0.0001$, ***: $p \leq 0.001$, **: $p \leq 0.01$, *: $p \leq 0.05$, ns: $p > 0.05$.
